# Benchmarking computational methods for multi-omics biomarker discovery in cancer

**DOI:** 10.64898/2025.12.18.695266

**Authors:** Athan Z. Li, Yuxuan Du, Yan Liu, Liang Chen, Ruishan Liu

## Abstract

Multi-omics profiling characterizes cancer biology and supports biomarker discovery for prognosis and therapy selection. Although numerous computational multi-omics biomarker identification methods have been proposed, their ability to identify clinically relevant biomarkers has not been systematically evaluated, leaving it unclear whether the resulting biomarker nominations are reliable for downstream validation. Here we systematically benchmark 20 representative statistical, machine learning and deep learning methods using curated gold-standard prognostic and therapeutic biomarkers across five real-world datasets. We evaluate performance in terms of both biomarker identification accuracy and stability. Overall, DeePathNet and Deep-KEGG achieve the best performance. Across methods, effective biomarker recovery is associated with the integration of biological knowledge, global feature interactions, multivariate feature attribution, and effective regularization. Analysis of omics type contributions reveals method- and modality-specific biases, highlighting the importance of broader omics integration. We further evaluate methods on simulated datasets to probe sensitivity with controlled signal and noise. By aggregating results from top-performing methods, we construct consensus biomarker panels that nominate candidates for potential investigations. Finally, we provide user-friendly interfaces to allow researchers to benchmark new methods against the 20 baselines or apply selected methods for biomarker identification on custom multi-omics datasets. Our benchmark is publicly available at https://github.com/athanzli/CancerMOBI-Bench.

## 1 Introduction

High-throughput technologies have enabled large-scale profiling of human cancers across genomic, transcriptomic, epigenomic, proteomic, and metabolomic layers [1, 2]. Collectively, these assays constitute a comprehensive and complementary view of tumor biology, substantially expanding the landscape for discovering biomarkers that can inform prognosis and therapy selection from a vast space of molecular candidates [3]. Nevertheless, high dimensionality, tumor heterogeneity, and complex cross-omics dependencies make robust and clinically meaningful biomarker discovery inherently challenging [4].

Over the past several years, a diverse array of computational methods has been developed to address this need (Supplementary Table 1). Statistical and machine learning (ML) methods extend classical frameworks such as matrix factorization, Canonical Correlation Analysis (CCA), or Uniform Manifold Approximation and Projection (UMAP) to extract shared patterns across omics and derive interpretable latent representations [10–12]. More recently, deep learning (DL) architectures, including feedforward neural networks (FNN), autoencoders, graph neural networks, and transformers, have been applied for multi-omics fusion by learning the nonlinear and hierarchical relationships across molecular layers [13–20]. These methods have achieved strong performance on a broad spectrum of cancer-related tasks, including prediction of cancer subtypes, metastatic status, pathological stages, survival outcomes, and therapeutic responses [13–15, 19–21]. Beyond predictions, many incorporate interpretability modules to identify biologically meaningful features, such as disease-associated subnetworks [22], dysregulated pathways [20], and most commonly, molecular biomarkers [12, 14–21, 23–26].

**Table 1.**
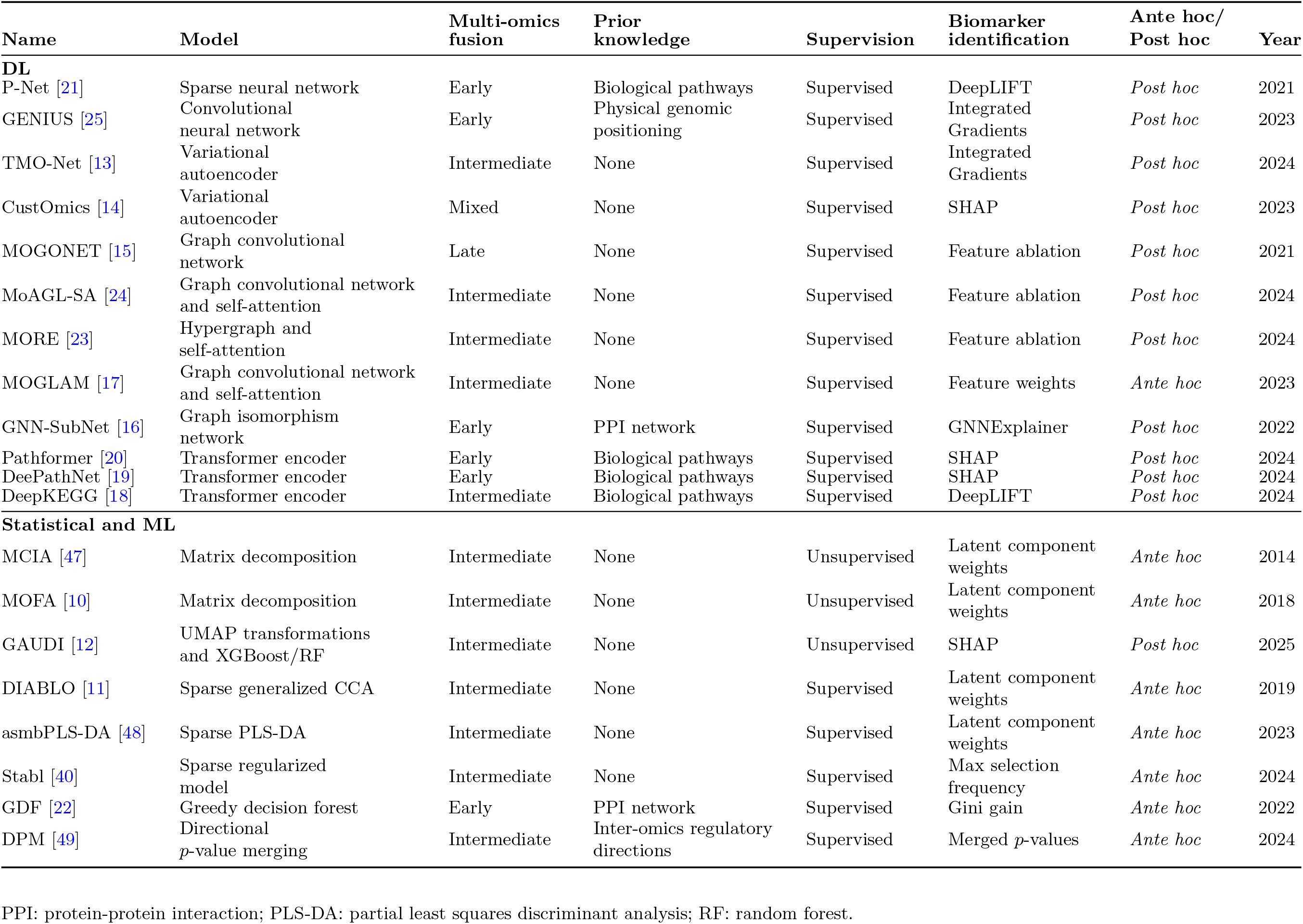
Overview of the computational multi-omics biomarker identification methods benchmarked in this study.

Despite the methodological progress, systematic evaluations of biomarker identification performance remain limited. Within individual method studies, evaluations of discovered biomarkers are typically proxy-based or qualitative, relying on functional enrichment analysis or literature support [12, 14–21, 23–26]. Such assessments are inferential and prone to selection bias, providing limited quantitative validation. Also, existing benchmark studies often rely on predictive performance as a proxy for biomarker identification performance [27–37]. However, high predictive accuracy does not necessarily imply reliable biomarker identification, as models can achieve strong prediction using features that are statistically discriminative but not biologically meaningful, particularly in high-dimensional, correlated multi-omics data [38]. Although some utilize synthetic datasets with predefined discriminative features for direct and quantitative assessment, these datasets are often simplistic and fail to match the complexity and heterogeneity of real-world cancer data [11, 31, 33, 39, 40]. Moreover, many recent multi-omics and DL methods remain absent from prior benchmarks [27, 29, 30, 34–37, 39, 41, 42].

Translational implications can arise from these gaps. As biomarker development proceeds from early discovery toward tests intended for specific clinical applications, successful translation requires analytical validity, clinical validity, and clinical utility [43, 44]. Because less than 1% of published cancer biomarkers enter clinical practice, more rigorous evaluation of discovery-stage methods is needed to better prioritize approaches with downstream translational potential [45].

To address these limitations, we provide a systematic benchmark of computational methods for multi-omics biomarker identification. A major obstacle has been the lack of task-specific biomarker sets for ground-truth evaluation. To address this, we collected Tier I biomarkers as defined by AMP/ASCO/CAP guidelines [8] from Clinical Interpretation of Variants in Cancer (CIViC) [5], MSK’s Precision Oncology Knowledge Base (OncoKB) [6], and Cancer Genome Interpreter (CGI) [7]. Using these gold-standard biomarkers, our benchmarking enables direct, quantitative, and clinically meaningful assessment of biomarker identification performance. Instead of relying on indirect evaluation through predictive performance, our approach directly quantifies each method’s ability to recover curated, clinically relevant biomarkers via rank-based metrics. We constructed five real-world task datasets from The Cancer Genome Atlas (TCGA) [46] through systematic filtering to achieve both computational feasibility and alignment between biomarkers and tasks.

In total, we benchmark 8 statistical and ML methods and 12 DL methods, covering a wide range of representative modeling and feature-identification approaches. Performance was evaluated across diverse experimental settings and metrics. Several methods exhibit strong ability to recover clinically validated biomarkers, frequently surfacing gold-standard biomarkers among the top ranks across experiments. Analysis of omics type contributions further characterizes biomarker- and method-specific biases. Transformer-based methods integrating biological pathways with advanced *post hoc* feature-attribution methods achieve the highest accuracy and strong stability, whereas univariate feature-ablation methods underperform on both dimensions. Simulation experiments show that statistical and ML methods are more sensitive to shifts in the discriminative signals of groundtruth features, but discrepancies with real data results underscore the indispensability of real-world cancer cohorts. Additionally, we derive consensus panels of multi-omics cancer biomarkers by aggregating the results of top-performing methods, providing candidates for potential downstream investigations. Our benchmark is publicly available with user-friendly evaluation pipelines that enable researchers to benchmark new methods against the 20 baselines on the curated task datasets, or to apply selected methods to other multi-omics data for biomarker identification.

## 2 Results

### 2.1 Benchmarking design and evaluation framework

We conducted a systematic benchmark of 20 computational methods for multi-omics biomarker identification, including 8 statistical and ML methods and 12 DL methods selected from literature (Supplementary Table 1) that encompass diverse methodological designs (Fig. 1c). Their model families, multi-omics fusion strategies, use of prior biological knowledge, and biomarker identification approaches are summarized in Table 1, with detailed descriptions provided in Supplementary Methods.

**Figure 1.**
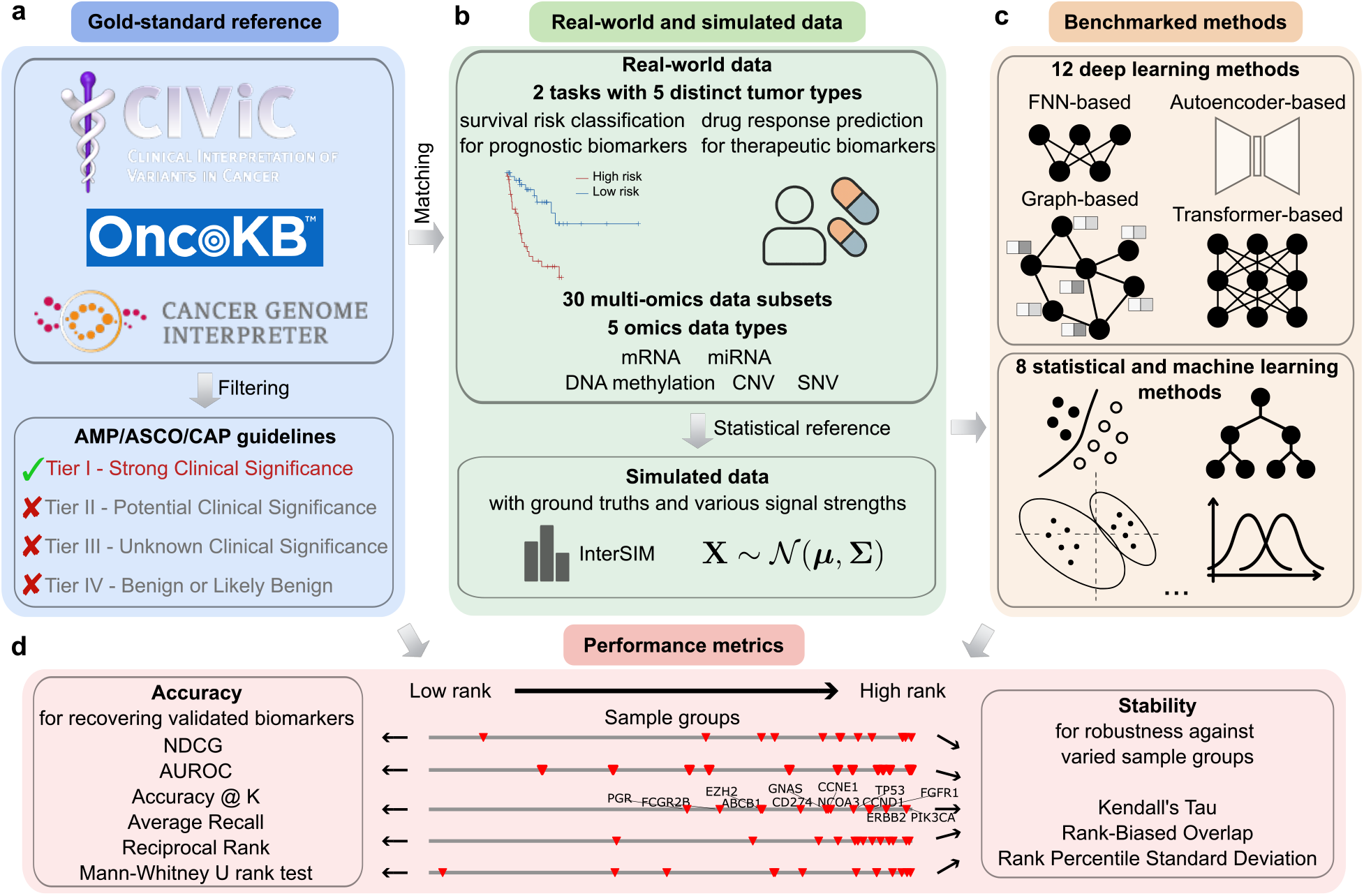
Overview of benchmarking workflow. **a**, Biomarkers were collected from three oncology knowledge bases, including CIViC [5], OncoKB [6], and CGI [7], and those with Tier I evidence determined by AMP/ASCO/CAP guidelines [8] were retained. **b**, Real-world data included 30 multi-omics data subsets encompassing five tasks and six omics combinations. Tumor types and drugs for prognostic and therapeutic tasks were matched to available biomarkers, and determined through a series of filtering rules (Supplementary Methods). Real-world data were used as statistical reference for simulated multi-omics data generation with InterSIM [9]. **c**, Twelve DL methods were benchmarked with different architectural backbones, including FNN, autoencoder, graph, and transformers. Eight statistical and ML methods were also benchmarked. **d**, Rank-based metrics for performance quantification. Six accuracy metrics and three stability metrics were used (Methods).

To anchor evaluation in clinical relevance, we curated Tier I cancer biomarkers, as defined by AMP/ASCO/CAP guidelines [8]. These biomarkers were harmonized across CIViC [5], OncoKB [6], and CGI [7] (Fig. 1a, Supplementary Tables 2, 3). After filtering, the curated biomarkers were mapped to five benchmark task datasets constructed from TCGA, including survival risk classification for Breast invasive carcinoma (BRCA), Lung adenocarcinoma (LUAD), and Colon and Rectum adenocarcinoma (COADREAD), and drug response prediction for Cisplatin with Bladder Urothelial Carcinoma (BLCA) and Temozolomide with Brain Lower Grade Glioma (LGG) (Fig. 1b, Supplementary Methods, Table 2). The curated biomarker set spans multiple clinically established alteration classes, including sequence variants, copy-number alterations, expression-based markers, DNA methylation markers, gene fusions, and limited protein-level markers (Supplementary Table 3), capturing major mechanisms with established clinical significance such as oncogenic driver activation, tumor suppressor inactivation, DNA-repair deficiency, epigenetic silencing, and immune-related expression states [5–8].

**Table 2.** Summary statistics of the real datasets used in this study.

| Task | Dataset | Sample size |  | #Features |  |  |  |  | #Biomarkers |
| --- | --- | --- | --- | --- | --- | --- | --- | --- | --- |
|  |  | Low risk | High risk | CNV | DNA<sub>m</sub> | mRNA | miRNA | SNV |  |
| Survival | BRCA | 324 | 323 | 18 645 | 46 859 | 19 504 | 1 581 | 15 118 | 14 |
|  | LUAD | 212 | 211 | 18 645 | 46 859 | 19 497 | 1 581 | 17 103 | 14 |
|  | COADREAD | 187 | 185 | 18 645 | 46 859 | 19 471 | 1 555 | 18 394 | 12 |
| Drug response |  | Response | Non-response |  |  |  |  |  |  |
|  | Cisplatin (BLCA) | 40 | 20 | 18 645 | 46 859 | 19 288 | 1 404 | 10 502 | 1 |
|  | Temozolomide (LGG) | 20 | 103 | 18 645 | 46 859 | 19 411 | 1 392 | 3 656 | 2 |

**Table 3.**
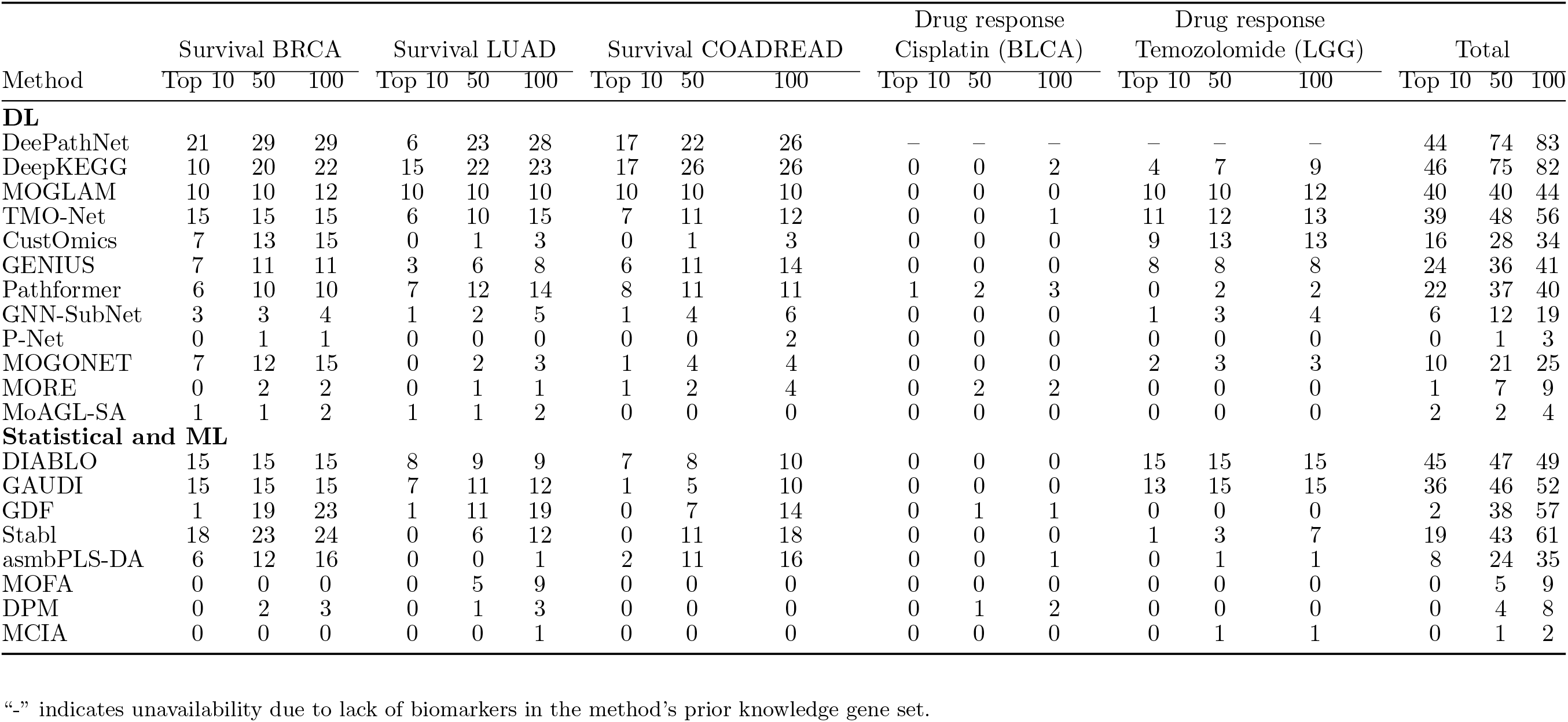
Frequency of surfacing biomarkers. The number of times a method ranks at least one gold-standard biomarker within the top 10, 50, or 100 out of all the 30 experiments, including six omics combinations with five-fold cross-validation.

Since the benchmarked methods employed different omics combinations in their original case studies (Supplementary Table 4), we defined a unified set of omics combinations to achieve comparability across methods. For each task, we combined messenger RNA (mRNA) expression with two additional omics types from microRNA (miRNA), DNA methylation (DNAm), Copy Number Variation (CNV), and Single Nucleotide Variation (SNV). This produced six distinct omics combinations per task, leading to 30 multi-omics subdatasets that were each evaluated with five-fold cross-validation (Fig. 1b). To complement real-world data, we also generated simulated multi-omics data using InterSIM [9]. These simulations contain predefined discriminative features serving as ground truth, enabling controlled assessment of method sensitivity across a range of signal levels (Fig. 1b, Supplementary Table 5, Supplementary Methods).

Biomarker identification performance was evaluated using nine complementary metrics capturing different aspects of accuracy and stability (Fig. 1d, Methods). For real-world cohorts where ground truth biomarkers are partially known, rank-based metrics were applied to reward the prioritization of validated biomarkers without penalizing unvalidated candidates. For simulated datasets with complete ground truth, evaluation adopted metrics such as AUROC and accuracy. Stability was quantified as the similarity of ranked gene lists across cross-validation, using metrics that reflect top-rank consistency, global ranking agreement, and biomarker ranking variability (Methods). For comparability, all method output was converted to gene-level ranking lists prior to metric calculation (Methods).

### 2.2 Accuracy on real-world data

Accurate identification of clinically relevant candidates from high-dimensional molecular space represents a hallmark of an effective biomarker discovery method. Therefore, we first examined the performance of each method for the recovery of clinically validated biomarkers in real-world cancer cohorts. Across five TCGA-derived tasks and six omics combinations, we compared methods using rank-based metrics that quantify different aspects of biomarker prioritization: Average Recall (AR) for overall prioritization, Reciprocal Rank (RR) for early retrieval of a single biomarker, and Normalized Discounted Cumulative Gain (NDCG) for early recovery of multiple biomarkers. These were complemented by a Mann-Whitney U test on global rankings and by success counts within the top 10, 50, and 100 ranks (Fig. 2a,b, Table 3, Methods). Detailed results of each omics combination are reported in Supplementary Figs. 1-5 and Supplementary Figs. 11-15.

**Figure 2.**
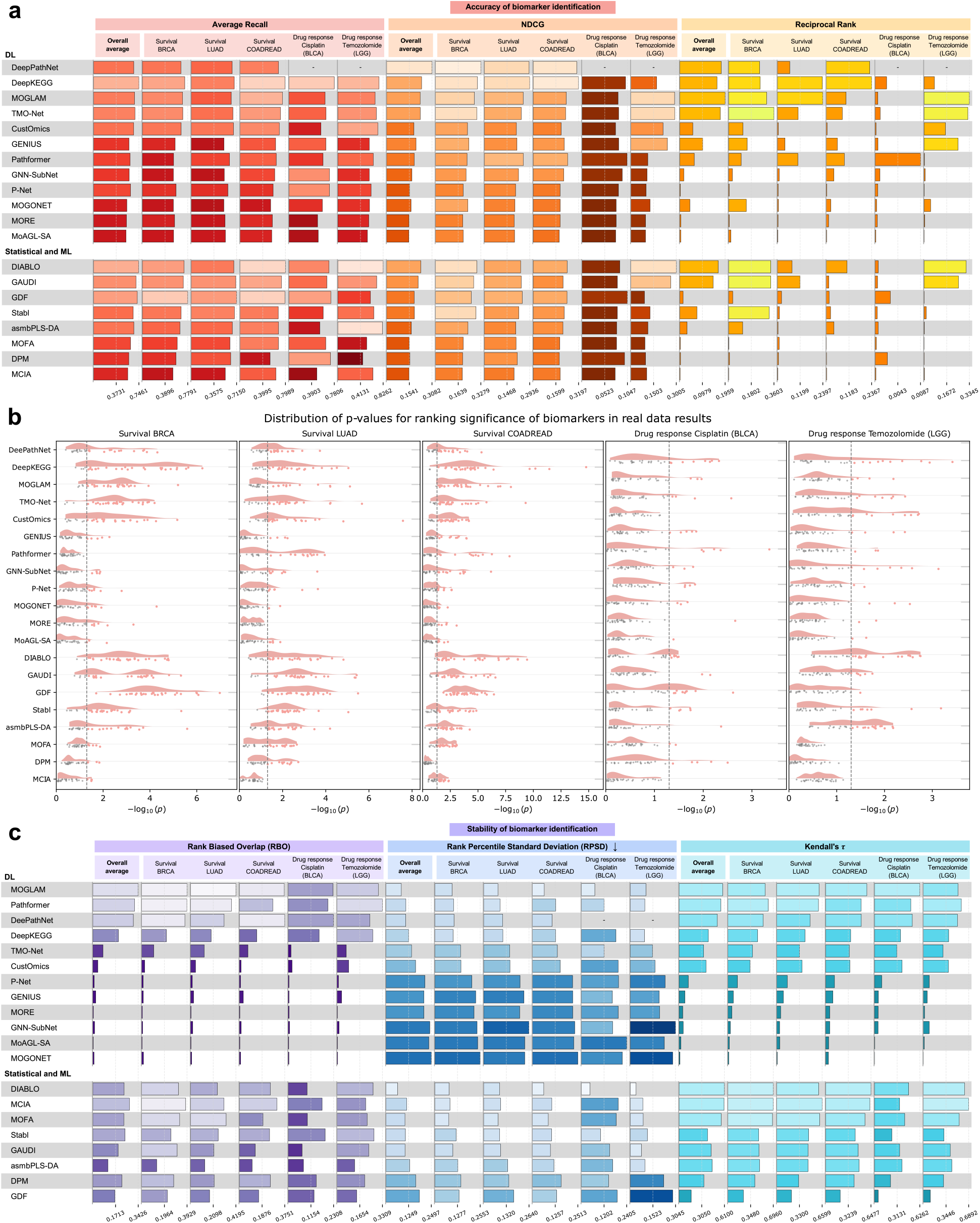
Overall performance on real-world data. **a**, Accuracy for all tasks and metrics. For a particular method and task, the height of the bar represents the averaged metric value of the cross-validation mean across all six omics combinations. **b**, Mann-Whitney U test *p*-values (negative log_10_-transformed). Each data point represents a single experiment corresponding to a specific omics combination and cross-validation fold. The vertical dashed line represents a *p*-value of 0.05. **c**, Stability for all tasks and metrics. For a particular method and task, the height of the bar represents the averaged metric value across all six omics combinations. NDCG: Normalized Discounted8Cumulative Gain. “-” indicates unavailability due to lack of biomarkers in the method’s prior knowledge gene set.

We identified a group of relatively accurate methods, including DeePathNet, DeepKEGG, MOGLAM, TMO-Net, DIABLO, and GAUDI, which frequently recovered validated biomarkers among the top ranks, with a top-10 success rate of around 26.7% (about 40 out of 150 experiments; Table 3) as well as comparable overall accuracy (Fig. 2a,b). Within this group, the transformer-based, pathway-informed methods DeePathNet and DeepKEGG were top performers. In survival prediction tasks, DeePathNet and DeepKEGG recovered a validated biomarker within the top 10 in around half of all experiments (44/90 and 42/90, respectively) and achieved the highest NDCG scores, indicating strong ability to surface multiple biomarkers simultaneously. Expanding the ranking threshold from top-10 to top-100 further accentuates this advantage, increasing the success rate of DeePathNet and DeepKEGG to around 50% across all tasks and 90% in survival tasks, whereas MOGLAM, TMO-Net, DIABLO, and GAUDI showed minimal improvement (Table 3). This divergence aligns with the omics contribution analysis (Fig. 3), suggesting that DeePathNet and DeepKEGG exploit a broader spectrum of omics combinations to retrieve relevant signals.

**Figure 3.**
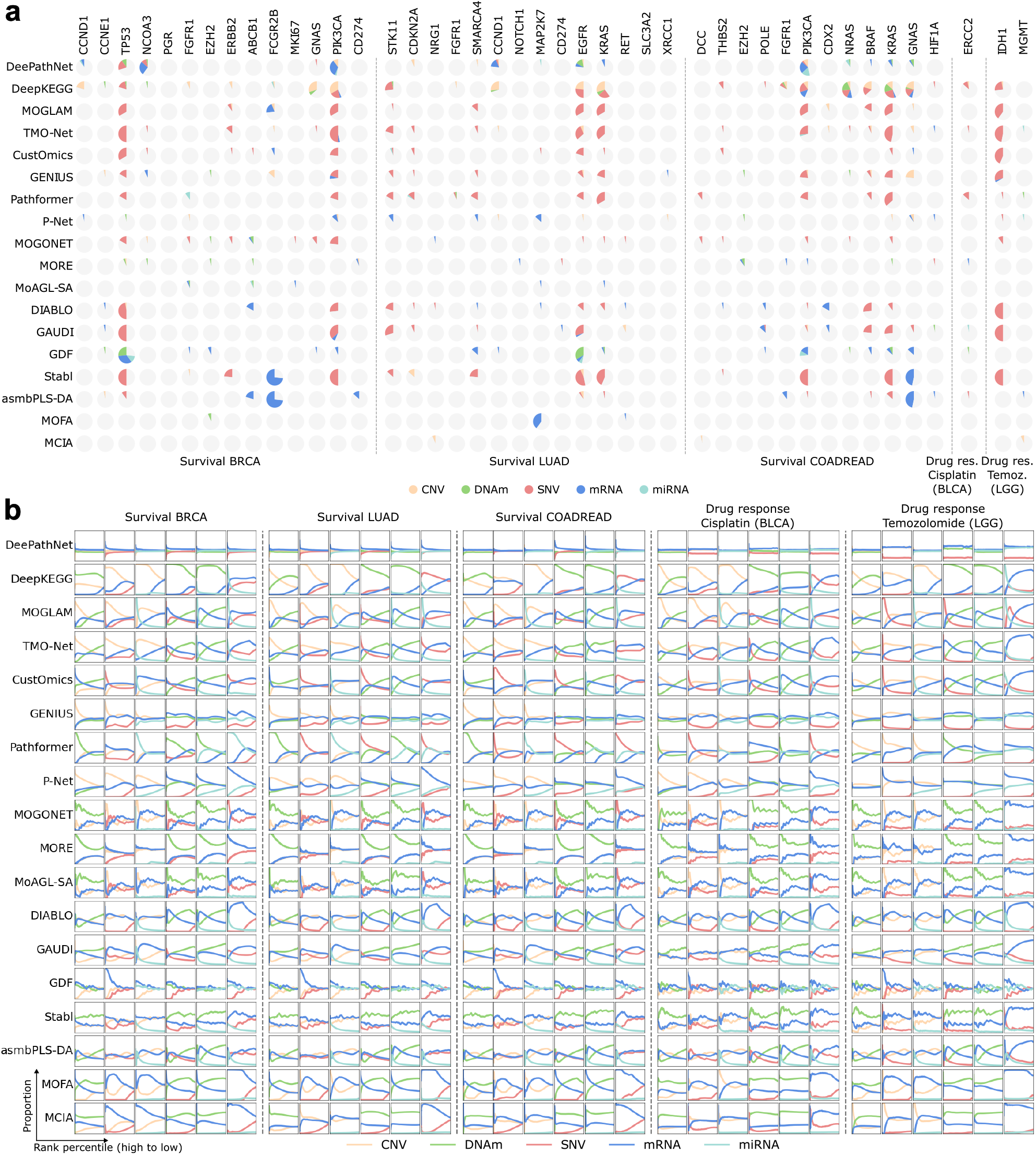
Omics type contributions in biomarker identification. **a**, Pie charts showing the omics types driving the identification of a biomarker. Each pie has 30 portions, with each representing a single experiment with a specific cross-validation fold (five in total) and omics combination (six in total). A portion is colored by the dominant omics type when the biomarker is ranked in the top 1% (Methods), and left gray otherwise. **b**, Omics type proportions (y-axis) from high (left) to low (right) ranks (x-axis). Each task contains six columns corresponding to the six omics combinations. Proportions were calculated at each of the 1000 rank percentile cutoffs. The ranks were based on the raw feature scores before gene-level conversion (Methods), which were concatenated from the five score lists output by cross-validation.

At the lower end of performance, methods relying on feature ablation (MOGONET, MoAGLSA, MORE) rarely recovered biomarkers across both success counts and accuracy metrics (Table 3, Fig. 2a, b). Their underperformance likely reflects the complementarity among correlated multiomics features, where ablation of a single feature is compensated by others [38]. Among statistical and ML methods, MOFA, DPM, and MCIA consistently ranked lowest across tasks and metrics. DPM is intrinsically univariate, while MOFA and MCIA rank features via unsupervised matrix decomposition that may not align with clinical endpoints (Supplementary Methods). In contrast, GAUDI, despite also being unsupervised, achieved substantially better performance by combining tree-based models with SHAP-based attribution, highlighting the benefit of multivariate attribution even in unsupervised settings (Fig. 2a, b).

Deep neural networks are not intrinsically superior to conventional statistical and ML models. Although DL methods exhibit a higher performance ceiling, with DeePathNet and DeepKEGG achieving the highest accuracy and maintaining it across multiple cancers and omics combinations (Supplementary Figs. 1-5), well-designed statistical methods such as DIABLO and GAUDI achieved competitive accuracy. Conversely, Pathformer achieved only moderate accuracy despite using a transformer backbone, and MOGLAM was the strongest graph-based method despite most graph methods ranking low. These contrasts indicate that specific methodological designs, rather than model family, primarily determine success. The factors underlying these differences are systematically analyzed in the following sections.

Task complexity also affects performance. Almost every method underperformed on the BLCA cisplatin drug-response prediction task, particularly on top-rank-biased metrics such as NDCG and RR, indicating the increased difficulty of surfacing ERCC2 relative to other targets (Fig. 2a, Supplementary Figs. 4, 14). This likely stems from two factors. Clinically, ERCC2’s predictive signal is specific to cisplatin-based neoadjuvant chemotherapy with pathologic downstaging endpoints, hence mixing metastatic settings, radiographic endpoints, or carboplatin treatments introduces label noise [50]. Biologically, only helicase-domain loss-of-function ERCC2 variants truly abrogate nucleotide-excision repair and confer cisplatin sensitivity, thus collapsing all ERCC2 mutations into a single feature further obscures the signal [51]. Nevertheless, Pathformer appeared as a notable exception, as it was the only method consistently identifying ERCC2 within the top ranks when using the mRNA+CNV+SNV omics combination (Supplementary Figs. 4, 14). Relative to other transformer-based methods, Pathformer employs a more expressive criss-cross attention mechanism and architectural design (Supplementary Methods). This suggests that more sophisticated architectures can extract weak signals from noisy data under certain omics combinations, though such advantages may trade off against performance on less difficult tasks.

**Figure 4.**
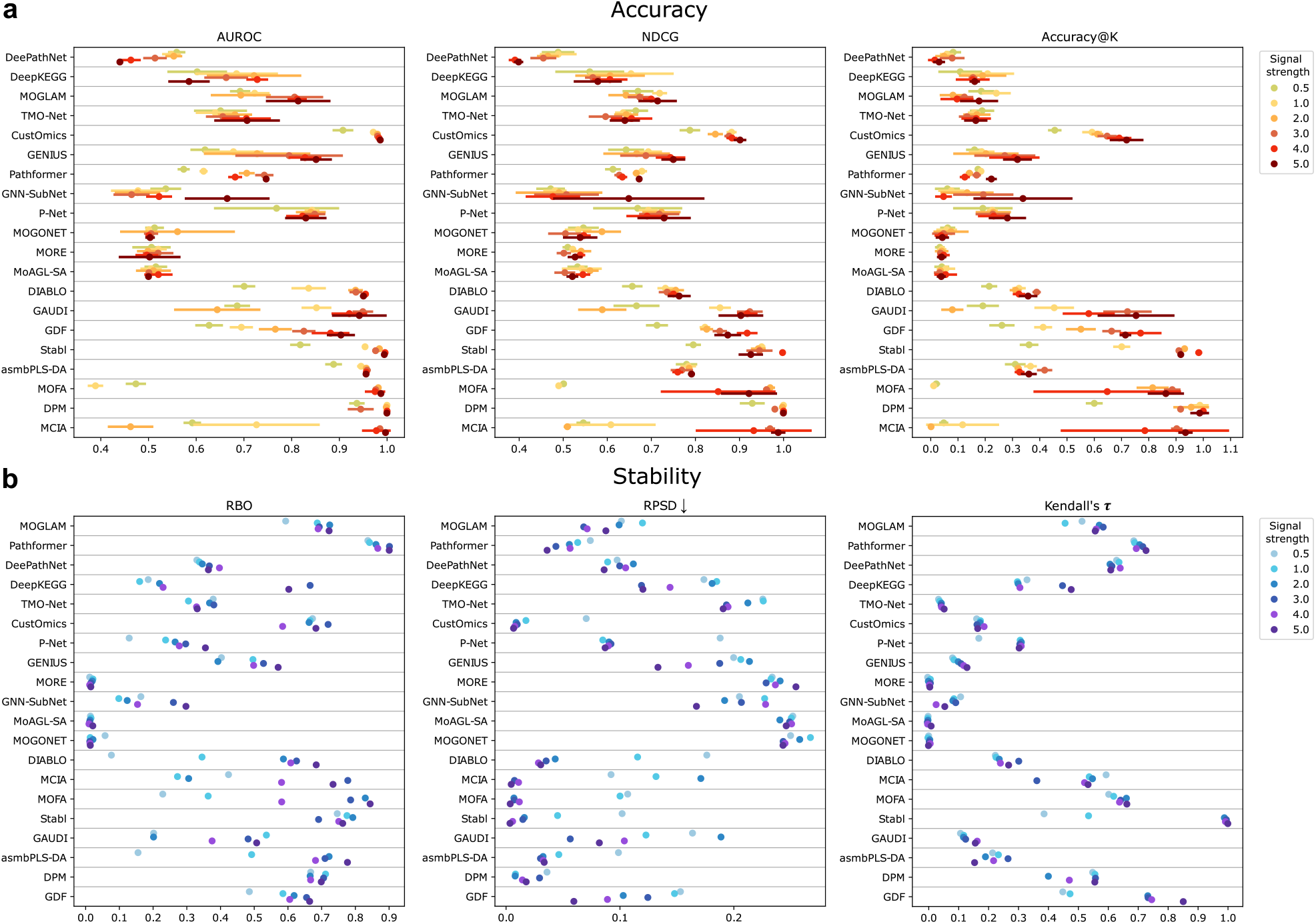
Performance on simulated data. Cleveland dot plots showing biomarker identification accuracy (**a**) (bars represent standard deviation across five-fold cross-validation) and stability (**b**). Colors represent a particular biomarker signal strength (Supplementary Methods).

### 2.3 Stability on real-world data

The reproducibility of biomarkers is essential for clinical translation, and high stability is a prerequisite for reproducible biomarker discovery across diverse cohorts. We quantified stability using the five cross-validation folds per omics combination per task, averaging the pairwise similarities between gene ranking lists (Methods). Rank-biased Overlap (RBO) assesses top-rank consistency, Rank Percentile Standard Deviation (RPSD) measures the dispersion of biomarker rankings across splits, and Kendall’s *τ* quantifies the overall consistency across full rankings (Fig. 2c, Methods). Stability results of each omics combination are reported in Supplementary Figs. 6-10.

In general, we found that statistical and ML methods were markedly more stable than DL methods, whereas the latter exhibited greater variability, encompassing both the most and least stable methods (Fig. 2c). A subset of DL methods, including MOGLAM, Pathformer, and DeePathNet, surpassed all statistical and ML methods, achieving both high top-rank stability (RBO) and strong global stability (low RPSD and high Kendall’s *τ*). By contrast, DIABLO, which is the most stable statistical and ML method, reached the best global stability but failed to attain comparable RBO, suggesting that although its rankings are internally coherent, they are less consistent at top than those produced by the most stable DL methods. At the opposite extreme, several DL methods, including P-Net, GENIUS, GNN-SubNet, MOGONET, MoAGL-SA, and MORE, were less stable than the least stable statistical and ML methods. Their RBO scores fall below that of asmbPLS-DA, the least stable statistical and ML baseline at the top ranks, and their global stability was lower than that of GDF, which had the highest RPSD and lowest Kendall’s *τ* among statistical and ML methods (Fig. 2c). These results reflect the sensitivity of deep neural network training, as small changes in the input samples can drive stochastic gradient-based optimization into distinct solution basins [52], producing substantially different top-ranked features identified by *post hoc* attribution methods.

Among the most stable DL methods, incorporating biological pathway structures (Pathformer, DeePathNet, DeepKEGG) and employing explicit regularization (MOGLAM) appeared to improve stability, while DL methods lacking such design choices tended to be among the least stable (Fig. 2c).

Jointly considering all methods, those that fail to capture global feature dependencies (GNN-SubNet, P-Net, GENIUS) were generally less stable and also underperformed in accuracy, suggesting that insufficient modeling of long-range molecular interactions diminishes the stabilizing contribution of global context [53] (Fig. 2c).

### 2.4 Determinants of biomarker identification performance

Joint analysis of accuracy and stability revealed substantial variation across the benchmarked methods. Autoencoder-based methods such as TMO-Net and CustOmics achieved strong accuracy but were unstable, Pathformer exhibited high stability but only moderate accuracy, and conventional statistical methods including MCIA and MOFA were inaccurate but highly stable. Nevertheless, high accuracy and stability are achievable simultaneously: MOGLAM, DeePathNet, DeepKEGG, and DIABLO were both accurate and stable, whereas P-Net, GNN-SubNet, MOGONET, MORE, MoAGL-SA, and DPM underperformed on both dimensions (Fig. 2). These contrasts indicate that the decisive factor is specific methodological design rather than model family. Across the diverse tasks and omics combinations evaluated, four methodological factors consistently distinguished high-from low-performing methods.

#### Feature attribution

The approach used to infer feature importance emerged as a primary performance determinant. Methods employing multivariate attribution, including SHAP (DeePathNet, Pathformer, CustOmics, GAUDI), DeepLIFT (DeepKEGG, P-Net), Integrated Gradients (TMO-Net, GENIUS), GNNExplainer (GNN-SubNet), and *ante-hoc* feature weighting (MOGLAM, DIABLO, asmbPLS-DA, Stabl), consistently outperformed those relying on univariate feature ablation (MOGONET, MoAGL-SA, MORE) or per-feature statistical tests (DPM) (Fig. 2a, b, Table 3). In high-dimensional multi-omics data, features are correlated both within and across molecular layers. Univariate approaches underestimate the importance of features whose effects are distributed across correlated molecular partners, because removing or testing a single feature can be compensated by its correlated counterparts [38]. Multivariate methods evaluate each feature’s contribution in the context of others, capturing the joint dependency structure essential for identifying clinically relevant biomarkers. The importance of this factor is also illustrated by GAUDI, which demonstrates that combining tree-based models with SHAP can compensate for the absence of supervised labels, outperforming several supervised DL methods that rely on weaker attribution approaches (Fig. 2a, b).

#### Integration of biological knowledge

Encoding biological pathway structures into model architectures was associated with high accuracy and stability. The two top-performing methods, DeePathNet and DeepKEGG, encode pathway knowledge into transformer backbones (Table 1, Supplementary Methods). Pathway-informed architectures constrain the hypothesis space to biologically plausible feature groups, reducing overfitting and directing the model toward functionally coherent molecular associations. However, knowledge encoding alone is insufficient. P-Net also incorporates pathways but underperforms due to sparse connectivity that suppresses inter-pathway communication, and GNN-SubNet uses PPI networks but is limited by local message-passing (Supplementary Methods). These contrasts indicate that the effectiveness of prior knowledge depends on the architecture’s capacity to propagate learned representations across the encoded biological structures.

#### Scope of feature interactions

Methods modeling long-range dependencies across the feature space exhibited markedly higher accuracy and stability than those restricted to local interactions. Transformer self-attention enables pathways to attend to one another, capturing cross-pathway dependencies that reflect the interconnected nature of biological processes. In contrast, CNNs (GENIUS) are constrained by local receptive fields, sparse networks (P-Net) suppress inter-node communication, and graph message-passing (GNN-SubNet) attenuates with distance [53]. Among statistical methods, DIABLO’s generalized CCA models global cross-omics covariance, contributing to its competitive performance (Supplementary Methods). These observations indicate that when biomarker-associated features span multiple interacting molecular layers, architectures restricted to local interactions cannot adequately capture the cross-layer dependencies required for accurate and stable biomarker identification (Fig. 2).

#### Regularization

Regularization improved stability without compromising accuracy. MOGLAM learns graphs adaptively from the data and applies regularization on its feature-indicator matrix, yielding the highest stability among methods (Fig. 2c, Supplementary Figs. 6-10). DL methods without explicit regularization (P-Net, GENIUS, GNN-SubNet, MOGONET, MoAGL-SA, MORE) exhibited low stability, reflecting the sensitivity of stochastic optimization to training set perturbations [52]. The co-occurrence of high accuracy and stability in methods such as MOGLAM, DIABLO and Stabl demonstrates that these objectives are simultaneously achievable through appropriate regularization (Fig. 2).

These four factors interact synergistically. DeePathNet and DeepKEGG combine all four (pathway-informed transformer architectures with global self-attention, multivariate attribution, and implicit regularization via structured pathway encodings), achieving the highest overall performance (Fig. 2, Table 3). Methods lacking multiple factors (e.g., MOGONET, MoAGL-SA, MORE with local graph interactions, univariate ablation, and no prior knowledge) consistently show weak performance. This indicates that strong biomarker identification is not driven by a single design choice, but instead emerges from the combination of multiple favorable design principles.

### 2.5 Contribution of omics types to biomarker identification

Different molecular layers characterize complementary aspects of tumor biology, but their relative contributions to biomarker discovery remain unclear. Although most methods evaluated one or two omics combinations in their original case studies (Supplementary Table 4), a systematic comparison across a broader set of omics combinations is essential for uncovering the underlying biological mechanisms and informing both method development and experimental design. Consistent with this need, we observed substantial differences in performance across omics combinations (Supplementary Figs. 1–15). In many cases, the omics combinations chosen in the original studies did not yield superior performance compared to untested alternatives, supporting our inclusion of an extended set of omics combinations. We also found that for almost all methods, high accuracy was confined to a specific omics combination, with DeePathNet and DeepKEGG being notable exceptions. In survival tasks, DeePathNet and DeepKEGG identified biomarkers within the top 10 ranks across nearly all omics combinations (RR ≥ 0.1; Supplementary Figs. 1c, 2c, 3c). To further dissect these results, we analyzed omics type contributions at both the biomarker and method level.

To pinpoint which omics types drive biomarker discovery, we examined the contributions of all five omics types for each identified biomarker (Fig. 3a). Here we define a biomarker as “identified” when it appeared within the top 1% of the gene-level ranking list (Methods). For each such case, we recorded the omics type with the highest feature score (Methods) as dominant. Figure 3a provides a comprehensive summary for all experiments. Detailed results of each omics combination are reported in Supplementary Figs. 16-21.

We found that most identified biomarkers were driven by one predominant omics layer. For instance, for the temozolomide drug response prediction task within lower-grade glioma (LGG), the top-performing experiments consistently required SNV data. IDH1 was typically ranked at the top owing to SNV (Fig. 3a, Supplementary Figs. 17, 19, 21), consistent with the causal IDH1 R132 mutation and its co-occurrence with MGMT promoter methylation, the canonical predictor of temozolomide benefit [54]. In breast cancer (BRCA) survival, TP53 was almost always recovered through SNV (Fig. 3a, Supplementary Figs. 17, 19, 21), reflecting the prevalence of missense mutations in the DNA-binding domain that stratify poor outcomes, which is especially common in aggressive basal-like/TNBC [55, 56]. By contrast, FCGR2B was primarily identified through mRNA expression, particularly by Stabl and asmbPLS-DA (Fig. 3a, Supplementary Figs. 16-21), where transcriptomic signatures reflect immune activation rather than direct tumor genetics in breast cancer prognosis [57]. These examples illustrate how the dominant layer responsible for identification often reflects underlying tumor biology.

Nevertheless, several biomarkers were occasionally identified through non-canonical omics layers. For example, IDH1 was identified by DeepKEGG through CNV (Supplementary Fig. 16), TP53 by DeePathNet and DeepKEGG through DNAm (Supplementary Figs. 16 and 19), and FCGR2B by DeepKEGG, MOGLAM, and TMO-Net through CNV or miRNA (Supplementary Figs. 16, 18, 21). These anomalous identifications underscore the complex interplay across omics modalities and the potential for cross-omics compensation.

Since methods can identify biomarkers through omics layers that do not commonly drive the underlying biology, we next interrogated whether methods have intrinsic preferences for specific omics types. We summarized the proportions of each omics type at different rank percentile cutoffs using each method’s raw feature scores or rankings, per omics combination and task (Fig. 3b). Some methods demonstrated clear biases. As an example, GDF relied almost exclusively on mRNA, DNAm, and miRNA, whereas CNV and SNV rarely appeared as dominant (Fig. 3). Consequently, whereas most methods identified TP53 primarily via SNV, GDF identified TP53 through mRNA, DNAm, and miRNA (Fig. 3a, Supplementary Figs. 16-21). Other methods exhibited more diverse patterns. Notably, DeePathNet, DeepKEGG, P-Net, and GAUDI were the only four methods for which all five omics types appeared as dominant contributors across biomarkers, and three of these (DeePathNet, DeepKEGG, and GAUDI) were among the top performers. DeePathNet and DeepKEGG, in particular, displayed especially diverse contributions. For a single biomarker, as many as four different omics types could be dominant (e.g., PIK3CA with DeePathNet; NRAS, KRAS, and GNAS with DeepKEGG in the COADREAD survival task). By contrast, most other methods typically have a single dominant omics type per biomarker (Fig. 3a).

As it may be challenging to know *a priori* which omics combination will perform best, these findings indicate that including a broader set of omics types is beneficial for biomarker discovery from multi-omics data. Broad inclusion increases the likelihood of incorporating the mechanistic information relevant to specific biomarkers and allows methods to perform optimally, whereas restricting experiments to a single omics combination may overlook biomarkers whose key information resides in an untested molecular layer.

### 2.6 Performance on simulated data

A major challenge in benchmarking biomarker identification methods is the absence of complete ground truths. Our use of curated cancer biomarker knowledge bases, task matching, and rank-based metrics offers a practical solution to this problem for real-world data. In parallel, as a complement to real-world evaluation, we generated simulated multi-omics data with predefined discriminative features, enabling evaluation with metrics such as AUROC and accuracy. We varied the mean-shift of ground-truth features to modulate their discriminative strength (Supplementary Methods). It was observed that simulations reproduced some trends from real-data results. Feature-ablation DL methods (MOGONET, MoAGL-SA, MORE), which were weak on real cohorts, also underperformed in simulations. DIABLO remained among the top performers (Fig. 4), consistent with its capacity to model coherent multivariate structure. These consistencies increase confidence in methods whose performance appears robust to changes in data generation complexity.

However, discrepancies were more common. DeePathNet, DeepKEGG, MOGLAM, and TMO-Net became less competitive, and many statistical and ML methods (e.g., MCIA, MOFA, DPM) improved substantially. Within either the DL or the statistical and ML group, accuracy rankings differed from those within real cohorts (Fig. 4a). These shifts indicate that simplified generative mechanisms introduce a bias toward methods optimized for less noisy discriminative effects, while attenuating the relative advantages of methods intended to recover more complex relationships.

Across discriminative strengths, statistical and ML methods exhibited greater variability than DL methods (Fig. 4), indicating higher sensitivity to the magnitude of predefined effects. This is consistent with the fact that simulated datasets lack the heterogeneity and cross-omics dependencies observed in real tumors, thus simulations tend to favor conventional statistical and ML methods that assume simpler and less noisy underlying structure. In contrast, the capacity of DL models to utilize subtle or composite multivariate patterns becomes evident primarily in real-world datasets characterized by substantial heterogeneity and nonlinear cross-omics relationships. These discrepancies underscore the limitations of relying on synthetic discriminative features as benchmarking ground truth. Although widely used in prior work [11, 31, 33, 39, 40], simulation-based evaluations did not align with assessments based on real-world cohorts and clinically established biomarkers.

Stability patterns on simulated data were more consistent with those from real cohorts than accuracy (Fig. 4b, Fig. 2c), suggesting that stability is driven more by methodological design than by the numerical properties of data. Nonetheless, statistical and ML methods showed larger stability shifts as discriminative strength varied, indicating higher sensitivity to the structures within simulated data.

### 2.7 Consensus multi-omics biomarker panels

Our unified benchmarking framework, which evaluates multiple methods across shared tasks and omics combinations, provides a basis for deriving consensus biomarker panels. Therefore, we prioritized candidate biomarkers within each omics type by aggregating results from multiple top-performing methods. Since design choices and omics-specific dependencies vary widely across methods, the resulting rankings show only modest concordance among related approaches and weak concordance across distinct methodological families (Supplementary Fig. 22). Integrating these rankings can therefore uncover robust biomarker candidates that are less susceptible to method-specific biases and more representative of integrative molecular signatures, producing an accurate and reproducible consensus.

For each benchmarked task, we selected top-performing method-omics combination pairs, and derived consensus rankings via Robust Rank Aggregation (RRA) [58] (Methods). RRA converts cross-method agreement into per-feature *p*-values, accommodates partial rankings, and alleviates outliers and method-specific biases. The consensus provides statistically robust significance levels with controlled false discovery rates (FDRs), enabling distillation of sparse, accurate, and reproducible biomarker candidate panels (Table 4). As an example, in BRCA survival, the consensus identified 10 mRNA biomarkers (FDR *<* 10^−3^), 10 miRNA biomarkers (FDR *<* 10^−3^), 14 DNA methylation (DNAm) biomarkers (FDR *<* 10^−3^), 14 CNV biomarkers (FDR *<* 10^−4^), and 8 SNV biomarkers (FDR *<* 10^−6^) (Table 4). Derived from diverse methods and omics combinations that accurately and consistently recover clinically validated biomarkers, these panels provide high-confidence candidates for biological and clinical investigation. Full consensus rankings for all tasks and multi-omics features are available in Supplementary Data 1.

**Table 4.**
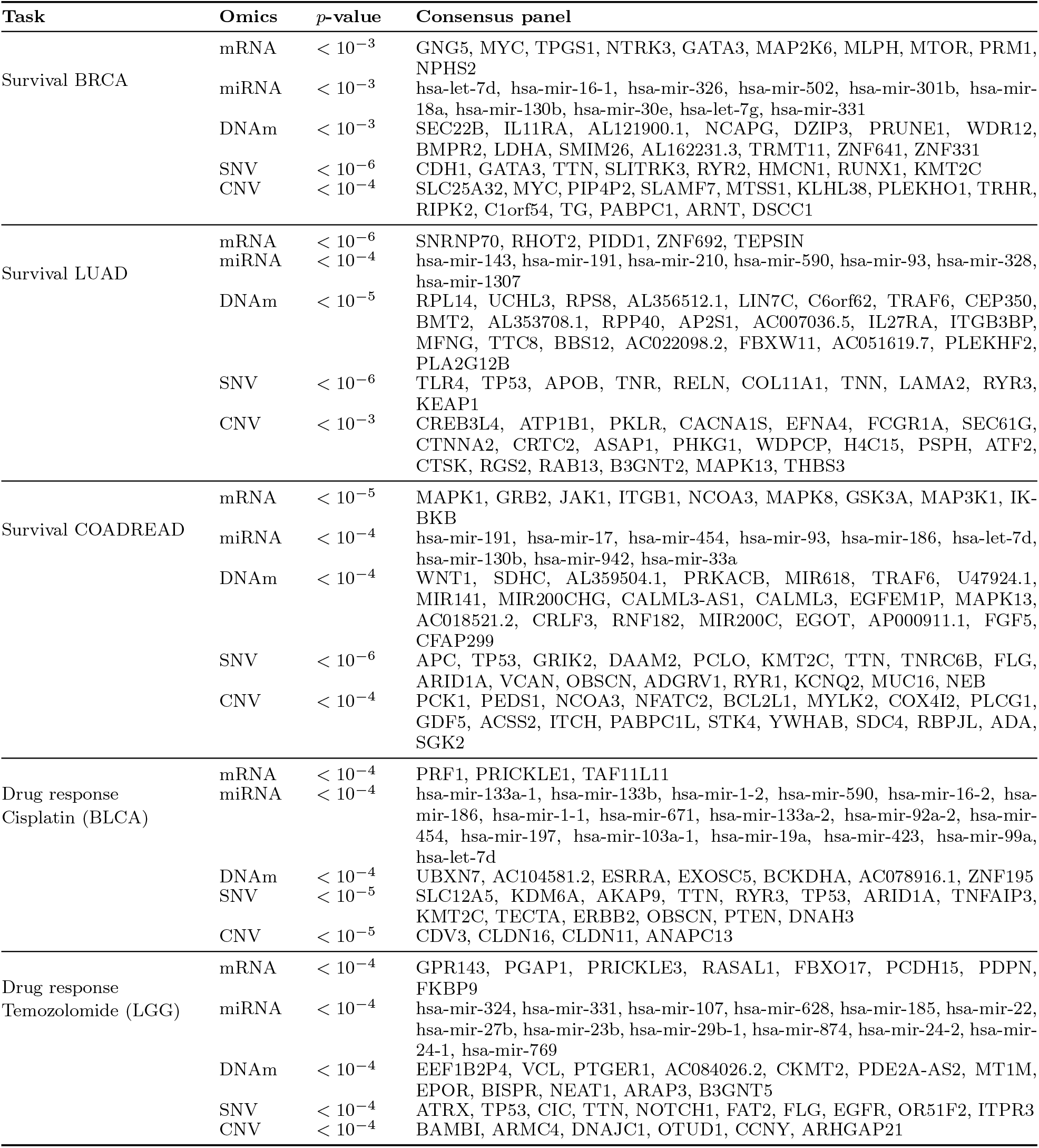
Consensus multi-omics biomarker panels by task and omics type. Benjamini-Hochberg corrected *p*-values from RRA [58] (Methods) are shown. Biomarkers already present in the gold-standard sets are omitted.

Many consensus panel candidates are established cancer genes, including CDH1 and GATA3 in the BRCA survival SNV panel [55], APC in the COADREAD panel [59], KDM6A in the cisplatin (BLCA) panel [60], and ATRX in the temozolomide (LGG) panel [61]. The recovery of such well-characterized genes provides evidence that the top-performing methods and RRA aggregation identify biologically meaningful candidates. All candidates are defined on routinely profiled molecular readouts and can be tested in independent cohorts using standard assays. However, they represent discovery-stage, computationally prioritized candidates that would require independent analytical and clinical validation before translational use [43].

### 2.8 Practical guidance for method selection

To facilitate practical use of the benchmark, we provide a method selection guide organized around three considerations: whether sample-level labels are available, whether GPU resources are accessible, and whether preservation of the original feature space is desired (Fig. 5). DeePathNet and DeepKEGG are the strongest general-purpose choices, achieving the highest accuracy with strong stability via pathway-informed transformer architectures with advanced attribution methods. When labels are unavailable, GAUDI is preferred; when GPU resources are unavailable, DIABLO offers strong performance with lightweight computation; when preservation of the original feature space is desired, MOGLAM is recommended. Methodological designs that are consistently unsuitable, including univariate feature ablation, univariate statistical scoring, and unsupervised matrix decomposition without multivariate attribution, are not recommended.

**Figure 5.**
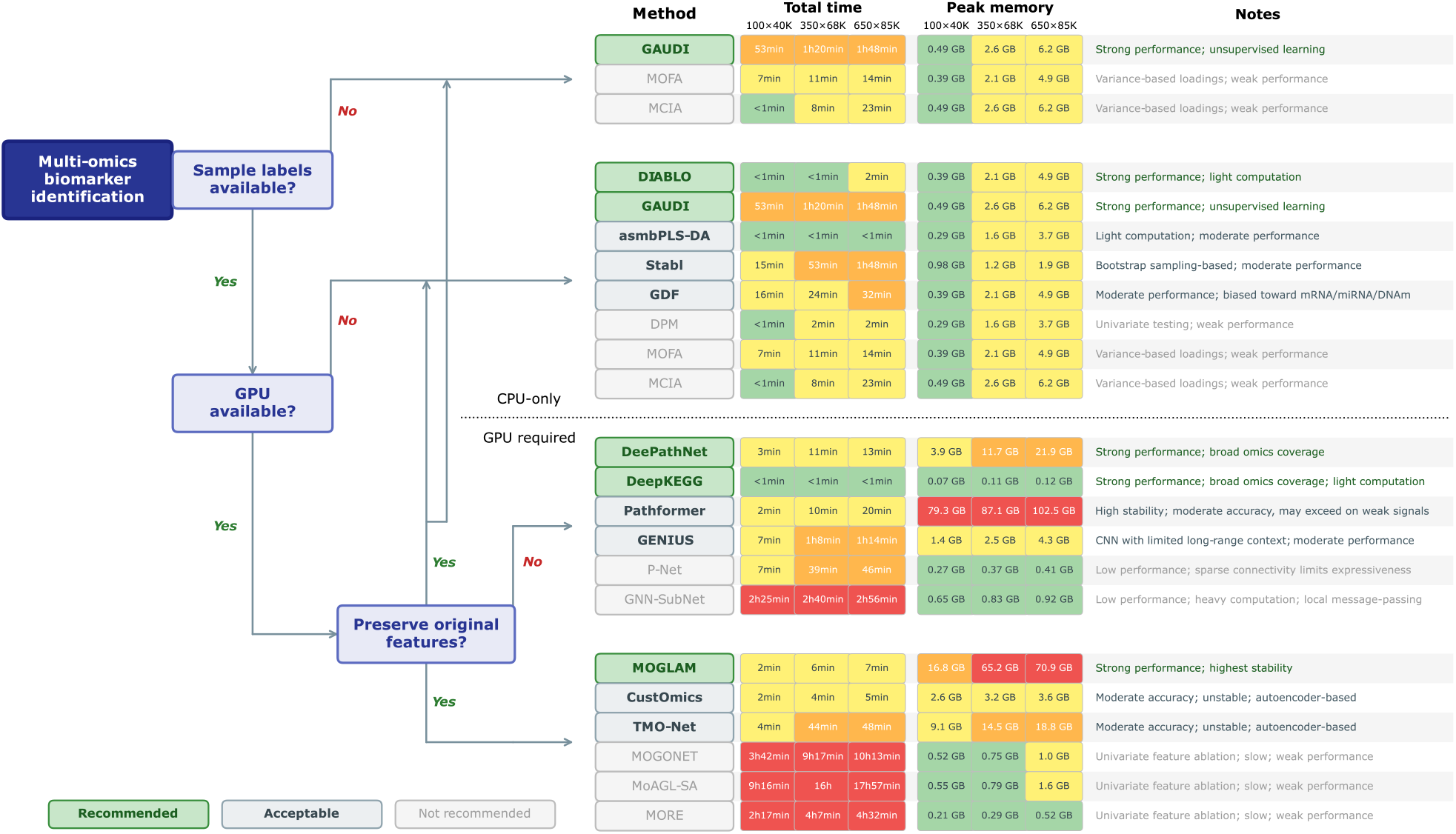
Practical guidance for method selection. Decision tree for selecting multi-omics biomarker identification methods based on sample-label availability, GPU accessibility, and whether preservation of the original feature space is desired. Methods are grouped into recommended, acceptable, and not recommended tiers according to overall benchmark performance. Representative runtime and peak memory across three data scales are provided for practical reference. GPU peak memory is provided for DL methods, and CPU peak memory is provided for non-DL methods.

Method selection does not depend on the specific omics types available, as all benchmarked methods support arbitrary omics combinations in our evaluation pipeline. We recommend including mRNA with two additional omics types to conform with our benchmark settings. For more robust biomarker prioritization, aggregating rankings from selected methods via RRA [58] can reduce method-specific biases.

## 3 Discussion

In this study, we presented a systematic benchmark for computational biomarker identification from multi-omics data, evaluated against clinically validated reference biomarkers. The results uncovered certain methodological designs that generally exhibited strong or weak accuracy and stability in biomarker identification, providing guiding principles for future method development and practical recommendations for method selection across different research scenarios. Analysis of omics type contributions indicates that biomarker identification is strongly influenced by the molecular modalities included and by model-specific biases, underscoring the importance of utilizing diverse omics layers for biomarker discovery. Results on simulated data showed that simplified generative assumptions can produce baseline results differing substantially from those observed in real-world cohorts, emphasizing the indispensability of real-world evaluations when assessing translational applicability. The consensus biomarker panels derived from the most accurate and stable methods and omics combinations provide computationally reproducible candidates that may inform potential biological and clinical investigations. Finally, the released evaluation framework enables researchers to benchmark new methods against the 20 baselines, or to apply selected methods for biomarker identification on other multi-omics data.

Challenges are identified that warrant future work. First, the incompleteness of available ground truth remains a fundamental limitation for real-world benchmarking. Although we compiled biomarkers from multiple curated cancer knowledge bases [5–7] and used rank-based metrics to mitigate the effects of missing annotations, biomarkers that have yet to be discovered or clinically characterized are naturally absent from evaluation. This is particularly relevant for therapeutic tasks, where the number of clinically validated biomarkers is smaller, which may lead to underestimation when models identify biologically relevant but undocumented candidates. As the coverage of cancer biomarker resources expands, future benchmarks will be able to incorporate broader reference sets. In addition, while our study focused on Tier I biomarkers [8] to maintain translational relevance, the inclusion of lower tiers in future work may help evaluate each method’s capacity to detect plausible but currently unvalidated candidates.

Another limitation is that all benchmark tasks were derived from TCGA and therefore do not constitute external validation on independent non-TCGA cohorts. Such datasets remain scarce when requiring matched mRNA, miRNA, DNA methylation, CNV, and SNV profiles together with biomarker-aligned clinical annotations and sufficient sample size. Nevertheless, TCGA includes substantial cross-center heterogeneity, as samples within each task originate from multiple tissue source sites, and their compositions vary across both tasks and cross-validation folds (Supplementary Fig. 24). This provides a partial internal test of robustness, but dedicated benchmarking on more independent cohorts will be a valuable direction for future work.

Our conclusions should also be interpreted within the scope of gene-centered multi-omics biomarkers. The curated biomarker set spans multiple clinically established alteration classes, covering major mechanisms with established clinical significance (Supplementary Table 3). However, many therapeutic interventions and drug responses are mediated through protein abundance, post-translational regulation, and signal transduction pathways, and the associated biomarkers remain underrepresented in both current oncology knowledge bases and cancer cohorts. Incorporating such modalities with matched clinical reference standards will be a potential direction for extensions.

## 4 Methods

### 4.1 Overview of benchmarked methods

Here we give an overview of the benchmarked methods. A detailed description of each is further provided in Supplementary Methods.

#### 4.1.1 Deep learning methods

Based on architectural backbones, DL methods can be broadly grouped into four categories: FNN-based (GENIUS [25], P-Net [21]), autoencoder-based (TMO-Net [13], CustOmics [14]), graph-based (MOGLAM [17], GNN-SubNet [16], MOGONET [15], MORE [23], MoAGL-SA [24]), and transformer-based (DeePathNet [19], DeepKEGG [18], Pathformer [20]) (Fig. 1c).

For FNN-based methods, P-Net constructs a hierarchical sparse neural network based on gene-pathway and pathway-biological process connections [21]. GENIUS employs convolutional neural networks (CNNs) and integrates prior biological knowledge by converting multi-omics data into “gene images”, using genomic coordinates to define spatial arrangements across omics layers as image channels [25].

Autoencoder-based methods learn low-dimensional representations across omics. Standard dense neural networks are typically employed as backbones. TMO-Net applies multiple variational autoencoders for intermediate fusion, capturing both self- and cross-modal associations [13]. CustOmics uses a hierarchical mixed-integration strategy, learning omics-specific sub-representations via separate autoencoders, which are then combined through a central variational autoencoder [14].

Graph-based methods can be generally divided into two types. Models using samples as graph nodes typically build a sample-level similarity graph for each omics type and perform fusion afterwards. MOGONET fuses the predictive output of each omic-specific graph convolutional network (GCN) in the label space, while MoAGL-SA, MORE, and MOGLAM use GCN or hypergraphs with self-attention mechanism to learn a joint graph representation of multi-omics features [15, 17, 23, 24]. The method using genes as graph nodes (GNN-SubNet) constructs the graph using protein-protein interaction (PPI) network as prior knowledge, learning graph feature representations followed by global-pooling for sample-level predictions [16].

Transformer-based methods integrate pathway knowledge into the model architecture. DeePathNet groups features into pathways and applies self-attention over pathway-level embeddings. Pathformer transforms gene embeddings constructed from per-omic gene-level statistics into pathway embeddings, and uses criss-cross attention for pathway crosstalk modeling. Both DeePathNet and Pathformer use early fusion by integrating multi-omics features before model inputs, while DeepKEGG applies intermediate fusion via omics-specific encoders [18–20].

Most DL methods use *post hoc* feature attribution methods for biomarker identification. These model-agnostic algorithms are applied after training, computing feature scores using gradient-based methods represented by DeepLIFT [62] and Integrated Gradients [63] (IG), as well as other types of methods such as SHAP [64] and GNNExplainer [65]. Some methods use performance drop after zeroing out features (feature ablation) to attribute feature importance. Notably, MOGLAM is an *ante hoc* method, learning a sparsity-regularized feature-indicator matrix during training, which directly encodes feature importance.

#### 4.1.2 Statistical and machine learning methods

In addition to DL methods, multi-omics biomarker identification methods based on more conventional statistical or ML algorithms can be traced earlier in the research line. MCIA [47] was one of the earliest models for multi-omics integration and biomarker identification, performing joint projections of omics into low-dimensional spaces by maximizing their shared covariance structure. MOFA [10] introduced a statistical framework using matrix factorization for factor analysis. More recently, GAUDI [12] applied UMAP-based transformations followed by XGBoost or random forest with SHAP [64] for biomarker identification. These methods are unsupervised without relying on sample-level labels.

In contrast, DIABLO [11] and asmbPLS-DA [48] are supervised learning methods based on sparse generalized canonical correlation analysis (sGCCA) and sparse partial least square discriminant analysis (sPLS-DA), respectively. Both impose sparsity constraints to identify biomarkers through non-zero component loadings. Stabl [40] wraps a sparse base learner (e.g., logistic regression with *l*_1_ regularization) in bootstrap subsampling to compute per-feature selection frequencies, and identify those above a reliability threshold as biomarkers. GDF [22] applies a greedy decision forest over a PPI network using Gini importance for feature scoring. DPM [49] aims at pathway-level modeling but identifies gene-level biomarkers by integrating regulatory directionality (positive/negative associations) across omics with a *p*-value merging algorithm. Unlike DL methods, most statistical and ML methods achieve *ante hoc* biomarker identification via feature weights updated during model optimization.

### 4.2 Collection of gold-standard biomarkers

A critical challenge precluding quantitative evaluation of biomarker identification performance lies in the lack of an off-the-shelf biomarker set that is both clinically validated and aligned with the task of interest. To address this, we collected biomarkers with strong clinical evidence from established oncology biomarker knowledge bases, including CIViC [5], OncoKB [6], and CGI [7]. To remove confounding effects introduced by biomarkers with weak evidence, such as those identified *in silico* by computational models without clinical validation, we harmonized the evidence levels of the three knowledge bases into a unified evidence system (Level A, B, C, and D) as defined by the AMP/ASCO/CAP guidelines [8] (see Supplementary Table 2 for details) and retained only those with Tier I evidence. These biomarkers are supported by professional guidelines, FDA-approved therapies, or well-powered clinical trials [8]. The curated dataset was stratified into prognostic markers (derived from OncoKB and CIViC) and predictive markers (aggregated from all three sources). To ensure consistency with TCGA data, cancer type names in each source were mapped to TCGA project identifiers. Molecular profiles in CIViC were mapped to HGNC gene symbols (Data availability), as were the gene names from the other two sources, to resolve aliases. Therapeutic agent names were harmonized using the standardized terms collected by Ding *et al*. [66], supplemented by a manually defined mapping to resolve investigational codes, salt forms, spelling errors, and formatting inconsistencies. This curation enables a direct and robust assessment of a method’s ability to identify biomarkers that have a high likelihood of succeeding clinical trials and achieving widespread application.

### 4.3 Derivation of gene-level rankings

Since the collected biomarkers are at the gene level, we adopted a gene-centric approach to enable metric calculations. Specifically, for DNA methylation and miRNA, the average scores of the CpG sites or miRNA molecules regulating the same gene were used as the DNAm- or miRNA-type score for that gene, according to regulatory annotations provided by TCGA and miRTarBase (Data availability). For mRNA, CNV, and SNV, and methods with a gene-centric design, including GENIUS, DeePathNet, DeepKEGG, Pathformer, P-Net, GNN-SubNet, GDF, and DPM (Supplementary Methods), this step was omitted since the output scores are already at gene-level. Following this, gene scores from different omics types were aggregated through max-pooling, resulting in a single score for each gene. For methods that output sample-specific feature scores, the gene scores were further averaged across samples within the same class, and then converted to absolute values and averaged across classes. Lastly, genes with zero scores were randomly permuted, and a final gene-level ranking was formed.

### 4.4 Derivation of consensus multi-omics biomarker panels

Performance varied substantially across different omics combinations (Supplementary Figs. 1-15). For this reason, we treated each five-fold experiment separately, where a five-fold experiment refers to a five-fold cross-validation performed within a specific method and omics combination. We first computed an overall performance score for each experiment,

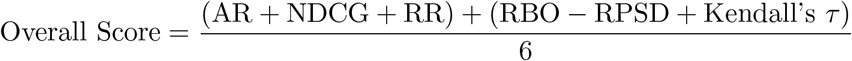

where each metric is averaged across the five folds. All experiments were then ranked in descending order according to this score.

Next, based on the dominant omics type within the top 1% rankings, for each of the five omics types (mRNA, miRNA, CNV, SNV, and DNAm), we selected the top three experiments that included that omics type. For each selected experiment, we applied the Robust Rank Aggregation (RRA) algorithm [58] using the RobustRankAggreg R package (v1.2.1) to integrate the rankings from the five folds. RRA produces an aggregated ranking with Benjamini-Hochberg corrected *p*-values that indicate consensus significance. After obtaining three aggregated rankings for a given omics type, we applied RRA again to integrate these three lists and derive a final consensus ranking for that omics type. During this process, for methods that do not require gene-level inputs (i.e., those except Pathformer, DeePathNet, DPM, GNN-SubNet, GENIUS, GDF, and P-Net; Supplementary Methods), we converted CpG scores to gene level using mean aggregation for DNAm. For gene-centric methods, we converted miRNA scores to gene level. Full results are provided in Supplementary Data 1.

### 4.5 Evaluation metrics

#### 4.5.1 Accuracy metrics

We adopt rank-based metrics for evaluating biomarker identification accuracy. Let *G* denote the gene set in a gene-level ranking list, and let *B* ⊆ *G* be the subset of genes in *G* that are reference biomarkers. Let *r*_*b*_ be the rank of biomarker *b* (*b* ∈ *B, r*_*b*_ ∈ *{*1, 2, …, |*G*|*}*).

**Average Recall**.

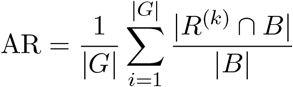

AR ∈ (0, 1), higher means more accurate.

##### Normalized Discounted Cumulative Gain Normalized Discounted Cumulative Gain

(NDCG) has been widely used in information retrieval [67, 68] studies and can measure the prioritization of relevant items in top rankings. The Discounted Cumulative Gain (DCG) is defined as

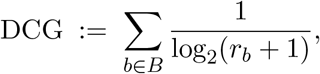

The ideal DCG (IDCG) is obtained by placing all biomarkers at the top of the list:

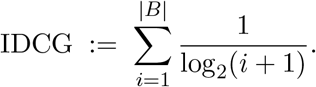

The Normalized DCG is then

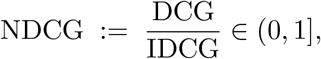

with 1 indicating a perfect ranking in which every biomarker precedes all non-biomarkers.

##### Reciprocal Rank

Reciprocal Rank (RR) is a metric commonly used in information retrieval and question answering to measure how highly the first relevant item is ranked [69]. Let *r*_(1)_:= min_*b*∈*B*_ *r*_*b*_ denote the rank of the highest-ranked biomarker, RR is defined as

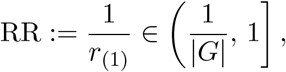

assigning a score of 1 when a biomarker occupies the highest rank and decaying hyperbolically as the first biomarker appears deeper in the ranking list.

##### Mann-Whitney U test *p*-value

We compute a one-sided Mann-Whitney test comparing *{r*_*g*_: *g* ∈ *B}* to *{r*_*g*_: *g* ∈*/B}* using `scipy.stats.mannwhitneyu(alternative=`‘`less`’`, method=`‘`exact`’) (SciPy v1.14.1). The reported quantity is the *p*-value *p* ∈ (0, 1], where smaller *p* indicates stronger evidence that biomarkers rank higher than non-biomarkers.

##### Area Under the ROC Curve

For each *g* ∈ *G*, let *s*_*g*_ be its gene-level score (larger *s*_*g*_ = more biomarker-like) and *y*_*g*_:= **1**[*g* ∈ *B*]. The AUROC is the area under the ROC curve obtained by thresholding *s*_*g*_. Equivalently,

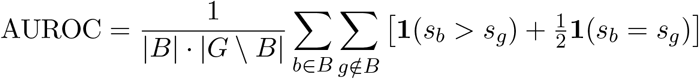

We compute AUROC with `sklearn.metrics.roc_auc_score` (scikit-learn v1.7.2).

**Accuracy@K**. Let *R*^(*K*)^ be the set of the top-*K* genes. Then

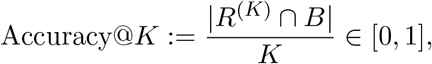

the fraction of the top-*K* predictions that are known biomarkers.

#### 4.5.2 Stability metrics

For stability, we calculated the average similarities between each pair of the five-fold ranking lists.

##### Average Kendall’s *τ*

Kendall’s *τ* [70] is widely used for measuring the overall similarity between two rankings. Given rankings *R*_*i*_ and *R*_*j*_, it is defined as

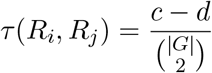

where *c* is the number of concordant pairs (same relative order in both rankings) and *d* the number of discordant pairs. A higher value indicates higher similarity, with 1 denoting perfectly concordant and −1 denoting the least concordant (*τ* ∈ [−1, 1]). We average *τ* over all pairs from the five-fold rankings as the final metric:

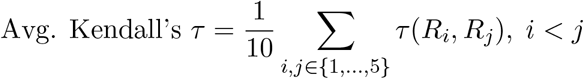

##### Rank-Biased Overlap

Although Kendall’s *τ* offers a global perspective on the overall ranking, including both top-ranked candidates and lower-ranked genes, it may be significantly affected by the instability at lower ranks. We therefore further adopted Rank-Biased Overlap (RBO) [71], a common metric to emphasize top-ranking similarity:

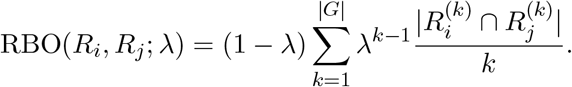

where 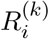 denotes the top-*k* genes in ranking *i*, and *λ* ∈ (0, 1) is the decay rate. Throughout our experiments, *λ* was set to 0.98 to effectively account for the top 50 ranks, as evaluated and reported by Webber *et al*. [71]. RBO∈ [0, 1), with higher values indicating higher concordance. Similar to Kendall’s *τ*, we average RBO over all pairs from the five-fold rankings as the final metric.

##### Rank Percentile Standard Deviation

We define Rank Percentile Standard Deviation (RPSD) to measure the variability of known biomarkers’ ranks across cross-validation folds. For each biomarker *b* and fold *i*, let

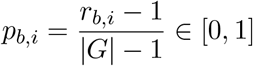

be its rank percentile, and let

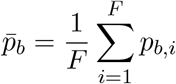

be the mean rank percentile for *b* across *F* folds. We first compute the per-biomarker standard deviation

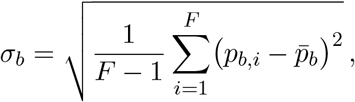

and then average over all biomarkers

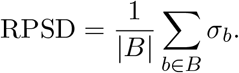

Lower RPSD indicates more consistent ranking of biomarkers across folds, with 0 denoting perfect stability.

## 5 Computational resources

Experiments were primarily conducted on a Linux Ubuntu 22.04 system equipped with 8 NVIDIA RTX A6000 GPUs (48 GB each), a dual-socket AMD EPYC 7763 processor (2*×*64 cores, 2 threads per core), and 1008 GB system memory. Experiments requiring larger GPU memory were run on a Linux Ubuntu 24.04 system equipped with one NVIDIA H200 GPU (SXM, 141 GB), a dual-socket Intel Xeon Platinum 8570 processor (2*×*56 cores, 2 threads per core), and 2 TB system memory.

## 6 Code availability

Our benchmark is publicly available at https://github.com/athanzli/CancerMOBI-Bench.

## 7 Data availability

Our benchmarking datasets and results are available at 10.5281/zenodo.17860662 [72]. TCGA omics and clinical data were downloaded from the GDC data portal at https://portal.gdc.cancer.gov (accessed November 06 2024). Reference biomarkers were retrieved from https://www.oncokb.org/actionable-genes for OncoKB (accessed June 15 2025), https://civicdb.org/releases/main for CIViC (2025-06-01 release), and https://www.cancergenomeinterpreter.org/data/biomarkers for CGI (latest version, accessed June 15 2025). The HGNC complete gene set information was downloaded from https://storage.googleapis.com/public-download-files/hgnc/tsv/tsv/hgnc_complete_set.txt (accessed Feb 11 2025). miRNA target gene information was retrieved from miRTarBase at https://mirtarbase.cuhk.edu.cn/~miRTarBase/ (accessed April 21 2025). Methylation array manifest file (`HM450.hg38.manifest.gencode.v36.tsv.gz`) and TCGA antibodies descriptions file (`TCGA_antibodies_descriptions.gencode.v36.tsv`) were downloaded from https://gdc.cancer.gov/about-data/gdc-data-processing/gdc-reference-files. All biological pathway data files were downloaded from the corresponding methods’ data repositories. PPI network topology file was downloaded from the STRING [73] database at https://string-db.org/ (accessed October 28 2024).

## 8 Author contributions statement

R.L. conceived and supervised the study. A.Z.L. conducted the literature review. A.Z.L. designed the methodology. A.Z.L. developed the software. A.Z.L. generated the visualizations. A.Z.L., Y.D., Y.L., L.C., and R.L. analyzed the results. A.Z.L. and R.L. drafted the manuscript.

## 11 Author description

Athan Z. Li is a Ph.D. student in the department of Computer Science at the University of Southern California.

Yuxuan Du is an Assistant Professor of Electrical Engineering at The University of Texas at San Antonio.

Yan Liu is a Professor of Computer Science with a joint appointment with Electrical Engineering at the University of Southern California.

Liang Chen is a Professor of Quantitative and Computational Biology at the University of Southern California.

Ruishan Liu is an Assistant Professor of Computer Science with a joint appointment with Quantitative and Computational Biology at the University of Southern California.

