## Supplementary Material for "Benchmarking computational methods for multi-omics biomarker discovery in cancer"

### 1 Supplementary Methods

#### 1.1 Selection of benchmarked methods

We searched the literature through a combination of keywords regarding multi-omics integration and biomarker identification, and gathered 82 computational multi-omics biomarker identification methods in Supplementary Table 1. To select the methods for benchmarking, we applied three essential criteria: methodological compatibility, accessibility/usability, and computational expense. Specifically, methods were included in benchmarking if they satisfy the following criteria:

##### 1.1.1 Methodological compatibility

(1) The method should be a general-purpose approach for multi-omics data integration, capable of accepting other omics types rather than being limited to the omics types demonstrated in the original case studies. This flexibility is essential for deployment, and for comparative benchmarking to ensure all methods are evaluated on identical omics combinations. (2) The biomarker identification module must output a global feature ranking or scoring list and should not produce an excessively sparse set of biomarkers. If only a small number of features receive non-zero scores, there is a high likelihood that these features will be unvalidated biomarkers absent from reference knowledge bases, while validated biomarkers cannot be evaluated using rank-based metrics. (3) For methods that generate sample-level predictions, classification must be supported.

##### 1.1.2 Accessibility and usability:

(1) Code required to run the method must be publicly available and complete, covering both the model training/inference and biomarker identification components. (2) The installation and execution of the codebase should not contain major errors. (3) Code should be implemented in Python or R and engineered in such a way that it can be integrated into our benchmarking pipeline either directly or after necessary modifications, without requiring prohibitive re-engineering efforts. (4) If the method relies on preprocessed prior knowledge data files, these files must be available for download. (5) Hyperparameters or guidance for setting them should be clearly documented, either in the manuscript, code, or accompanying documentation.

##### 1.1.3 Computational expense

The method must complete execution within one day on a typical real-world dataset (approximately 400 samples $\times$ 40,000 features) for a single run.

#### 1.2 TCGA Data preprocessing

##### 1.2.1 TCGA omics data

TCGA multi-omics data were downloaded from the Genomic Data Commons (GDC) Data Portal (see Data availability in the main text). Based on the omics type choices by the benchmarked methods (Supplementary Table 4) and their availability in TCGA, we opted for five omics data types, namely messenger RNA (mRNA), microRNA (miRNA), DNA methylation (DNAm), Copy Number Variation (CNV), and Single Nucleotide Variation (SNV). Proteomics data were excluded for real-world data benchmarking because TCGA RPPA quantifies only a pre-selected panel of high-quality antibodies (n=487) targeting canonical cancer-pathway proteins[1], which introduces bias towards identifying well-known targets. For mRNA-seq, we used TPM (unstranded) from STAR-Counts, retained protein-coding genes, and applied log2-transformation. For miRNA-seq, we used reads-per-million (RPM) miRNA expression quantifications and performed log2-transformation. For DNA methylation (Illumina HumanMethylation450), we used SeSAMe beta-values and removed CpGs without gene annotations in the Illumina manifest file (see Data availability in the main text). For SNVs, we used the masked somatic mutation data, and assigned 1 if a variant mapped to a gene, else 0. For CNVs (Affymetrix SNP 6.0), gene-level copy numbers were restricted to protein-coding genes, centered by subtracting the diploid baseline (2 copies), and capped at 2 for amplifications. Based on GISTIC2.0[2], values were grouped into five levels of alterations, namely deep deletion (-2), shallow deletion (-1), diploid (0), gain (1) and amplification (2). For simulated data which requires proteomics as reference, we used RPPA protein expression profiles, removed features without a corresponding gene in the TCGA antibodies descriptions file (see Data availability in the main text), and excluded samples with more than 50% missing features. After each task dataset was constructed, features with missing values or zero variance were further removed. Finally, standard scaling was applied to each task dataset by fitting train sets and transforming validation and test sets.

##### 1.2.2 TCGA survival and drug response data

Based on the clinical data file from TCGA (Data availability), for survival, we first excluded patient samples without available survival time information ('days_to_death' or 'days_to_last_follow-up'). Then in each task dataset, samples with survival times longer than the median were categorized as high-risk, and those below the median as low-risk. For drug response data, we first excluded patient samples without available 'therapeutic_agent' (for drug labels) or 'treatment_outcome' (for clinical response labels) annotations. Then, drug-patient pairs with inconsistent treatment outcomes were removed. Afterwards, clinical responses were binarized into response (complete response, partial response) and non-response (progressive disease, stable disease, no response, and persistent disease) according to the RECIST standard[3]. Drug names were standardized using the TCGA drug name standardization information from Ding *et al.*[4], supplemented by a manually defined mapping to resolve investigational codes, salt forms and formatting inconsistencies.

#### 1.3 Construction of task datasets

To construct task datasets for biomarker identification, we followed multiple filtering steps to achieve both computational feasibility and biomarker-task alignment.

First, we ensured that the candidate tasks align with the evidence types of biomarkers. Particularly, it was observed that the prognostic and therapeutic biomarkers constitute the primary portion of all the collected biomarker profile entries (Supplementary Fig. 23). Therefore, we opted for two major tasks, survival risk classification and drug response prediction, consistent with the primary evaluation tasks employed in the original studies[5–9].

Second, we filtered candidate tasks by sample and biomarker counts. For survival risk classification tasks, sample size and biomarker counts for each cancer type are summarized in Supplementary Table 6, and only those with $\geq$ 300 samples and $\geq$ 10 biomarkers were further considered. Here, only samples with all five omics data types (i.e., mRNA, miRNA, DNAm, CNV, and SNV) available were counted. Following this, we inspected each candidate cancer type's biomarker evidence statements in the source knowledge records, and excluded those with low rating or non-significant correlation with the corresponding cancer type's prognosis (Supplementary Table 7). After this, BRCA, LUAD, LUSC, and COADREAD remained as candidates with $\geq$ 10 biomarkers. For drug response prediction tasks, the sample and biomarker numbers for each drug-cancer pair are summarized into Supplementary Table 8 using TCGA clinical data file. Samples with missing omics data from the five omics types were further removed, and the drug-cancer-type pairs with $\geq$ 20 responders and $\geq$ 20 non-responders were selected. Manual inspections on biomarker evidence statements were also conducted to ensure alignment with drug efficacy, and no biomarker was excluded during this process (Supplementary Table 7). Finally, the following drug-cancer-type pairs were considered as candidates (#responders:#non-responders): Cisplatin-BLCA (40:20), Fluorouracil-STAD (51:29), Gemcitabine-PAAD (28:33), Temozolomide-LGG (20:103).

For each candidate task, we prepared its corresponding multi-omics dataset using samples with mRNA, miRNA, DNAm, CNV, and SNV data, along with their survival or drug response labels. Features with missing values or zero variance across samples were removed. Stratified splitting was then applied to generate five cross-validation folds, each dividing the data into an 80% train-validation set and a 20% test set. The train-validation portion was further split into training and validation subsets at a 7:1 ratio, leading to final proportions of 70% training, 10% validation, and 20% testing. For survival tasks, survival times were first grouped into 20 quantile bins before stratification for similar time distributions across the training, validation, and test sets.

Third, for each candidate task dataset, we used basic machine learning classifiers to test whether each dataset possessed sufficient signal for classification. Datasets whose classification performance is below the random baseline were discarded. This guarantees the benchmarked methods can capture distinguishable signals and thus provide trustworthy feature scores or ranks for biomarker identification. To achieve this, we used a random forest and a support vector machine as classifiers to run the 30 experiments (`sklearn.ensemble.RandomForestClassifier` and `sklearn.svm.LinearSVC` from scikit-learn v1.7.2, with default parameters), including 6 omics combinations across 5 cross-validation folds. Tasks with at least one result below the random baseline (mean cross-validation AUROC = 0.5) were excluded, and the remaining ones were selected as the final tasks (Supplementary Table 9). The final benchmarking consists of five multi-omics datasets: BRCA, LUAD, and COADREAD for survival risk classification, and BLCA treated with cisplatin and LGG treated with temozolomide for the drug response prediction. The 43 matched biomarkers are listed in Supplementary Table 3, the task dataset statistics are summarized in Table 2 in the main text, and methods' predictive performance is reported in Supplementary Tables 10–14.

#### 1.4 Simulation of multi-omics data

We employed InterSIM[10] (v2.3.0), a multi-omics data generation tool that simulates DNA methylation, gene expression, and protein expression based on key statistics of real reference data for generating simulated multi-omics datasets. InterSIM has been applied in previous benchmark studies of multi-omics integration methods[8,11], and is balanced between the complexity and interpretability of its generation for different sample groups, making it a suitable tool for our benchmarking.

InterSIM provides the option to use its built-in TCGA ovarian cancer dataset as reference for simulation, but it was limited to a few hundred CpGs, genes, and proteins. To better align with the statistical properties of real-world data such as high dimensionality, and to allow running prior-knowledge-based methods, we used real reference based on the BRCA survival task dataset due to its largest sample size (Table 2 in the main text). Specifically, we first included proteomics from RPPA as an additional omics type, and removed samples without proteomics data in the BRCA survival task dataset. Then, for a feasible generation time (less than one day per simulation), we removed CpG sites with less than 0.025 variance or without corresponding gene labels in the methylation array manifest file (Data availability). Following this, the genes regulated by the remaining CpGs were kept, and proteins without a corresponding gene in the RPPA antibodies description file were removed, leaving 6273 CpGs, 3862 genes, and 95 proteins as reference.

For simulation, InterSIM first generates methylation M-values $\boldsymbol{X}_1 \in \mathbb{R}^{N \times F_1}$ for $N$ samples and $F_1$ CpGs via a multivariate Gaussian distribution

$$\boldsymbol{X}_1 \sim \mathcal{N}(\boldsymbol{\mu}_1 + \delta_1 \boldsymbol{d}_1, \boldsymbol{\Sigma}_1),$$

where $\boldsymbol{\mu}_1 \in \mathbb{R}^{F_1}$ is the reference mean M-values across samples at each CpG site, and $\boldsymbol{\Sigma}_1 \in \mathbb{R}^{F_1 \times F_1}$ is the reference covariance matrix, both pre-calculated from reference data. For simulating ground truth biomarkers, $\boldsymbol{d}_1 \in \{0,1\}^{F_1}$ is an indicating vector generated by Bernoulli trials with $p = 0.01$ for differentially methylated CpGs, and $\delta_1$ determines the power of mean shift.

Afterwards, the reference methylation data is converted into gene-level by taking the median M-values of CpGs mapping to the same gene, and the Pearson correlation coefficients $\boldsymbol{\rho_{12}} \in \mathbb{R}^{F_2}$ is computed between the median M-values and the corresponding gene expression for each gene, where $F_2$ denotes the number of genes. Then, gene expression data is simulated via

$$\boldsymbol{X}_2 \sim \mathcal{N}((\boldsymbol{\rho}_{12} \circ \boldsymbol{\mu}_1^{(g)} + \sqrt{1 - \boldsymbol{\rho}_{12}^2} \circ \boldsymbol{\mu}_2) + \delta_2 \boldsymbol{d}_2, \boldsymbol{\Sigma}_2),$$

where $\boldsymbol{\mu}_1^{(g)} \in \mathbb{R}^{F_2}$ is the reference vector for mean gene-level methylation, and $\boldsymbol{\mu}_2$ is the reference mean gene expression. The differentially expressed gene indicating vector $\boldsymbol{d}_2 \in \{0,1\}^{F_2}$ is determined by the regulatory mappings between CpGs and genes according to real reference data. Particularly, a gene is a biomarker if one of its regulating CpG is differentially methylated as in $\boldsymbol{d}_1$. $\boldsymbol{\Sigma}_2 \in \mathbb{R}^{F_2 \times F_2}$ is the reference covariance matrix calculated from real gene expression data, and $\delta_2$ determines the power of mean shifts.

For protein expression, first, the Pearson correlation coefficients $\boldsymbol{\rho_{23}} \in \mathbb{R}^{F_3}$ is computed between the expression of each protein and the corresponding gene, where $F_3$ denotes the number of proteins. Then, protein expression is simulated via

$$\boldsymbol{X}_3 \sim \mathcal{N}((\boldsymbol{\rho}_{23} \circ \boldsymbol{\mu}_2^{(p)} + \sqrt{1 - \boldsymbol{\rho}_{23}^2} \circ \boldsymbol{\mu}_3) + \delta_3 \boldsymbol{d}_3, \boldsymbol{\Sigma}_3),$$

where $\boldsymbol{\mu}_2^{(p)} \in \mathbb{R}^{F_3}$ is the reference vector of mean gene expression corresponding to proteins, and $\boldsymbol{\mu}_3$ is the reference mean protein expression. The differentially expressed protein indicating vector $\boldsymbol{d}_3 \in \{0,1\}^{F_3}$ is determined by the regulatory mappings between genes and proteins, as in the real reference data. $\boldsymbol{\Sigma}_3 \in \mathbb{R}^{F_3 \times F_3}$ is the reference covariance matrix calculated from real protein expression data, and $\delta_3$ determines the power of mean shifts.

Notably, given the significant difference between the number of reference genes (3862) and proteins (95), it is possible that $\boldsymbol{d}_3 = \boldsymbol{0}$ when the Bernoulli trials for $\boldsymbol{d}_1$ did not produce any CpG regulating a protein-corresponding gene. Therefore, we separated the set of protein-corresponding genes for an additional Bernoulli trial with $p = 0.1$, and combined the resulting indicating vector with the original one as the final indicating vectors for both gene expression and protein expression data. To simulate datasets with a range of biomarker signal strengths, we varied the mean shifts parameter by $0.5, 1, 2, 3, 4, 5$ ($\delta_1 = \delta_2 = \delta_3$). All simulations generate datasets with 100 samples for two groups, with 50 samples in each. Finally, the generated methylation M-values were converted to beta-values via logit-transformation. Train, validation, and test sets splitting and standard scaling were applied in the same way as for real task datasets. The statistics of the simulated datasets are summarized in Supplementary Table 5.

#### 1.5 Implementation and experimental setups for the benchmarked methods

##### 1.5.1 General experimental settings

All DL methods were incorporated into unified Python pipelines with PyTorch, using their publicly available code. Unless noted otherwise (for example, MOGONET, MOGLAM, MORE, MoAGL-SA, and Pathformer, which employ fewer training epochs by default), each model was trained for up to 1,000 epochs. Early stopping with a patience of 100 epochs was applied to all models, and the checkpoint with the lowest validation loss was retained for both prediction and biomarker identification. Hyperparameters were set according to the default optimal values reported in the original publications or associated codebases. To account for class imbalance in drug response tasks, class weights were introduced into the cross-entropy loss. Concretely, for $C$ classes with sample sizes $n_1, n_2, \ldots, n_C$, the weight for class $i$ was computed as $w_i = \frac{N}{C\, n_i}$, for $i = 1, 2, \ldots, C$, where $N$ is the total number of samples. All methods were evaluated using five-fold cross-validation within each task dataset. For *post hoc* feature attribution methods, including Integrated Gradients (IG), SHAP, and DeepLIFT, feature scoring was performed on the test sets, with the training set provided as background data for SHAP. To overcome memory constraints without affecting the attribution results, mini-batch processing was used for IG (step size = 50) and SHAP. Statistical and ML methods were incorporated using their publicly available code or documentation, with default optimal parameters. For methods implemented in R, we used the Python package `rpy2` (v3.5.16) to integrate into our unified Python pipeline.

##### 1.5.2 Method-specific implementation and experimental settings

**P-Net** P-Net was originally implemented with TensorFlow. To conform with our benchmarking pipeline, which uses PyTorch, we used the PyTorch implementation of P-Net from Pathformer's repository `https://github.com/lulab/Pathformer/tree/main/comparison_methods/P_net`. The Reactome pathway data files were retrieved from the same repository. We implemented the biomarker identification module strictly according to the method descriptions from P-Net's manuscript, while also referring to their original code at `https://github.com/marakeby/pnet_prostate_paper`. All parameters and training setups were ensured to be the same as P-Net's original settings.

**GENIUS** We used the code from `https://github.com/mxs3203/GENIUS` with default parameters. The file for the physical positions of genes on genome was downloaded from the same repository.

**TMO-Net** We used the code from `https://github.com/FengAoWang/TMO-Net` with default parameters.

**CustOmics** We used the code from `https://github.com/HakimBenkirane/CustOmics` with default parameters.

**MOGONET** We used the code from `https://github.com/txWang/MOGONET`, with default parameters.

**MoAGL-SA** We used the code from `https://github.com/gpxzmu/MoAGL-SA`, with default parameters.

**MORE** We used the code from `https://github.com/Wangyuhanxx/MORE`, with default parameters.

**MOGLAM** We used the code from `https://github.com/Ouyang-Dong/MOGLAM` with default parameters.

**GNN-SubNet** We used the code from `https://github.com/pievos101/GNN-SubNet` with default parameters. PPI data was downloaded from StringDB[12] (see Data Availability in the main text), and a combined score threshold of 0.95 was adopted as in GNN-SubNet's original setting.

**Pathformer** We used the code from `https://github.com/lulab/Pathformer` with default parameters. The files for biological pathways and precomputed pathway cross-talk weights were retrieved from the same repository.

**DeePathNet** We used the code from `https://github.com/CMRI-ProCan/DeePathNet`, with default parameter configuration. The LCPathway[13] data file was retrieved from `https://figshare.com/articles/dataset/Intermediate_files_for_reproducing_results/24137619`.

**DeepKEGG** DeepKEGG was originally implemented under TensorFlow. To conform with our PyTorch-based pipeline, we implemented a PyTorch-based version by using their original code from `https://github`

.com/lanbiolab/DeepKEGG as strict reference. Default parameters as in the original implementation were adopted, and the KEGG pathway data file was retrieved from the same repository.

**DIABLO** We used the mixOmics (v6.28.0) package, and followed the tutorial at https://mixomics.org/mixdiablo/diablo-tcga-case-study/. Specifically, ncomp was set to 3 according to the reported tuning result. keepX (i.e., the number of features to keep for each omics type) was set to 1% of each omics type's total feature number, and the predictive performance was reported using the model with the 1% keepX. To obtain a single set of biomarkers from the weights of multiple components, the authors took the union of the non-zero weighted features across components. As a similar approach, we opted for max-pooling along the components to obtain a single weight for each feature. The selected features are ranked according to their absolute weights. To assign ranks to the remaining zero-weighted features, we ran DIABLO again with the default argument for keepX to obtain non-zero weights for all features, and the remaining features are ranked following the ranks of the top 1% features. We treat the reversed rankings as scores before deriving gene-level rankings following the procedure detailed in Methods of the main text.

**MCIA** We used the omicade4 (v1.44.0) R package, and followed the tutorial at https://www.bioconductor.org/packages/devel/bioc/vignettes/omicade4/inst/doc/omicade4.pdf. Max-pooling was adopted to use the largest absolute weight across axes for each feature.

**MOFA** We used the python implementation of MOFA with package mofapy2 (v0.7.2), and followed the tutorial at https://github.com/bioFAM/mofapy2/blob/master/mofapy2/notebooks/getting_started_python.ipynb. Max-pooling was adopted to use the largest absolute weight across factors for each feature.

**GAUDI** We followed the tutorial at https://github.com/hirscheylab/gaudi, with default parameter settings. We opted for the random forest method for broader feature scoring, as suggested by the authors. Also, per the authors' recommendation, we reduce the default n_neighbors to 5 for the drug response tasks due to smaller sample size.

**asmbPLS-DA** We followed the guide at https://github.com/RunzhiZ/asmbPLS, with default parameter settings. Similar to the setting for DIABLO described above, we set 0.99 as the quantile for each block (this corresponds to the quantile.comb argument in asmbPLSDA.fit) to retain 1% features. The largest absolute weight across axes is used as the score for each feature, and features are ranked according to the absolute values of these scores. The rest features are ranked following the previous ranks based on the absolute scores obtained by re-running asmbPLS-DA with a 0 quantile. We treat the reversed rankings as scores before deriving gene-level rankings following the procedure detailed in Methods of the main text.

**Stabl** We followed the guide at https://github.com/gregbellan/Stabl, with default parameter settings. In particular, we set the argument artificial_type to 'knockoff' to account for the internally correlated omics data, as demonstrated in the manuscript's analysis.

**GDF** We used the code from https://github.com/pievos101/DFNET, with default parameters. PPI data was downloaded from StringDB[12] (see Data Availability in the main text), and a combined score threshold of 0.95 was adopted as in GDF's original setting.

**DPM** We followed the tutorial of directional $p$-value merging at https://github.com/reimandlab/ActivePathways. Mann-Whitney U test was adopted for obtaining $p$-values, as in the paper's analysis for omics data. For the constraints vector required by DPM input, we encoded expected cross-omics relationships with $c \in \{-1, 0, +1\}^K$. Specifically, we set mRNA$= +1$, CNV$= +1$, DNAm$= -1$, miRNA$= -1$, and $SNV = 0$. Since DPM outputs $p$-values, we applied $-log_{10}$ to obtain gene-level scores.

#### 1.6 Detailed description of the benchmarked methods

Here we provide a detailed description of each benchmarked method, focusing on multi-omics integration, sample predictions, and molecular biomarker identification. For further details, readers are advised to refer to the corresponding manuscripts.

##### 1.6.1 P-Net

Elmarakeby *et al.*[14] proposed P-Net, a biologically informed sparse deep neural network whose neurons are connected by known parent-child relationships between genes, pathways, and biological processes from the Reactome pathway database[15].

**Inputs.** P-Net follows a gene-centric design by mapping omics features to gene level. For $G$ genes and $K$ omics types, each sample $s$ provides $K$ per-gene inputs $\mathbf{x}_g^{(s)} \in \mathbb{R}^K$. For each gene, the gene layer aggregates these by

$$z_{g,s}^{(1)} = \sum_{t=1}^{K} w_{g,t}^{(1)}\, x_{g,t}^{(s)} + b_g^{(1)}, \qquad h_{g,s}^{(1)} = \tanh\big(z_{g,s}^{(1)}\big).$$

where $w_{g,t}^{(1)}$ is a learnable weight parameter and tanh is the hidden activation function.

**Architecture.** Connectivity is enforced by a binary mask $\mathbf{M}^{(\ell)}$ that zeros out connections absent from Reactome knowledge. For hidden layers $\ell = 2, \ldots, L-1$,

$$\mathbf{h}_s^{(\ell)} = \tanh\Big(((\mathbf{W}^{(\ell)} \odot \mathbf{M}^{(\ell)})^\top \mathbf{h}_s^{(\ell-1)}) + \mathbf{b}^{(\ell)}\Big).$$

where $\mathbf{W}^\ell$ are learnable weight parameters and $\mathbf{b}^\ell$ is the bias term. A sigmoid predictive head is attached to each hidden layer:

$$p_s^{(\ell)} = \sigma\big(\mathbf{v}^{(\ell)\top}\mathbf{h}_s^{(\ell)} + a^{(\ell)}\big), \qquad \hat{p}_s = \frac{1}{|\mathcal{H}|}\sum_{\ell\in\mathcal{H}} p_s^{(\ell)},$$

where $\mathbf{v}^{(\ell)} \in \mathbb{R}^{n_\ell}$ is the predictive-head weight vector, $a^{(\ell)} \in \mathbb{R}$ is its scalar bias, $\mathbf{h}_s^{(\ell)} \in \mathbb{R}^{n_\ell}$ is the hidden activation vector at layer $\ell$ for sample $s$, $n_\ell$ is the number of nodes in layer $\ell$, $\sigma(z) = 1/(1+e^{-z})$, and $\mathcal{H} = \{1, \ldots, L-1\}$.

**Training objectives.** Training is performed using binary cross-entropy loss with class and layer weightings that emphasize later layers. For class weights $(w_1, w_0)$ and head weights $\alpha_\ell$,

$$\mathcal{L} = \frac{1}{|\mathcal{B}|}\sum_{s\in\mathcal{B}}\sum_{\ell\in\mathcal{H}} \alpha_\ell \left[-w_1\, y_s \log p_s^{(\ell)} - w_0\,(1-y_s)\log(1-p_s^{(\ell)})\right],$$

where $\mathcal{B}$ represents the samples in one training batch.

**Biomarker identification.** Gene-level node attributions are computed with DeepLIFT [16] w.r.t. the final prediction. Let $c_{i,s}$ denote the DeepLIFT contribution of the gene-layer node $i$ for sample $s$, then the total node importance is the absolute value of the sum over the test set,

$$I_i = \Big|\sum_{s\in\mathcal{S}_{\text{test}}} c_{i,s}\Big|.$$

The final importance score is adjusted by node degree. Specifically, if $d_i$ is the degree of node $i$, with layer-wise mean $\mu$ and standard deviation $\sigma$,

$$\tilde{I}_i = \begin{cases} I_i/d_i, & d_i > \mu + 5\sigma, \\ I_i, & \text{otherwise.} \end{cases}$$

##### 1.6.2 GENIUS

Sokač *et al.* [17] proposed GENIUS, a framework that transforms multi-omics features into spatial "genome images", with genes as pixels and omics features as channels. GENIUS uses convolutional neural networks (CNNs) as backbones, and gene-level importance is attributed by Integrated Gradients (IG) [18].

**Inputs.** Given $M$ omics types measured on a common gene set $\mathcal{G}$ and sample $s$ with per-gene features $\mathbf{X}^{(s)} \in \mathbb{R}^{|\mathcal{G}|\times M}$, GENIUS builds a multi-channel image $\mathbf{I}^{(s)} \in \mathbb{R}^{H\times W\times M}$ via a fixed gene-to-pixel mapping $\pi : \mathcal{G} \to [H] \times [W]$:

$$\mathbf{I}_{u,v,m}^{(s)} \;=\; x_{g,m}^{(s)} \quad \text{for } (u,v) = \pi(g),$$

where $x_{g,m}^{(s)}$ is the preprocessed feature value of omics $m$ for gene $g$, and $\pi$ places genes by chromosomal position while ordering chromosomes by Hi-C interaction strength [17]. Specifically, $H = W = 198$.

**Architecture.** A CNN processes $\mathbf{I}^{(s)}$ with four parts: encoder, decoder, extractor, and predictor. Let

$$\mathbf{L}^{(s)} = f_{\text{enc}}\big(\mathbf{I}^{(s)}; \theta_e\big) \in \mathbb{R}^{128}, \quad \widehat{\mathbf{I}}^{(s)} = f_{\text{dec}}\big(\mathbf{L}^{(s)}; \theta_d\big), \quad \mathbf{z}^{(s)} = f_{\text{ext}}\big(\widehat{\mathbf{I}}^{(s)}; \theta_x\big),$$

For classifications, the prediction head concatenates features with $\mathbf{L}^{(s)}$:

$$\mathbf{o}^{(s)} = \mathbf{W}_p\,[\mathbf{z}^{(s)}; \mathbf{L}^{(s)}] + \mathbf{b}_p, \qquad \hat{\mathbf{p}}^{(s)} = \text{softmax}\big(\mathbf{o}^{(s)}\big)$$

where $f_{\text{enc}}, f_{\text{dec}}, f_{\text{ext}}$ are convolutional subnetworks with parameters $\theta_e, \theta_d, \theta_x$, $\mathbf{L}^{(s)}$ is the latent vector, $[\cdot\,;\cdot]$ denotes concatenation, $\mathbf{W}_p, \mathbf{b}_p$ are predictor parameters.

**Training objectives.** For classification with labels $y_s \in \{1, \ldots, C\}$ and probabilities $\hat{\mathbf{p}}^{(s)}$,

$$\mathcal{L}_{\text{clf}} = -\frac{1}{|\mathcal{B}|} \sum_{s \in \mathcal{B}} \log \hat{\mathbf{p}}^{(s)}_{y_s}$$

where $\mathcal{B}$ represents the training batch.

**Biomarker identification.** After training, IG is computed for each pixel (gene-level) per channel. For baseline $\mathbf{I}^{(0)}$ and scalar output $F(\mathbf{I})$,

$$\text{IG}_i\big(\mathbf{I}^{(s)}\big) = \big(\mathbf{I}^{(s)}_i - \mathbf{I}^{(0)}_i\big) \int_0^1 \frac{\partial F\big(\mathbf{I}^{(0)} + \alpha(\mathbf{I}^{(s)} - \mathbf{I}^{(0)})\big)}{\partial \mathbf{I}_i} \, d\alpha,$$

where $i$ is the index for a particular pixel-channel. Per-gene attributions for sample $s$ are

$$S^{(s)}_g = \sum_{m=1}^{M} \text{IG}_{(\pi(g),m)}\big(\mathbf{I}^{(s)}\big)$$

channel-wise $S^{(s,m)}_g = \text{IG}_{(\pi(g),m)}(\cdot)$ allows scoring per-omic importances.

##### 1.6.3 TMO-Net

Wang *et al.*[7] proposed TMO-Net, an explainable, pre-trained multi-omics model built on self- and cross-modal variational autoencoders (VAEs) with a product-of-experts fusion that learns cross-omics interactions.

**Inputs.** Let $\{\mathbf{X}^{(m)}\}_{m=1}^{M}$ be omics feature matrices with $\mathbf{X}^{(m)} \in \mathbb{R}^{N \times d_m}$ and per-sample vectors $\mathbf{x}^{(s,m)} \in \mathbb{R}^{d_m}$. Each sample's multi-omics input is $\{\mathbf{x}^{(s,1)}, \ldots, \mathbf{x}^{(s,M)}\}$.

**Architecture:** Per-omic self-encoders produce latent $\mathbf{z}^{(s)}_{m,\text{self}} \sim q_\phi(\mathbf{z_m} \mid \mathbf{x}^{(s,m)})$. Cross-encoders infer a latent for target omic $m$ from other observed omics $\nu^m \subseteq \{1, \ldots, M\} \backslash \{m\}$ using a product-of-expert posterior

$$q_\phi\big(\mathbf{z_m} \mid \mathbf{x}^{(s,\nu^m)}\big) = \prod_{m' \in \nu^m} q_\phi\big(\mathbf{z_m} \mid \mathbf{x}^{(s,m')}\big),$$

and draw $\mathbf{z}^{(s)}_{m,\text{cross}} \sim q_\phi(\mathbf{z_m} \mid \mathbf{x}^{(s,\nu^m)})$. The per-omic fused latent and the joint embedding are

$$\mathbf{z}^{(s)}_{m,\text{fusion}} = \begin{cases} \mathbf{z}^{(s)}_{m,\text{self}} \| \mathbf{z}^{(s)}_{m,\text{cross}}, & \text{if } \mathbf{x}^{(s,m)} \text{ observed,} \\ \mathbf{z}^{(s)}_{m,\text{cross}}, & \text{if } \mathbf{x}^{(s,m)} \text{ missing,} \end{cases} \qquad \mathbf{z}^{(s)}_{\text{joint}} = \|_{m=1}^{M} \mathbf{z}^{(s)}_{m,\text{fusion}}.$$

where $q_\phi(\cdot)$ are variational posteriors parameterized by encoders, and $\|$ denotes concatenation.

**Training objectives.** Pre-training uses a multi-objective self-supervised loss combining self-modal evidence lower bound (ELBO), cross-modal ELBO, discriminator, and contrastive terms. For downstream classification tasks, with logits

$$\mathbf{o}^{(s)} = f_{\text{clf}}\big(\mathbf{z}^{(s)}_{\text{joint}}; \theta_{\text{clf}}\big), \qquad \hat{\mathbf{p}}^{(s)} = \text{softmax}(\mathbf{o}^{(s)}),$$

train with loss

$$\mathcal{L}_{\text{CE}} = -\frac{1}{|\mathcal{B}|} \sum_{s \in \mathcal{B}} \sum_{c=1}^{C} \mathbf{y}_{s,c} \log \hat{\mathbf{p}}^{(s)}_c,$$

where $\mathbf{y}_{s,\cdot}$ is a one-hot label.

**Biomarker identification.** Integrated Gradients (IG) attribute predictions to input features. For baseline $x'$ and scalar output $F(x)$,

$$\text{IG}_i(x) = \big(x_i - x'_i\big) \int_0^1 \frac{\partial F\big(x' + \alpha(x - x')\big)}{\partial x_i} \, d\alpha, \qquad I(i) = \frac{1}{|S|} \sum_{x \in S} \big|\text{IG}_i(x)\big|,$$

where $i$ indexes an input feature, and $\mathcal{S}$ is the sample set over which global importance is averaged.

##### 1.6.4 CustOmics

Benkirane *et al.*[19] proposed CustOmics, a two-phase, mixed-integration deep-learning framework that first learns source-specific representations with per-omic autoencoders and then fuses them with a central variational autoencoder.

**Inputs.** Let $\{\mathbf{X}^{(m)}\}_{m=1}^{M}$ be omics features with $\mathbf{X}^{(m)} \in \mathbb{R}^{N\times d_m}$ and per-sample vectors $\mathbf{x}^{(s,m)} \in \mathbb{R}^{d_m}$.

**Architecture.** *Phase 1 (source-specific).* For each omics type $m$:

$$\mathbf{z}^{(s,m)} = f_{\text{enc}}^{(m)}\big(\mathbf{x}^{(s,m)};\theta_e^{(m)}\big), \qquad \hat{\mathbf{x}}^{(s,m)} = f_{\text{dec}}^{(m)}\big(\mathbf{z}^{(s,m)};\theta_d^{(m)}\big).$$

*Phase 2 (fusion).* Concatenate sub-representations and learn a joint representation with a central VAE:

$$\mathbf{z}^{(s)} = f_{\text{enc}}^{\text{central}}\Big(\big[\mathbf{z}^{(s,1)};\ldots;\mathbf{z}^{(s,M)}\big];\theta_c\Big), \qquad \hat{\mathbf{z}}^{(s)} = f_{\text{dec}}^{\text{central}}\big(\mathbf{z}^{(s)};\tilde{\theta}_c\big),$$

then a task head reads $\mathbf{z}^{(s)}$:

$$\mathbf{o}^{(s)} = f_{\text{clf}}\big(\mathbf{z}^{(s)};\theta_{\text{clf}}\big), \qquad \hat{\mathbf{p}}^{(s)} = \text{softmax}\big(\mathbf{o}^{(s)}\big)$$

where $[\cdot\,;\cdot]$ denotes concatenation; $f_{\text{enc}}^{(m)}, f_{\text{dec}}^{(m)}$ are per-omic encoders and decoders, respectively, and $f_{\text{enc}}^{\text{central}}, f_{\text{dec}}^{\text{central}}$ form the central variational autoencoder.

**Training objectives.** For classification, training proceeds in two phases, each including reconstruction and classification objectives.

*Phase 1 (source-specific).* For each omics source $m$,

$$\mathcal{L}^{(1,m)} = \mathcal{L}_{\text{rec}}^{(m)} + \alpha\,\mathcal{L}_{\text{CE}}^{(m)}, \qquad \mathcal{L}^{(1)} = \sum_{m=1}^{M}\mathcal{L}^{(1,m)}.$$

where $\alpha$ is a weight hyperparameter.

*Phase 2 (fusion).* After concatenating $\{\mathbf{z}^{(s,m)}\}$ and encoding to a joint representation $\mathbf{z}^{(s)}$, optimize

$$\mathcal{L}^{(2)} = \mathcal{L}_{\text{rec}}^{\text{central}} + \lambda_{\text{mmd}}\,\text{MMD}\big(q(z)\,\|\,p(z)\big) + \alpha\,\mathcal{L}_{\text{CE}}^{\text{central}}$$

where Maximum Mean Discrepancy (MMD) penalizes deviation of the aggregated posterior $q(z)$ from a prior $p(z)$ (Gaussian kernel).

**Biomarker identification.** After training, compute SHapley Additive exPlanations (SHAP)[20] values $\phi_g^{(m)}(\mathbf{x}^{(s,m)})$ per feature $g$ and sample $s$ within each omic $m$, and aggregate across a specific sample set to obtain global importance

$$I_g^{(m)} = \frac{1}{|S|}\sum_{s\in S}\big|\phi_g^{(m)}(\mathbf{x}^{(s,m)})\big|,$$

##### 1.6.5 MOGONET

Wang *et al.*[21] proposed MOGONET, which combines omics-specific graph convolutional networks (GCNs) with label-space late integration via a View Correlation Discovery Network (VCDN). Each omics type is modeled on its own sample-sample graph to produce class probabilities, and then these per-omic label distributions are fused at a higher-order tensor level for final predictions.

**Inputs.** Let $\{\mathbf{X}^{(i)}\}_{i=1}^{m}$ be multi-omics feature matrices with $\mathbf{X}^{(i)} \in \mathbb{R}^{N\times P_i}$ ($N$ samples, $P_i$ features). To build sample-sample graphs, for each omic $i$, a weighted adjacency matrix $A^{(i)} \in \mathbb{R}^{N\times N}$ is computed using cosine similarity. An adaptive threshold is chosen so the average node degree approximates a tuned hyperparameter $k$.

**Architecture.** The inputs $(\mathbf{X}^{(i)}, A^{(i)})$ are fed into an $L$-layer GCN to obtain per-omic class probabilities

$$\widehat{\mathbf{Y}}^{(i)} \;=\; \text{GCN}^{(i)}\big(\mathbf{X}^{(i)}, A^{(i)}\big) \in \mathbb{R}^{N\times C}.$$

Each GCN is trained with class-weighted cross-entropy loss.

For label-space integration, let $\widehat{\mathbf{y}}_n^{(i)} \in \mathbb{R}^{C}$ be the per-omic distribution for sample $n$. Construct the cross-omics discovery tensor by the outer-product of per-omic distributions:

$$\mathcal{C}_n[a_1, \ldots, a_m] \;=\; \prod_{i=1}^{m} \widehat{\mathbf{y}}^{(i)}_{n,a_i}, \qquad a_i \in \{1, \ldots, C\}.$$

Then, flatten $\mathbf{c}_n = \mathrm{vec}(\mathcal{C}_n) \in \mathbb{R}^{C^m}$ and pass to a fully connected network (VCDN head):

$$\mathbf{z}_n \;=\; \mathrm{VCDN}(\mathbf{c}_n) \in \mathbb{R}^C, \qquad \widehat{\mathbf{y}}_n \;=\; \mathrm{softmax}(\mathbf{z}_n).$$

**Training objectives.** The training objective for the VCDN head is a cross-entropy loss

$$\mathcal{L}_{VCDN} = \sum_{n=1}^{N} \mathcal{L}_{CE}(\widehat{\mathbf{y}}_n, \mathbf{y}_n)$$

and the total loss is

$$\mathcal{L} \;=\; \sum_{i=1}^{m} \mathcal{L}^{(i)}_{\mathrm{GCN}} \;+\; \gamma\, \mathcal{L}_{\mathrm{VCDN}}$$

The optimization process adopts an alternating training scheme. Specifically, after each GCN is pretrained, the procedure alternates (i) updating all GCNs with VCDN fixed and (ii) updating VCDN with GCNs fixed, until convergence. At test time, a new sample is linked to training nodes via cosine similarity to form the augmented graph for each omic, then forwarded through the trained GCNs and VCDN.

**Biomarker identification.** MOGONET uses the feature ablation technique for scoring the importance of features, specifically by zeroing an individual feature and measuring the performance metric drop in the test set. After training, for omic $i$ and feature $f$:

$$\Delta M^{(i)}(f) \;=\; M(\text{original feature values}) - M(\text{feature } f \text{ set to } 0),$$

using F1 for binary classification tasks.

##### 1.6.6 MoAGL-SA

Cheng *et al.*[22] proposed MoAGL-SA, an adaptive multi-omics integration framework that learns per-omic sample graphs directly from data, extracts omic-specific graph embeddings with GCNs, and fuses them via self-attention for downstream tasks.

**Inputs.** Let $\{\mathbf{X}^{(m)}\}_{m=1}^{M}$ be matched omics with $\mathbf{X}^{(m)} \in \mathbb{R}^{N \times D_m}$. For each omic $m$, an adjacency $\mathbf{A}^{(m)}$ is learned to reflect sample similarity. Nearby samples receive larger edge weights, and a small penalty induces sparsity. Self-loops are added and the adjacency is normalized with the standard GCN normalization.

**Architecture.** A GCN run on $(\mathbf{X}^{(m)}, \mathbf{A}^{(m)})$ produces per-omic embeddings

$$\mathbf{Z}^{(m)} = \mathrm{GCN}\big(\mathbf{X}^{(m)},\, \tilde{\mathbf{A}}^{(m)}; \Theta^{(m)}\big),$$

and the per-sample, per-omic vector is $\mathbf{z}^{(s,m)}$ (row $s$ of $\mathbf{Z}^{(m)}$). For each sample $s$, compute dot-product self-attention over omics:

$$\bar{\mathbf{z}}^{(s)} = \tfrac{1}{M} \sum_{m=1}^{M} \mathbf{z}^{(s,m)}, \quad \mathbf{q}^{(s)} = \mathbf{W}_Q\, \bar{\mathbf{z}}^{(s)}, \quad \mathbf{k}^{(s,m)} = \mathbf{W}_K\, \mathbf{z}^{(s,m)}, \quad \mathbf{v}^{(s,m)} = \big[g_{\mathrm{enc}}(\tilde{\mathbf{A}}^{(m)}, \mathbf{Z}^{(m)})\big]_s$$

with $g_{\mathrm{enc}}(\tilde{\mathbf{A}}^{(m)}, \mathbf{Z}^{(m)}) = \tilde{\mathbf{A}}^{(m)} \mathbf{Z}^{(m)} \mathbf{W}_V$ being a graph encoder.

$$e^{(s)}_m = \frac{\big(\mathbf{q}^{(s)}\big)^{\top} \mathbf{k}^{(s,m)}}{\sqrt{d_k}}, \qquad a^{(s)}_m = \frac{\exp\big(e^{(s)}_m\big)}{\sum_{n=1}^{M} \exp\big(e^{(s)}_n\big)}, \qquad \mathbf{z}^{(s)} = \sum_{m=1}^{M} a^{(s)}_m\, \mathbf{v}^{(s,m)}.$$

where $\mathbf{W}_Q \in \mathbb{R}^{d_k \times d}$, $\mathbf{W}_K \in \mathbb{R}^{d_k \times d}$, $\mathbf{W}_V \in \mathbb{R}^{d_v \times d}$ are learned projections; $\mathbf{z}^{(s,m)} \in \mathbb{R}^d$ is the embedding for the $m$ omics type; $d_k$ is the key/query dimension; and the softmax is taken over $m = 1, \ldots, M$.

**Training objectives.** A classifier follows

$$\mathbf{o}^{(s)} = f_{\mathrm{clf}}\big(\mathbf{z}^{(s)}; \theta_{\mathrm{clf}}\big), \qquad \hat{\mathbf{p}}^{(s)} = \mathrm{softmax}(\mathbf{o}^{(s)}),$$

where $f_{\text{clf}}$ is a feed-forward network with residual and normalization. Then, a decoder maps $\mathbf{Z}' = \sum_{m=1}^{M} \mathbf{Z}^{(m)}$ back to each omic embedding

$$\hat{\mathbf{Z}}^{(m)} = g_{\text{dec}}\big(\tilde{\mathbf{A}}^{(m)}, \mathbf{Z}'\big)$$

with a reconstruction loss which is Mean Squared Error (MSE) between $\mathbf{Z}^{(m)}$ and $\hat{\mathbf{Z}}^{(m)}$. The overall training objective for classification is

$$\mathcal{L} \;=\; -\frac{1}{|\mathcal{B}|}\sum_{s\in\mathcal{B}}\sum_{c=1}^{C} \mathbf{y}_{s,c}\log \hat{\mathbf{p}}_c^{(s)} \;+\; \alpha\sum_{m=1}^{M}\mathcal{L}_{\text{GL}}^{(m)} \;+\; \beta\,\mathcal{L}_{\text{rec}},$$

where $\mathbf{y}_{s,\cdot}$ is one-hot encoded labels, $\mathcal{L}_{GL}^{m}$ is the omic-specific graph-learning loss (with sparsity and locality constraint), and $\alpha, \beta \geq 0$ are weight parameters for the auxiliary terms.

**Biomarker identification.** MoAGL-SA uses feature ablation for feature importances. For feature $i$ in omic $m$, set its test values to zero and use a performance metric (e.g., accuracy) drop as the importance.

##### 1.6.7 MORE

Wang *et al.*[23] proposed MORE, a hypergraph-based multi-omics integration framework.

**Inputs.** Let $\{\mathbf{X}^{(m)}\}_{m=1}^{M}$ be the multi-omics features with $\mathbf{X}^{(m)} \in \mathbb{R}^{N\times P_m}$. MORE builds a $k$NN hypergraph per omic to obtain incidence matrices $\{\mathbf{H}^{(m)}\}_{m=1}^{M}$ and then forms a fused hypergraph by concatenation:

$$\mathbf{H} \;=\; \text{Concat}\big[\mathbf{H}^{(1)}, \ldots, \mathbf{H}^{(M)}\big].$$

**Architecture.** *Hypergraph convolution (MOHE).* Omics-specific embeddings are obtained with a standard hypergraph convolution using the fused hypergraph:

$$\mathbf{Z}^{(m)} \;=\; \sigma\Big(\mathbf{D}_v^{-\frac{1}{2}}\,\mathbf{H}\,\mathbf{W}_e\,\mathbf{D}_e^{-1}\,\mathbf{H}^{\top}\mathbf{D}_v^{-\frac{1}{2}}\,\mathbf{X}^{(m)}\,\boldsymbol{\Theta}^{(m)}\Big),$$

where $\mathbf{Z}^{(m)} \in \mathbb{R}^{N\times d}$, $\sigma(\cdot)$ is a nonlinearity, $\boldsymbol{\Theta}^{(m)}$ are learnable weights, $\mathbf{W}_e$ is a diagonal hyperedge-weight matrix, $\mathbf{D}_e = \text{diag}(\mathbf{H}^{\top}\mathbf{1})$ is the hyperedge-degree matrix, and $\mathbf{D}_v = \text{diag}(\mathbf{H}\mathbf{W}_e\mathbf{1})$ is the vertex-degree matrix. The per-sample, per-omic vector is $\mathbf{z}^{(s,m)}$ (row $s$ of $\mathbf{Z}^{(m)}$).

*Cross-omics integration (MOSA).* For each sample $s$, stack the omic-specific vectors into $\hat{\mathbf{X}}^{(s)} = [\,\mathbf{z}^{(s,1)} \;\cdots\; \mathbf{z}^{(s,M)}\,] \in \mathbb{R}^{d\times M}$ and compute self-attention:

$$\mathbf{Q}^{(s)} = \mathbf{W}_Q\hat{\mathbf{X}}^{(s)} + \mathbf{b}_Q, \quad \mathbf{K}^{(s)} = \mathbf{W}_K\hat{\mathbf{X}}^{(s)} + \mathbf{b}_K, \quad \mathbf{V}^{(s)} = \mathbf{W}_V\hat{\mathbf{X}}^{(s)} + \mathbf{b}_V,$$

$$\mathbf{A}^{(s)}(m,n) \;=\; \frac{\exp\big(\mathbf{q}^{(s,m)\top}\mathbf{k}^{(s,n)}/\sqrt{d_f}\big)}{\sum_{n'=1}^{M}\exp\big(\mathbf{q}^{(s,m)\top}\mathbf{k}^{(s,n')}/\sqrt{d_f}\big)}, \qquad \hat{\mathbf{V}}^{(s)} \;=\; \mathbf{A}^{(s)}\big(\mathbf{V}^{(s)}\big)^{\top},$$

where $\mathbf{W}_Q, \mathbf{W}_K, \mathbf{W}_V \in \mathbb{R}^{d_f\times d}$ and $\mathbf{b}_Q, \mathbf{b}_K, \mathbf{b}_V$ are learnable; $\mathbf{q}^{(s,m)}$ and $\mathbf{k}^{(s,n)}$ are the $m$-th and $n$-th columns of $\mathbf{Q}^{(s)}$ and $\mathbf{K}^{(s)}$; $d_f$ is the attention dimension. Then, take the integrated representation $\mathbf{z}^{(s)}$ as a linear projection or flattening of $\hat{\mathbf{V}}^{(s)}$. For multi-heads, the above repeats and concatenation is performed.

**Training objectives.** A linear softmax classifier takes as input $\mathbf{z}^{(s)}$:

$$\hat{\mathbf{p}}^{(s)} = \text{softmax}\big(\mathbf{W}\mathbf{z}^{(s)} + \mathbf{b}\big), \quad \mathbf{W} \in \mathbb{R}^{C\times d'}, \; \mathbf{b} \in \mathbb{R}^{C}.$$

First, MORE adopts a warmup step by training each MOHE branch with its own classifier on $\mathbf{z}^{(s,m)}$:

$$\mathcal{L}_{\text{omics-spec}}^{(m)} = \sum_{s=1}^{N}\mathcal{L}_{\text{CE}}\big(\hat{\mathbf{p}}^{(s,m)}, \mathbf{y}_s\big), \qquad \hat{\mathbf{p}}^{(s,m)} = \text{softmax}\big(\mathbf{W}^{(m)}\mathbf{z}^{(s,m)} + \mathbf{b}^{(m)}\big),$$

where $\mathbf{y}_s$ is one-hot encoded labels. Second, the sum of omic-specific and integrated losses is optimized:

$$\mathcal{L} \;=\; \sum_{m=1}^{M}\mathcal{L}_{\text{omics-spec}}^{(m)} \;+\; \sum_{s=1}^{N}\mathcal{L}_{\text{CE}}\big(\hat{\mathbf{p}}^{(s)}, \mathbf{y}_s\big).$$

**Biomarker identification.** MORE uses feature ablation for feature importances. For feature $i$ in omic

$m$, set its test-set values to zero and measure the performance drop (F1 for binary classification).

##### 1.6.8 MOGLAM

Ouyang *et al.*[24] proposed MOGLAM, a multi-omics integration framework that combines omic-specific adaptive graph learning and feature selection, multi-omics attention over omic embeddings, and omic-integrated representation learning for sample classification and biomarker identification.

**Inputs.** Let $\{\mathbf{X}^{(m)}\}_{m=1}^{M}$ be multi-omics feature matrices with $\mathbf{X}^{(m)} \in \mathbb{R}^{N \times P_m}$.

**Architecture.** MOGLAM has three main modules: an omic-specific dynamic GCN with feature selection (FSDGCN), a multi-omics attention mechanism (MOAM), and omic-integrated representation learning (OIRL).

*Omic-specific FSDGCN.* For omic $m$, a learnable cosine-similarity graph is built from sample features:

$$S_{ij}^{(m)} = \cos\big(\mathbf{W}_s^{(m)} \mathbf{x}_i^{(m)},\ \mathbf{W}_s^{(m)} \mathbf{x}_j^{(m)}\big),$$

with thresholds to keep only sufficiently similar neighbors. This learned graph is fused with an initial graph $\mathbf{A}^{(m)}$:

$$\hat{\mathbf{A}}^{(m)} = \eta\, \mathbf{A}^{(m)} + (1 - \eta)\, \mathbf{S}^{(m)},$$

then normalized with the standard GCN normalization to produce $\tilde{\mathbf{A}}^{(m)} = \hat{\mathbf{D}}^{-\frac{1}{2}} (\hat{\mathbf{A}}^{(m)} + \mathbf{I}) \hat{\mathbf{D}}^{-\frac{1}{2}}$, where $\hat{\mathbf{D}}$ is a diagonal matrix of $\hat{\mathbf{A}}^{(m)}$. A two-layer GCN on $\big(\mathbf{X}^{(m)}, \tilde{\mathbf{A}}^{(m)}\big)$ produces omic-specific embeddings $\mathbf{Z}^{(m)} \in \mathbb{R}^{N \times d_f}$. Specifically,

$$\mathbf{H}^{(1,m)} = \mathrm{GCN}\big(\mathbf{H}^{(0,m)}, \tilde{\mathbf{A}}^{(m)}; \Theta^{(0,m)}\big)$$

$$\mathbf{Z}^{(m)} = \mathrm{GCN}\big(\mathbf{H}^{(1,m)}, \tilde{\mathbf{A}}^{(m)}; \Theta^{(1,m)}\big)$$

Feature selection is realized via an omic-specific projection/indicator matrix $\mathbf{W}_f^{(m)} \in \mathbb{R}^{P_m \times d_f}$ and a regularizer

$$\Omega\big(\mathbf{W}_f^{(m)}\big) = \big\|\mathbf{W}_f^{(m)} \mathbf{W}_f^{(m)\top}\big\|_1 - \big\|\mathbf{W}_f^{(m)}\big\|_F^2,$$

which encourages correlated features to share large scores, and $\mathbf{H}^{(0,m)} = \mathbf{X}^{(m)} \mathbf{W}_f^{(m)}$.

*Multi-omics attention (MOAM).* For each omic $m$, MOGLAM computes a scalar summary $g_m$ by averaging all entries of $\mathbf{Z}^{(m)}$, and forms $\mathbf{g} = [g_1, \ldots, g_M]^\top$. A small gating network produces omic weights

$$\mathbf{g}_{\text{att}} = \sigma\big(\mathbf{W}_2\, \mathrm{ReLU}(\mathbf{W}_1 \mathbf{g})\big),$$

where $\sigma$ is the sigmoid function and $\mathbf{W}_1, \mathbf{W}_2$ are learnable weights. The attended omic embeddings are

$$\tilde{\mathbf{Z}}^{(m)} = g_{\text{att}}^{(m)}\, \mathbf{Z}^{(m)},$$

and per-sample vectors $\tilde{\mathbf{z}}^{(s,m)}$ are stacked across $m$.

*Omic-integrated representation learning (OIRL).* For each sample $s$, the set $\{\tilde{\mathbf{z}}^{(s,1)}, \ldots, \tilde{\mathbf{z}}^{(s,M)}\}$ is treated as $M$ "omic tokens" and processed by multi-head scaled dot-product self-attention over omics, followed by a small feedforward network. This yields a common representation $\mathbf{h}_{\text{cm}}^{(s)}$ and a complementary representation $\mathbf{h}_{\text{cp}}^{(s)}$ (constructed from the attention weights), which are concatenated:

$$\mathbf{h}^{(s)} = \big[\mathbf{h}_{\text{cm}}^{(s)}; \mathbf{h}_{\text{cp}}^{(s)}\big],$$

and fed into an MLP classifier:

$$\hat{\mathbf{Y}}_{\text{MLP}} = \mathrm{softmax}\big(f_{\text{MLP}}(\mathbf{H}; \theta_{\text{MLP}})\big).$$

where $\hat{\mathbf{Y}}_{\text{MLP}} \in \mathbb{R}^{N \times C}$ is the omic-integrated class probabilities over $N$ samples.

**Training objectives.** Each omic-specific FSDGCN branch has a loss

$$\mathcal{L}_{\text{FSDGCN}}^{(m)} = \mathcal{L}_{\text{CE}}\big(\mathbf{Y}, \hat{\mathbf{Y}}_{\text{FSDGCN}}^{(m)}\big) + \alpha\, \mathcal{L}_{\text{gl}}^{(m)} + \beta\, \Omega\big(\mathbf{W}_f^{(m)}\big),$$

where $\mathcal{L}_{\text{CE}}$ is cross-entropy on omic-specific predictions, $\mathcal{L}_{\text{gl}}^{(m)}$ is the graph-learning penalty (encouraging smoothness and proximity to the initial graph), $\alpha, \beta > 0$ are trade-off parameters, and $\hat{\mathbf{Y}}_{\text{FSDGCN}}^{(m)} \in \mathbb{R}^{N \times C}$

denotes the class-probability outputs of the $m$-th FSDGCN branch. The final objective is:

$$\mathcal{L} = \sum_{m=1}^{M} \mathcal{L}_{\mathrm{FSDGCN}}^{(m)} + \mu\, \mathcal{L}_{\mathrm{CE}}\big(\mathbf{Y}, \hat{\mathbf{Y}}_{\mathrm{MLP}}\big),$$

where $\hat{\mathbf{Y}}_{\mathrm{MLP}}$ comes from the omic-integrated representation and $\mu > 0$ balances omic-specific and integrated losses. Training first pretrains each FSDGCN branch, then alternates between updating omic-specific FSDGCNs (with MOAM/OIRL fixed) and updating MOAM and OIRL (with FSDGCNs fixed) until convergence.

**Biomarker identification.** Feature importance is derived from the feature-selection matrices $\mathbf{W}_f^{(m)}$. For feature $i$ in omic $m$, its importance is calculated as the mean of the absolute values of the corresponding row $\mathbf{w}_{f,i}^{(m)} \in \mathbb{R}^{d_f}$.

##### 1.6.9 GNN-SubNet

Pfeifer *et al.*[25] proposed GNN-SubNet, a graph-based framework for disease subnetwork detection that represents each patient as a protein-protein interaction (PPI) graph with multi-omics node features. It performs graph-level classification with a Graph Isomorphism Network (GIN)[26], and adapted GNNExplainer[27] to obtain model-wide node/edge importances for subnetwork discovery.

**Inputs.** Let $\mathbf{A} \in \mathbb{R}^{|\mathcal{V}| \times |\mathcal{V}|}$ be the fixed adjacency of a PPI network and $\mathbf{X}^{(s)} \in \mathbb{R}^{|\mathcal{V}| \times P}$ the node-feature matrix for patient $s$ (rows are genes, columns are omics features such as mRNA expression and DNA methylation). Each patient graph is $(\mathbf{A}, \mathbf{X}^{(s)})$, with shared topology.

**Architecture.** GNN-SubNet uses a standard GIN for graph classification. Let $\mathbf{h}_v^{(0)}$ be the initial feature of node $v$ (row of $\mathbf{X}^{(s)}$). At layer $k$,

$$\mathbf{h}_v^{(k)} = \mathrm{MLP}^{(k)}\Big(\big(1 + \epsilon^{(k)}\big)\, \mathbf{h}_v^{(k-1)} + \sum_{u \in \mathcal{N}(v)} \mathbf{h}_u^{(k-1)}\Big),$$

where $\epsilon^{(k)}$ and the MLP parameters are learned. After each GIN layer, global sum pooling is applied and the pooled representations are concatenated into a graph embedding $\mathbf{g}^{(s)}$, which is fed to fully connected layers and a final softmax classifier

$$\hat{\mathbf{p}}^{(s)} = \mathrm{softmax}\big(f_{\mathrm{clf}}(\mathbf{g}^{(s)}; \theta_{\mathrm{clf}})\big),$$

**Training objectives.** The GIN is trained with standard cross-entropy over graphs:

$$\mathcal{L}_{\mathrm{CE}} = -\frac{1}{|\mathcal{B}|} \sum_{s \in \mathcal{B}} \sum_{c=1}^{C} \mathbf{y}_{s,c} \,\log \hat{\mathbf{p}}_c^{(s)},$$

where $\mathbf{y}_{s,\cdot}$ is the one-hot encoded label for patient $s$ and $\mathcal{B}$ is a batch of graphs.

**Biomarker identification.** GNN-SubNet adapts GNNExplainer to learn a model-wide node-importance mask. Let $\mathbf{n} \in [0,1]^{|\mathcal{V}|}$ be a node mask shared across patients. For a given patient $s$, masked features are

$$\tilde{\mathbf{X}}^{(s)} = \mathrm{diag}(\mathbf{n})\, \mathbf{X}^{(s)},$$

and the classifier output on $(\mathbf{A}, \tilde{\mathbf{X}}^{(s)})$ is used in a loss that penalizes changes in the predicted label:

$$\mathcal{L}(\mathbf{n}) = -\log P_{\Theta}\big(Y = y^{(s)} \mid \mathbf{A}, \tilde{\mathbf{X}}^{(s)}\big),$$

By repeatedly sampling patients and updating the single mask $\mathbf{n}$, the optimization converges to global node scores $n_v$ indicating how important node $v$ is for the trained GIN.

##### 1.6.10 Pathformer

Liu *et al.*[5] proposed Pathformer, a transformer-based model utilizing biological pathway knowledge by mapping genes to pathway tokens via a biologically constrained sparse layer, and performs criss-cross attention to integrate pathway-pathway crosstalk and cross-modality information.

**Inputs.** Let $\mathcal{M}$ denote the set of omics types with $|\mathcal{M}| = m$. For modality $i \in \mathcal{M}$, let $\mathbf{X}^{(i)} \in \mathbb{R}^{N \times P_i}$ be the input matrix with $N$ samples and $P_i$ features. Let $\mathcal{G}$ be the reference gene set with $|\mathcal{G}| = G$ and let $\mathbf{R}^{(i)} \in \{0,1\}^{P_i \times G}$ be a feature-gene mapping matrix (entry 1 iff a feature maps to a gene). Each modality is converted to gene-level features via a fixed set of summary operators (e.g. count/mean/min/max/entropy/weighted means) applied over features mapped to each gene:

$$\mathbf{G}^{(i)} = \Phi^{(i)}\big(\mathbf{X}^{(i)}, \mathbf{R}^{(i)}\big) \in \mathbb{R}^{N \times G \times d_i},$$

followed by a linear projection to a shared width $d$:

$$\tilde{\mathbf{G}}^{(i)} = \mathbf{G}^{(i)}\mathbf{W}^{(i)} \in \mathbb{R}^{N \times G \times d}, \qquad \mathbf{W}^{(i)} \in \mathbb{R}^{d_i \times d}.$$

Missing modalities are filled with zero at the gene level.

**Architecture.** *Gene-to-pathway encoder.* Let $\mathcal{P}$ be the curated pathway set with $|\mathcal{P}| = p$, and let $\mathbf{M}_{\text{gp}} \in \{0,1\}^{G \times p}$ denote the gene-pathway incidence matrix. For each modality $i$, genes are mapped to pathway tokens by a pathway-sparse linear map

$$\mathbf{H}^{(i)} = \big(\mathbf{W}_{\text{gp}} \odot \mathbf{M}_{\text{gp}}\big)^{\top} \tilde{\mathbf{G}}^{(i)} + \mathbf{1}\,\mathbf{b}^{\top} \in \mathbb{R}^{N \times p \times d},$$

where $\mathbf{W}_{\text{gp}} \in \mathbb{R}^{G \times p}$ is trainable only where $\mathbf{M}_{\text{gp}} = 1$ ($\odot$ is the Hadamard product), $\mathbf{b} \in \mathbb{R}^{d \times 1}$ is a bias shared across modalities, and $\mathbf{1} \in \mathbb{R}^{N \times 1}$ is a vector of ones.

*Criss-cross pathway-modality attention.* Stack $\{\mathbf{H}^{(i)}\}_{i=1}^{m}$ along the modality axis to obtain $\mathbf{T} \in \mathbb{R}^{N \times m \times p \times d}$. Define modality-averaged pathway tokens

$$\bar{\mathbf{H}} = \frac{1}{m}\sum_{i=1}^{m} \mathbf{H}^{(i)} \in \mathbb{R}^{N \times p \times d},$$

and let $\mathbf{P} \in \mathbb{R}^{p \times p}$ be a weighted pathway crosstalk matrix computed by BinoX [28] (significance or effect size of crosstalk).

For each sample (sample index omitted here), pathway self-attention with crosstalk bias is

$$\mathbf{Q} = \bar{\mathbf{H}}\mathbf{W}_q, \quad \mathbf{K} = \bar{\mathbf{H}}\mathbf{W}_k, \quad \mathbf{V} = \bar{\mathbf{H}}\mathbf{W}_v, \qquad \mathbf{U}_{\text{path}} = \operatorname{softmax}\!\left(\frac{\mathbf{Q}\mathbf{K}^{\top}}{\sqrt{d_k}} + \beta\,\mathbf{P}\right)\mathbf{V},$$

where $\mathbf{W}_q, \mathbf{W}_k, \mathbf{W}_v \in \mathbb{R}^{d \times d_k}$ are projections and $\beta$ controls the crosstalk prior strength.

For modality self-attention, for each pathway $t$ consider $\mathbf{T}_{:,t} \in \mathbb{R}^{m \times d}$ and compute

$$\mathbf{Q}_{\text{mod}} = \mathbf{T}_{:,t}\mathbf{W}_q^{\text{mod}}, \quad \mathbf{K}_{\text{mod}} = \mathbf{T}_{:,t}\mathbf{W}_k^{\text{mod}}, \quad \mathbf{V}_{\text{mod}} = \mathbf{T}_{:,t}\mathbf{W}_v^{\text{mod}},$$

$$\mathbf{U}_{\text{mod},t} = \operatorname{softmax}\!\left(\frac{\mathbf{Q}_{\text{mod}}\mathbf{K}_{\text{mod}}^{\top}}{\sqrt{d_k}}\right)\mathbf{V}_{\text{mod}} \in \mathbb{R}^{m \times d},$$

with $\mathbf{W}_q^{\text{mod}}, \mathbf{W}_k^{\text{mod}}, \mathbf{W}_v^{\text{mod}} \in \mathbb{R}^{d \times d_k}$. Residual connections, LayerNorm, and an MLP fuse $\mathbf{U}_{\text{path}}$ and $\{\mathbf{U}_{\text{mod},t}\}_{t=1}^{p}$ to yield modality-fused pathway tokens $\mathbf{Z} \in \mathbb{R}^{N \times p \times d}$. Between blocks, $\mathbf{P}$ can be updated from the current pathway embeddings (e.g. correlation-based), allowing the crosstalk prior to be refined.

**Training objective.** Flatten pathway tokens per sample and apply an MLP:

$$\hat{\mathbf{Y}} = \operatorname{MLP}\big(\operatorname{vec}(\mathbf{Z})\big) \in \mathbb{R}^{N \times C}, \qquad \operatorname{vec}(\mathbf{Z}) \in \mathbb{R}^{N \times (pd)}.$$

For classification, Pathformer is trained with cross-entropy

$$\mathcal{L}_{\text{cls}} = -\frac{1}{N}\sum_{i=1}^{N}\sum_{c=1}^{C} \mathbf{y}_{i,c}\log\hat{\mathbf{Y}}_{i,c},$$

where $\mathbf{y}_{i,\cdot}$ is the one-hot encoded label of sample $i$.

**Biomarker identification.** Gene-level importance is obtained by SHAP [20] with respect to gene-level inputs. For modality $i$, let $\mathbf{S}^{(i)} \in \mathbb{R}^{N \times G}$ be SHAP attributions (genes absent in modality $i$ are filled with zero values). The global score for gene $g$ aggregates absolute attributions across classes, samples, and modalities:

$$s_g = \frac{1}{C}\sum_{c=1}^{C}\frac{1}{N_c}\sum_{n\in\mathcal{I}_c}\sum_{i=1}^{m}|\mathbf{S}^{(i)}_{n,g}|,$$

where $N_c$ is the number of samples in class $c$ and $\mathcal{I}_c$ are sample indices.

##### 1.6.11 DeePathNet

Cai *et al.*[6] proposed DeePathNet, a transformer-based model that utilizes biological pathways for multi-omics integration. The model first encodes gene-level multi-omics features into pathway-level vectors, then applies transformer encoders to learn pathway-pathway interactions.

**Inputs.** Let $\mathcal{M}$ denote the set of omics types ($|\mathcal{M}| = m$). For pathway index $p \in \{1, \ldots, P\}$ with gene set $G_p$, define the pathway-level feature matrix by concatenating per-gene features across all available omics types (missing omics types are filled with zeros):

$$\mathbf{A}^{(p)}_{\text{omics}} \in \mathbb{R}^{N\times(m\,|G_p|)},$$

where $N$ is the number of samples.

**Architecture.** *Pathway encoder.* Each pathway is projected to a $d$-dimensional embedding via a fully connected layer:

$$\mathbf{A}^{(p)}_{\text{enc}} \;=\; \mathbf{A}^{(p)}_{\text{omics}}\,\mathbf{W}_p^{\top} \;+\; \mathbf{1}_N\,\mathbf{b}_p^{\top},$$

where $\mathbf{W}_p \in \mathbb{R}^{d\times(m\,|G_p|)}$ and $\mathbf{b}_p \in \mathbb{R}^d$ are learnable parameters, and $\mathbf{1}_N \in \mathbb{R}^{N\times 1}$ is the all-ones column vector. Stacking all $P$ pathway tokens yields

$$\mathbf{A}^{(0)} \;=\; \big[\mathbf{A}^{(1)}_{\text{enc}}, \ldots, \mathbf{A}^{(P)}_{\text{enc}}\big] \;\in\; \mathbb{R}^{N\times P\times d},$$

where $P$ is the number of pathways ($P = 241$ from LCPathways[13]).

*Transformer over pathways.* A dropout layer (probability 0.5) followed by standard transformer-encoder blocks (LayerNorm, multi-head self-attention, MLP with residual connections) is applied across the $P$ pathway tokens:

$$\mathbf{A}^{(1)} \;=\; \text{Transformer}\big(\mathbf{A}^{(0)}\big),$$
$$\mathbf{A}^{(2)} \;=\; \text{Transformer}\big(\mathbf{A}^{(1)}\big).$$

**Training objectives.** The final pathway representations are flattened and passed to an MLP:

$$\mathbf{Z} \;=\; \text{MLP}\big(\text{vec}(\mathbf{A}^{(2)})\big), \qquad \text{vec}(\mathbf{A}^{(2)}) \in \mathbb{R}^{N\times(Pd)},$$

where $\mathbf{Z} \in \mathbb{R}^{N\times C}$ are class logits for $C$ classes. Define predicted class probabilities and ground-truth labels as:

$$\widehat{\mathbf{Y}} \;=\; \text{softmax}(\mathbf{Z}) \in \mathbb{R}^{N\times C}, \qquad \mathbf{Y} \in \{0,1\}^{N\times C}$$

The training objective is:

$$\mathcal{L}_{\text{CE}} \;=\; -\frac{1}{N}\sum_{i=1}^{N}\sum_{c=1}^{C}\mathbf{Y}_{ic}\,\log\big(\widehat{\mathbf{Y}}_{ic}\big)$$

**Biomarker identification.** DeePathNet uses SHAP[20] to attribute per-modality features. Given multi-omics input data $\mathbf{X}^{(1)}, ..., \mathbf{X}^{(m)}$, where $\mathbf{X}^{(i)} \in \mathbb{R}^{N\times P_i}$ is the omics data matrix with $N$ samples and $P_i$ features for the $i$-th omics type, SHAP attribution produces feature scores $\mathbf{S}^{(1)}, ..., \mathbf{S}^{(m)}$, where $\mathbf{S}^{(i)} \in \mathbb{R}^{N\times P_i}$ is the score matrix for the $i$-th omics type. The final score for the $j$-th feature in the $i$-th omics type $s^{(i)}_j$ is calculated by

$$s^{(i)}_j = \frac{1}{C}\sum_{c=1}^{C}\frac{1}{N_c}\sum_{n=1}^{N_c}|\mathbf{S}^{(i)}_{n,j}|$$

where $C$ is the number of classes, and $N_c$ denotes the number of samples in class $c$. The model's predictions can be explained by top-scored features and those features are interpreted as potential biomarkers.

###### 1.6.12 DeepKEGG

Lan *et al.*[29] proposed DeepKEGG, a multi-omics method that maps genes/miRNAs to KEGG[30] pathways via a biologically constrained hierarchical module. It then learns sample-sample correlations through a pathway-level self-attention block for each omics type, and performs predictions using an MLP over the concatenated output embeddings of each block.

**Inputs.** Let $\mathbf{X}^{(i)}, ..., \mathbf{X}^{(m)}$ be the input multi-omics data, with $\mathbf{X}^{(i)} \in \mathbb{R}^{N \times P_i}$ being the feature matrix with $N$ samples and $P_i$ features for the $i$-th omics type. Let $\mathcal{P}$ be the KEGG pathway set. Construct a modality-specific feature-pathway incidence matrix $\mathbf{M}^{(i)} \in \{0,1\}^{P_i \times |\mathcal{P}|}$ ($\mathbf{M}^{(i)}_{f,t} = 1$ iff. pathway $p$ contains feature $f$).

**Architecture.** *Biological hierarchical module.* Pathway features are propagated from omics features via

$$\mathbf{H}^{(i)} \;=\; h\Big(\mathbf{X}^{(i)}\mathbf{M}^{(i)} \;+\; \mathbf{1}\,\mathbf{b}^{(i)\top}\Big)$$

where $\mathbf{H}^{(i)} \in \mathbb{R}^{N \times |\mathcal{P}|}$ are the pathway features, $h(\cdot)$ is an elementwise nonlinearity, $\mathbf{b}^{(i)} \in \mathbb{R}^{|\mathcal{P}| \times 1}$, and $\mathbf{1}_{|\mathcal{P}| \times 1}$ is a vector of ones.

*Sample-wise self-attention.* For each omics type $i$, pathway features are mapped to queries/keys/values via linear transformations:

$$\mathbf{Q}^{(i)} = \mathbf{H}^{(i)}\mathbf{W}^{(i)}_q, \quad \mathbf{K}^{(i)} = \mathbf{H}^{(i)}\mathbf{W}^{(i)}_k, \quad \mathbf{V}^{(i)} = \mathbf{H}^{(i)}\mathbf{W}^{(i)}_v,$$

with $\mathbf{W}^{(i)}_q, \mathbf{W}^{(i)}_k, \mathbf{W}^{(i)}_v \in \mathbb{R}^{|\mathcal{P}| \times d_k}$ and $\mathbf{Q}^{(i)}, \mathbf{K}^{(i)}, \mathbf{V}^{(i)} \in \mathbb{R}^{N \times d_k}$. For samples $a, b$,

$$\alpha^{(i)}_{ab} \;=\; \frac{\mathbf{Q}^{(i)}_a\,\mathbf{K}^{(i)\top}_b}{\sqrt{d_k}}, \qquad w^{(i)}_{ab} \;=\; \mathrm{softmax}_b\big(\alpha^{(i)}_{ab}\big),$$

and the final feature matrix for the $i$-th omics type is calculated as: $\mathbf{B}^{(i)} \in \mathbb{R}^{N \times d_k}$, where

$$\mathbf{B}^{(i)}_a \;=\; \textstyle\sum_b w^{(i)}_{ab}\,\mathbf{V}^{(i)}_b$$

**Training objectives.** Finally, feature matrices from all modalities are concatenated

$$\mathbf{Z} \;=\; \mathrm{concat}\Big(\mathbf{B}^{(1)}, \,...,\, \mathbf{B}^{(m)}\Big) \in \mathbb{R}^{N \times (md_k)}.$$

and an MLP is applied for the final output

$$\hat{\mathbf{y}} \;=\; \mathrm{MLP}(\mathbf{Z}) \in (0,1)^N.$$

using class-weighted cross-entropy with $\ell_2$ regularization (for binary classification):

$$\mathcal{L}_{\mathrm{CE}} = -\frac{1}{N}\sum_{n=1}^{N}\Big[w_1\,\mathbf{y}_n \log \hat{\mathbf{y}}_n \;+\; w_0\,(1-\mathbf{y}_n)\log(1-\hat{\mathbf{y}}_n)\Big] \;+\; \gamma\,\|\boldsymbol{\theta}\|_2^2,$$

with $w_j = \frac{N}{C\,N_c}$ for $j \in \{0,1\}$, where $C$ is the number of classes ($C = 2$ for binary classification), $N_c$ is the number of samples in class $c$, and $\boldsymbol{\theta}$ denotes all trainable parameters.

**Biomarker identification.** DeepKEGG adopts DeepLIFT[16] to compute per-feature attributions. For omics type $i$, let $\mathbf{S}^{(i)} \in \mathbb{R}^{N \times P_i}$ be attribution scores. For a particular sample $n$, its importance score of feature $j$ in the $i$-th modality is calculated as

$$\mathbf{S}^{(i)}_{n,j} = \frac{\mathrm{d}\hat{\mathbf{y}}_n}{\mathrm{d}\mathbf{X}^{(i)}_{n,j}}\Delta\mathbf{T}^i_{n,j}$$

where $\frac{\mathrm{d}\hat{\mathbf{y}}_n}{\mathrm{d}\mathbf{X}^{(i)}_{n,j}}$ represents the gradient of the predicted output with respect to an input feature, and $\Delta\mathbf{T}^i_{n,j} = (\mathbf{T}^i_{n,j} - \mathbf{T_{ref}}^i_{n,j})$ represents the difference between the actual activation of current input and the reference activation. The reference activation values are set to 0. Finally, the importance score of feature $j$ in the $i$-th omics type is

$$s_j^{(i)} = \sum_{n\in\mathcal{I}_+} \mathbf{S}_{n,j}^{(i)}$$

where $\mathcal{I}_+$ denotes the positive class.

###### 1.6.13 MCIA

Meng *et al.*[31] proposed Multiple Co-Inertia Analysis (MCIA), a multivariate integration model that jointly projects several high-dimensional omics datasets into a common low-dimensional space by maximizing their shared covariance structure.

**Inputs.** Let $\{\mathbf{M}^{(k)}\}_{k=1}^{K}$ be $K$ omics feature tables measured on the same $N$ samples, with $\mathbf{M}^{(k)} \in \mathbb{R}^{N\times P_k}$. Each table is first transformed by a one-table ordination method (e.g., PCA, correspondence analysis, or non-symmetric correspondence analysis) into a centered, scaled space:

$$\mathbf{X}^{(k)} \leftarrow \text{Ordinate}\big(\mathbf{M}^{(k)}\big),$$

thus placing heterogeneous omics on a comparable metric while preserving their major variance structure.

**Model.** MCIA generalizes co-inertia analysis to $K$ tables. For each transformed table $\mathbf{X}^{(k)}$, let $\mathbf{Q}^{(k)} \in \mathbb{R}^{P_k\times P_k}$ be a feature metric (typically diagonal) and let $\mathbf{D} \in \mathbb{R}^{N\times N}$ be a sample metric (often $\mathbf{D} = \mathbf{I}_N$). MCIA seeks table-specific axes $\{\mathbf{u}_k\}$ and a consensus sample axis $\mathbf{v}$ that maximize the sum of squared covariances between table scores and the synthetic axis:

$$\max_{\{\mathbf{u}_k\},\,\mathbf{v}} \sum_{k=1}^{K} w_k \operatorname{cov}^2\Big(\mathbf{X}^{(k)}\mathbf{Q}^{(k)}\mathbf{u}_k,\ \mathbf{v}\Big) \quad \text{s.t.} \quad \mathbf{v}^\top\mathbf{D}\mathbf{v} = 1, \ \ \mathbf{u}_k^\top\mathbf{Q}^{(k)}\mathbf{u}_k = 1,$$

where $w_k > 0$ are table weights. Concatenating the weighted tables

$$\mathbf{X} = \big[\sqrt{w_1}\,\mathbf{X}^{(1)} \,\|\, \cdots \,\|\, \sqrt{w_K}\,\mathbf{X}^{(K)}\big], \qquad \mathbf{Q} = \operatorname{diag}\big(\mathbf{Q}^{(1)}, \ldots, \mathbf{Q}^{(K)}\big),$$

the first MCIA axis $\mathbf{v}_1$ is obtained as the leading eigenvector of

$$\mathbf{X}\mathbf{Q}\mathbf{X}^\top\mathbf{D}\,\mathbf{v}_1 = \lambda_1\,\mathbf{v}_1,$$

and table-specific axes follow as

$$\mathbf{u}_k^{(1)} \propto \mathbf{X}^{(k)\top}\mathbf{D}\mathbf{v}_1, \qquad k = 1, \ldots, K.$$

Higher-order axes $\big\{\mathbf{v}_\ell, \mathbf{u}_k^{(\ell)}\big\}_{\ell\geq 2}$ are obtained sequentially on deflated residuals, with $\mathbf{v}_\ell^\top\mathbf{D}\mathbf{v}_{\ell'} = 0$ and $\big(\mathbf{u}_k^{(\ell)}\big)^\top\mathbf{Q}^{(k)}\mathbf{u}_k^{(\ell')} = 0$ for $\ell' < \ell$. Sample scores for table $k$ on axis $\ell$ are

$$\mathbf{t}_\ell^{(k)} = \mathbf{X}^{(k)}\mathbf{Q}^{(k)}\mathbf{u}_k^{(\ell)} \in \mathbb{R}^N,$$

Feature coordinates (loadings) for table $k$ on axis $\ell$ are $\mathbf{a}_\ell^{(k)} = \mathbf{Q}^{(k)}\mathbf{u}_k^{(\ell)} \in \mathbb{R}^{P_k}$, whose $j$-th entry $a_{j\ell}^{(k)}$ is the coordinate of feature $j$.

**Biomarker identification.** MCIA is unsupervised, phenotypic or clinical labels (e.g., tissue type) are overlaid on the MCIA sample scores *a posteriori*, and axes that visibly separate the classes of interest are chosen (typically the first two). For a given dataset $k$ and MCIA axis $\ell$, the feature coordinates (loadings) are the entries of

$$\mathbf{a}_{\cdot,\ell}^{(k)} = \begin{bmatrix} a_{1\ell}^{(k)} \\ \vdots \\ a_{P_k\ell}^{(k)} \end{bmatrix},$$

where $a_{j\ell}^{(k)}$ is the coordinate of feature $j$ on axis $\ell$. Meng *et al.* interpret features at either end of an axis as markers of the samples projected in that direction. In practice, for each dataset $k$ and each selected axis $\ell$ (e.g. $\ell = 1, 2$), they use the magnitude of the axis-specific loading

$$I_{j,\ell}^{(k)} = \big|a_{j\ell}^{(k)}\big|$$

as an importance score, rank features by $I_{j,\ell}^{(k)}$ separately for the positive ($a_{j\ell}^{(k)} > 0$) and negative ($a_{j\ell}^{(k)} < 0$) ends of the axis, and select the top features from each end.

##### 1.6.14 MOFA

Argelaguet *et al.*[32] proposed Multi-Omics Factor Analysis (MOFA), an unsupervised multi-omics factor model that decomposes several omics matrices into a shared low-dimensional representation, disentangling sources of variation that are shared across omics or specific to individual modalities.

**Inputs.** Let $\{\mathbf{Y}^{(m)}\}_{m=1}^{M}$ be $M$ omics views measured on $N$ samples, with $\mathbf{Y}^{(m)} \in \mathbb{R}^{N\times D_m}$. Different views can have different numbers of features $D_m$, and entire views can be missing for some samples.

**Model.** MOFA is a group factor analysis model. Each view $\mathbf{Y}^{(m)}$ is decomposed as

$$\mathbf{Y}^{(m)} = \mathbf{Z}\,\mathbf{W}^{(m)^\top} + \mathbf{E}^{(m)}, \qquad m = 1,\ldots,M,$$

where $\mathbf{Z} \in \mathbb{R}^{N\times K}$ is the factor matrix (rows are samples, columns are latent factors), $\mathbf{W}^{(m)} \in \mathbb{R}^{D_m\times K}$ is the loading matrix for view $m$, $\mathbf{E}^{(m)}$ is a view-specific residual noise term.

A standard normal prior is placed on the factors (rows of $\mathbf{Z}$). Each view uses its own likelihood: Gaussian for continuous data, Bernoulli for binary data, and Poisson for counts. For Gaussian views, residuals are modelled as $e_{nd}^{(m)} \sim \mathcal{N}(0, 1/\tau_d^{(m)})$ with feature-specific precisions $\tau_d^{(m)}$, whereas for Bernoulli and Poisson views the natural parameters are given by $(\mathbf{Z}\mathbf{W}^{(m)^\top})_{nd}$ via logistic or log-link functions.

To obtain interpretable factors with both view- and feature-specific sparsity, MOFA combines an automatic relevance determination (ARD) prior with a spike-and-slab decomposition of the loadings. Concretely, $w_{dk}^{(m)} = s_{dk}^{(m)}\,c_{dk}^{(m)}$, where $s_{dk}^{(m)} \sim \mathrm{Ber}(h_k^{(m)})$ encodes feature-wise sparsity for factor $k$ in view $m$ and $c_{dk}^{(m)} \sim \mathcal{N}(0, 1/\alpha_k^{(m)})$ has a view- and factor-specific precision $\alpha_k^{(m)}$ (ARD). Gamma priors on $\alpha_k^{(m)}$ and $h_k^{(m)}$ control which factors are active in which views and how many features load on each factor.

**Biomarker identification.** MOFA is unsupervised, biomarkers are derived from factors that align with clinically or biologically relevant axes. For a chosen factor $k$ and view $m$, the feature-level loadings are the $k$-th column of $\mathbf{W}^{(m)}$:

$$\mathbf{w}_{\cdot,k}^{(m)} = \begin{bmatrix} w_{1k}^{(m)} \\ \vdots \\ w_{D_m k}^{(m)} \end{bmatrix}.$$

MOFA inspects these loadings (typically in absolute value) and performs gene set enrichment on $\mathbf{w}_{\cdot,k}^{(m)}$ to annotate the factor. Genes with the largest absolute weights are reported as key drivers of that axis. Concretely, for view $m$ and factor $k$, a simple factor-specific importance score for feature (gene) $j$ is

$$I_{j,k}^{(m)} = \left|w_{jk}^{(m)}\right|,$$

and features are ranked by $I_{j,k}^{(m)}$ within that factor and view. Biomarker lists are then constructed by selecting the top-ranked genes for factors that show strong association with the clinical outcome or biological contrast of interest, optionally followed by gene set enrichment and manual curation, as done in the original MOFA analyses.

##### 1.6.15 GAUDI

Castellano-Escuder *et al.*[8] proposed GAUDI, a non-linear, unsupervised multi-omics integration method that embeds each omic with UMAP, clusters samples, and derives feature importance via XGBoost[33] (or random forest) and SHAP on the integrated latent space to explain sample clustering.

**Inputs.** Let $\{\mathbf{X}^{(m)}\}_{m=1}^{M}$ be multi-omics feature matrices with $\mathbf{X}^{(m)} \in \mathbb{R}^{N\times P_m}$.

**Model.** GAUDI proceeds in five steps:

*Independent UMAP embeddings.* Each omic is first embedded separately with UMAP to a low-dimensional space of fixed width $d$ (default $d$=4):

$$\mathbf{Z}^{(m)} = \mathrm{UMAP}\big(\mathbf{X}^{(m)}; \theta^{(m)}\big) \in \mathbb{R}^{N\times d}, \qquad m = 1,\ldots,M,$$

where $\theta^{(m)}$ are view-specific UMAP parameters (neighbors, metric, etc.). This standardizes omics to comparable latent dimensions regardless of $P_m$.

*Embedding concatenation and integrated UMAP.* The view-specific embeddings are concatenated:

$$\mathbf{Z} = \big[\mathbf{Z}^{(1)} \parallel \cdots \parallel \mathbf{Z}^{(M)}\big] \in \mathbb{R}^{N\times(Md)},$$

and a second UMAP produces an integrated latent space

$$\mathbf{Z}_{\text{int}} = \text{UMAP}\big(\mathbf{Z}; \theta_{\text{int}}\big) \in \mathbb{R}^{N\times d'}, \quad d' = 2,$$

which is used for visualization and downstream analysis.

*Density-based clustering.* A Hierarchical Density-Based Spatial Clustering (HDBSCAN) is applied on $\mathbf{Z}_{\text{int}}$ to obtain cluster labels

$$c_s = \text{HDBSCAN}\big(\mathbf{z}_{\text{int}}^{(s)}\big), \qquad s = 1, \ldots, N,$$

where $\mathbf{z}_{\text{int}}^{(s)}$ is row $s$ of $\mathbf{Z}_{\text{int}}$. This produces sample groups of varying density and shape without fixing the number of clusters.

*Surrogate models for metagenes.* To relate molecular features back to the integrated space, GAUDI trains XGBoost (or random forest) models that predict the integrated UMAP coordinates from each omic separately. For omic $m$ and latent dimension $r \in \{1, \ldots, d'\}$,

$$\hat{z}_{\text{int},r}^{(s)} = f_r^{(m)}\big(\mathbf{x}^{(s,m)}\big),$$

with $f_r^{(m)}$ a tree-boosting regressor fitted by minimizing squared error between $\hat{z}_{\text{int},r}^{(s)}$ and the true $z_{\text{int},r}^{(s)}$. This is done both for individual omics and for a combined feature set to define "metagenes" (latent factors summarized by a small set of influential features).

**Biomarker identification.** Feature importance is calculated using SHAP [20] on the trained surrogate models. For omic $m$, feature $j$, sample $s$, and latent dimension $r$, let

$$\phi_{s,j,r}^{(m)}$$

denote the SHAP value (contribution of feature $j$ to the prediction of $z_{\text{int},r}^{(s)}$ by $f_r^{(m)}$). In the original analyses, GAUDI selects top-ranked features per omic and per dimension as top genes contributing to GAUDI's latent space, then applies pathway enrichment (e.g. KEGG GSEA) and differential expression between GAUDI-defined clusters to interpret these signatures and report candidate biomarkers.

##### 1.6.16 DIABLO

Singh *et al.*[9] proposed Data Integration Analysis for Biomarker discovery using Latent cOmponents (DIABLO), a supervised multi-omics integrative method that extends sparse Generalized Canonical Correlation Analysis (sGCCA)[34] to jointly maximize shared covariance across omics datasets and a class-indicator matrix, and select a small multi-omics feature panel that discriminates phenotypic groups.

**Inputs.** Let $\{\mathbf{X}^{(q)}\}_{q=1}^{Q-1}$ be $Q-1$ omics datasets measured on the same $N$ samples, with $\mathbf{X}^{(q)} \in \mathbb{R}^{N\times P_q}$. Let $\mathbf{Y} \in \mathbb{R}^{N\times G}$ be the dummy indicator matrix encoding $G$ phenotype groups (columns sum to 1). Define $\{\mathbf{X}^{(q)}\}_{q=1}^{Q} := \{\mathbf{Y}, \mathbf{X}^{(1)}, \ldots, \mathbf{X}^{(Q-1)}\}$.

**Model.** DIABLO treats $\mathbf{Y}$ as an additional "omic" block and extends sGCCA to a supervised setting. Define a design matrix $\mathbf{C} \in [0,1]^{Q\times Q}$ specifying which blocks should be correlated (e.g. "full" vs. "null" designs). For component $h = 1, \ldots, H$, DIABLO estimates loading vectors $\{\mathbf{a}_h^{(q)}\}_{q=1}^{Q}$ by solving

$$\max_{\{\mathbf{a}_h^{(q)}\}} \sum_{i\neq j} c_{ij}\, \text{cov}\big(\mathbf{X}_h^{(i)}\mathbf{a}_h^{(i)}, \mathbf{X}_h^{(j)}\mathbf{a}_h^{(j)}\big) \quad \text{s.t.} \quad \big\|\mathbf{a}_h^{(q)}\big\|_2 = 1,\ \big\|\mathbf{a}_h^{(q)}\big\|_1 \le k^{(q)},$$

where $\mathbf{X}_h^{(q)}$ is the residual matrix for block $q$ at step $h$, $\mathbf{C} = (c_{ij})$ is the design, and $k^{(q)}$ controls sparsity. The latent components are

$$\mathbf{t}_h^{(q)} = \mathbf{X}_h^{(q)}\mathbf{a}_h^{(q)} \in \mathbb{R}^N,$$

**Training objectives.** Supervision is achieved by including $\mathbf{Y}$ as block $q = 1$ in the covariance objective above. In practice, $\mathbf{C}$ always connects each omic to $\mathbf{Y}$ so that the resulting latent components $\{\mathbf{t}_h^{(q)}\}$ are both inter-omics correlated and discriminative for the phenotype. For a new sample $s$, the block-$q$ component scores are

$$\mathbf{t}_{\text{new}}^{(q)} = \left(\mathbf{x}_{\text{new}}^{(q)}\right)^\top \mathbf{A}^{(q)} \in \mathbb{R}^H, \quad \mathbf{A}^{(q)} = \left[\mathbf{a}_1^{(q)}, \ldots, \mathbf{a}_H^{(q)}\right],$$

and class prediction for block $q$ is obtained by assigning $\mathbf{t}_{\text{new}}^{(q)}$ to a class using a distance measure in component space (e.g. centroid, maximum, or Mahalanobis distance), producing one predicted label per omic block. These per-omic predictions are then combined via majority vote or weighted vote (weights based on correlation between block-specific components and $\mathbf{Y}$) to produce a consensus class label.

**Biomarker identification.** Variable selection is achieved via the $\ell_1$ constraints on the loading vectors. For each block $q$ and component $h$, DIABLO retains only the features with non-zero entries in $\mathbf{a}_h^{(q)}$. The user specifies how many features to keep per block and component. In the benchmark cancer studies, for example, each biomarker panel consisted of a fixed number of features per omic (e.g. 90 variables with the largest absolute weights on each of two components, resulting in 180 features in total, as done in DIABLO's analyses).

##### 1.6.17 asmbPLS-DA

Zhang and Datta[35] proposed adaptive sparse multi-block partial least square (PLS) discriminant analysis (asmbPLS-DA). It integrates multiple omics blocks, learns block-specific sparse PLS components, and uses them both for classification and for selecting predictive biomarkers across blocks.

**Inputs.** Let $\{\mathbf{X}^{(b)}\}_{b=1}^B$ be $B$ omics blocks with $\mathbf{X}^{(b)} \in \mathbb{R}^{n\times p_b}$, and let $\mathbf{Y} \in \mathbb{R}^{n\times G}$ be the dummy-coded outcome matrix (one column for binary, $G$ columns for multiclass).

**Model.** asmbPLS-DA constructs $J$ orthogonal PLS components that maximize the covariance between a multi-block "super score" and an outcome score. For component $j$ and block $b$,

$$\mathbf{t}_j^{(b)} = \mathbf{X}_j^{(b)} \boldsymbol{\omega}_j^{(b)} / \sqrt{p_b}, \qquad \mathbf{T}_j = \left[\mathbf{t}_j^{(1)}, \ldots, \mathbf{t}_j^{(B)}\right],$$

$$\mathbf{t}_j^{\text{super}} = \mathbf{T}_j \boldsymbol{\omega}_j^{\text{super}}, \qquad \mathbf{u}_j = \mathbf{Y}\mathbf{q}_j,$$

and the component is obtained by

$$\max_{\{\boldsymbol{\omega}_j^{(b)}\},\, \boldsymbol{\omega}_j^{\text{super}},\, \mathbf{q}_j} \quad \operatorname{cov}\left(\mathbf{t}_j^{\text{super}}, \mathbf{u}_j\right) \quad \text{s.t.} \quad \left\|\boldsymbol{\omega}_j^{(b)}\right\|_2 = 1, \; \left\|\boldsymbol{\omega}_j^{\text{super}}\right\|_2 = 1.$$

Sparsity and block adaptivity are enforced by soft-thresholding the block weight vectors,

$$\boldsymbol{\omega}_j^{(b)} \leftarrow \operatorname{sign}\left(\boldsymbol{\omega}_j^{(b)}\right) \circ \left(\left|\boldsymbol{\omega}_j^{(b)}\right| - \lambda_b\right)_+,$$

where $\lambda_b$ is a block-specific threshold chosen as a quantile of $|\boldsymbol{\omega}_j^{(b)}|$, and $(\cdot)_+$ denotes positive part. This retains only the most relevant features in each block.

**Training objectives.** Given the $J$ components, the outcome is predicted as a linear combination of super scores:

$$\hat{\mathbf{Y}} = \sum_{j=1}^{J} \mathbf{t}_j^{\text{super}} \mathbf{q}_j^\top.$$

Multiple decision rules map $\hat{\mathbf{Y}}$ or $\mathbf{t}_j^{\text{super}}$ to class labels, including fixed cutoff (binary), "Max $\mathbf{Y}$" (multiclass), and nearest-centroid rules in super-score space.

**Biomarker identification.** Within each block $b$, feature weights $\boldsymbol{\omega}_j^{(b)}$ from the first few PLS components encode variable importance. The soft-thresholding above yields a set of selected features (non-zero weights) per component and block. The union across components forms the asmbPLS-DA biomarker set in block $b$. In practice, features are ranked by the magnitude of their coefficients, and genes with the largest $I_g^{(b)}$ in each omic are reported as key biomarkers. In the TCGA-BRCA analysis, for example, the authors highlighted top-ranked genes and miRNAs per block by absolute coefficients.

##### 1.6.18 Stabl

Hédou *et al.*[36] proposed Stabl, a supervised feature-selection framework that wraps around sparse regularized models (SRMs, e.g. Lasso/Elastic Net) to identify a set of biomarkers by combining noise injection, subsampling, and a signal-to-noise threshold, while preserving predictive performance.

**Inputs.** Let $\mathbf{X} \in \mathbb{R}^{N\times P}$ be a single-omics feature matrix and $\mathbf{y} \in \{0,1\}^N$ the class labels. For multi-omics, Stabl is typically applied separately to each omic to obtain omic-specific reliable feature sets $\widehat{\mathcal{S}}^{(m)}$ and thresholds $\theta^{(m)}$, which are then concatenated and used in a final SRM.

**Model.** Stabl is a wrapper around a base SRM, typically a sparse logistic regression with Lasso, Elastic Net, Adaptive Lasso, or Sparse Group Lasso penalties. The core steps are:

*Noise injection.* Construct $P$ artificial "null" features $\tilde{\mathbf{X}} \in \mathbb{R}^{N\times P}$ using either Model-X knockoffs[37] or random permutations and form the augmented design

$$\mathbb{X} = [\,\mathbf{X} \mid \tilde{\mathbf{X}}\,] \in \mathbb{R}^{N\times 2P},$$

with index sets of original features $\mathcal{O} = \{1,\ldots,P\}$ and artificial features $\mathcal{A} = \{P+1,\ldots,2P\}$.

*Subsampling and stability paths.* For $k = 1,\ldots,B$ subsampling iterations, draw a half-sample of rows and fit the base SRM on $(\mathbf{y},\mathbb{X})_k$ over a grid of regularization parameters $\lambda \in \Lambda$:

$$\hat{\boldsymbol{\beta}}^{(k)}(\lambda) = \arg\min_{\boldsymbol{\beta}} \Big[\mathcal{L}_{\text{logit}}\big(\mathbf{y}, \mathbb{X}_k\boldsymbol{\beta}\big) + \Omega_\lambda(\boldsymbol{\beta})\Big],$$

where $\mathcal{L}_{\text{logit}}$ is logistic loss and $\Omega_\lambda$ is the chosen sparsity-inducing penalty (e.g. $\ell_1$, $\ell_1+\ell_2$). For feature $j$ and $\lambda$, define the selection frequency

$$f_j(\lambda) = \frac{1}{B}\sum_{k=1}^{B} \mathbb{I}\big[\hat{\beta}_j^{(k)}(\lambda) \neq 0\big], \qquad f_j = \max_{\lambda\in\Lambda} f_j(\lambda),$$

which produces one stability path $f_j(\lambda)$ per feature.

*Data-driven reliability threshold.* For a frequency threshold $t \in [0,1]$, select features with $f_j \geq t$ and define the surrogate false discovery proportion

$$\text{FDP}^+(t) = \frac{1 + \sum_{j\in\mathcal{A}} \mathbb{I}[f_j \geq t]}{\sum_{j\in\mathcal{O}} \mathbb{I}[f_j \geq t] \;\vee\; 1}.$$

Stabl chooses the reliability threshold

$$\theta \in \arg\min_{t\in[0,1]} \text{FDP}^+(t),$$

and the reliable feature set

$$\widehat{\mathcal{S}} = \big\{j \in \mathcal{O} : f_j \geq \theta\big\}.$$

*Final predictive model.* The base SRM is refit on the original features restricted to $\widehat{\mathcal{S}}$:

$$\hat{\boldsymbol{\beta}}^{\text{Stabl}} = \arg\min_{\boldsymbol{\beta}} \Big[\mathcal{L}_{\text{logit}}\big(\mathbf{y}, \mathbf{X}_{\widehat{\mathcal{S}}}\boldsymbol{\beta}\big) + \Omega_{\lambda^\star}(\boldsymbol{\beta})\Big],$$

yielding class probabilities

$$\hat{\mathbf{p}}^{(s)} = \sigma\big(\mathbf{x}_{\widehat{\mathcal{S}}}^{(s)\top}\hat{\boldsymbol{\beta}}^{\text{Stabl}}\big),$$

with $\sigma$ the logistic sigmoid.

**Training objectives.** The base learner is a sparse logistic model optimized with cross-entropy plus regularization:

$$\mathcal{L}_{\text{CE}} = -\frac{1}{|\mathcal{B}|}\sum_{s\in\mathcal{B}} \Big[y_s \log \hat{p}^{(s)} + (1-y_s)\log\big(1-\hat{p}^{(s)}\big)\Big] + \Omega_\lambda(\boldsymbol{\beta}),$$

where $\Omega_\lambda$ is e.g. Lasso, Elastic Net, Adaptive Lasso, or Sparse Group Lasso penalty, and $\mathcal{B}$ is a training batch.

**Biomarker identification.** Candidate biomarkers are the features deemed reliable by Stabl. For feature $j$, its selection frequency $f_j$ summarizes robustness across subsamples and regularization strengths. Features with $f_j \geq \theta$ form $\widehat{\mathcal{S}}$ and enter the final Stabl model. A direct importance score is $I_j = f_j$. In practice, Stabl reports a short list of features with $f_j \geq \theta$.

###### 1.6.19 GDF

Pfeifer *et al.*[38] proposed Greedy Decision Forest (GDF), a network-based classifier that builds decision trees whose features are restricted to connected nodes in a biological graph. It performs a greedy search over such trees to select high-performing disease modules, and assigns importance scores to subnetworks, edges, and node-level multi-omics features.

**Inputs.** Let $\mathbf{A} \in \mathbb{R}^{|\mathcal{V}|\times|\mathcal{V}|}$ be the adjacency matrix of a PPI network $G = (\mathcal{V}, \mathcal{E})$, and let $\mathbf{X} \in \mathbb{R}^{N\times P}$ be the multi-omics feature matrix for $N$ samples and $P$ molecular features. Each node $v \in \mathcal{V}$ corresponds to multi-omics features. We write $f(v)$ for the set of feature indices associated with node $v$. Class labels are $\mathbf{y} \in \{0, 1, \ldots, C-1\}^N$.

**Model.** GDF constructs a decision forest whose trees are restricted to connected node sets on $G$ and then runs a greedy module-selection procedure.

*Network-constrained trees (random-walk sampling).* To build the $k$-th tree, a random walk of fixed depth $L$ (initially $L \approx \sqrt{|\mathcal{V}|}$) is started at a random node on $G$. The set of visited nodes defines a module $X_k^G \subseteq \mathcal{V}$. The associated feature subset is

$$\mathcal{F}_k \;=\; \bigcup_{v\in X_k^G} f(v) \;\subseteq\; \{1, \ldots, P\}.$$

A decision tree $T_k$ is trained on $\mathbf{X}_{\mathcal{F}_k}$ to predict $\mathbf{y}$, using Gini impurity for splits. Repeating this procedure leads to an initial forest $\{T_k\}_{k=1}^{n_{\text{tree}}}$ and corresponding node sets $\{X_k^G\}$.

*Greedy module refinement.* The forest is refined over $n_{\text{iter}}$ greedy iterations. At iteration $t$, each tree $T_k[t]$ is evaluated by its out-of-bag performance

$$\text{Perf}\big(T_k[t]\big) \in [0, 1] \quad \text{(e.g. AUROC)}.$$

If $\text{Perf}(T_k[t]) < \text{Perf}(T_k[t-1])$, the tree and its node set are kept from the previous iteration; otherwise the depth is reduced by one and a new random walk on the induced subgraph is used to update $X_k^G[t]$. After updating all trees, a new set of $n_{\text{tree}}$ trees is resampled in proportion to $\text{Perf}(T_k[t])$ and the procedure repeats. After $n_{\text{iter}}$ iterations, each surviving tree $T_m$ defines a candidate disease module $X_m^G$.

**Training objectives.** Each decision tree in the forest is trained with standard Gini-based splitting. For a node $v_t$ with class proportions $\hat{p}_{v_t,c}$,

$$\text{Gini}(v_t) = \sum_{c=1}^{C} \hat{p}_{v_t,c}\,\big(1 - \hat{p}_{v_t,c}\big),$$

and the information gain of feature $j$ at $v_t$ is

$$\text{Gain}(X_j, v_t) = \text{Gini}(v_t) - w_L\,\text{Gini}(v_L) - w_R\,\text{Gini}(v_R),$$

where $v_L, v_R$ are child nodes and $w_L, w_R$ are their sample proportions. The splitting rule chooses the feature and threshold with maximal gain at each node.

**Biomarker identification.** Here we focus on the descriptions of node-level importance derivation. For a node $v$ with feature set $f(v)$, GDF aggregates Gini gains over all splits and all trees where any feature from $f(v)$ is used:

$$\text{IMP}_f\big(f(v) \in X_m^G\big) = \frac{1}{n_{\text{iter}}\, n_{\text{tree}}} \sum_{t,k:\, v\in X_m^G} \; \sum_{j\in f(v)} \; \sum_{v_t \; \text{splits on } j} \text{Gain}(X_j, v_t).$$

producing a node-level score that can be interpreted as gene-level importance in the detected modules.

###### 1.6.20 DPM

Slobodyanyuk *et al.*[39] proposed Directional $p$-value Merging (DPM), a data fusion method that integrates multi-omics at the gene level by combining $p$-values and direction signs under user-defined directional constraints.

**Inputs.** Let $\{\mathbf{P}^{(i)}\}_{i=1}^{k}$ be $k$ omics datasets, each providing gene-level $p$-values, with $\mathbf{P}^{(i)} \in \mathbb{R}^G$ for $G$ genes. Such $p$-values could be derived via Mann-Whitney U-tests across different sample groups, as demonstrated in the original analyses. For dataset $i$, denote the $p$-value of gene $g$ by $P_{g,i}$. Directional information is encoded as unit signs

$$o_{g,i} \in \{-1, 0, +1\},$$

typically derived from log-fold changes, correlation coefficients, log-hazard ratios, or regression coefficients (sign only; 0 for directionless inputs). A user-defined constraints vector

$$\mathbf{e} = (e_1, \ldots, e_k)^\top, \qquad e_i \in \{-1, 0, +1\},$$

specifies how each dataset $i$ is expected to relate directionally to the others (e.g. $e_i = +1$ for direct, $e_i = -1$ for inverse, $e_i = 0$ for no directional constraint).

**Model.** Datasets are partitioned into a directional set $\{1, \ldots, j\}$ and a directionless set $\{j+1, \ldots, k\}$. For gene $g$, DPM forms a directionally weighted Fisher-like score

$$S_g^{\text{dir}} = \sum_{i=1}^{j} \ln P_{g,i}\, o_{g,i} e_i, \qquad S_g^{\text{nodir}} = \sum_{i=j+1}^{k} \ln P_{g,i},$$

and defines

$$X_g^{\text{DPM}} = -2\Big(\big|S_g^{\text{dir}}\big| + S_g^{\text{nodir}}\Big).$$

Directional agreements ($o_{g,i}e_i$ having the same sign across datasets) increase $\big|S_g^{\text{dir}}\big|$ and thus make $X_g^{\text{DPM}}$ more significant, whereas conflicts shrink $\big|S_g^{\text{dir}}\big|$ and penalise the gene. The absolute value ensures global sign invariance of the constraints vector (e.g. $[+1, +1] \equiv [-1, -1]$).

To obtain a merged $p$-value that accounts for $p$-value dependencies across datasets, DPM uses the empirical Brown's method: $X_g^{\text{DPM}}$ is approximated by a scaled $\chi^2$ distribution $c\,\chi^2_{k'}$, where the scaling factor $c$ and effective degrees of freedom $k'$ are estimated from the covariance of log $p$-values across datasets. The merged $p$-value is then

$$P_g^{\text{DPM}} = 1 - F_{\chi^2_{k'}}\left(\frac{X_g^{\text{DPM}}}{c}\right),$$

where $F_{\chi^2_{k'}}$ is the CDF of the $\chi^2$ distribution with $k'$ degrees of freedom. Small $P_g^{\text{DPM}}$ indicate genes with significant and directionally consistent multi-omics evidence under the chosen $\mathbf{e}$.

**Biomarker identification.** DPM produces a ranked gene list based on the merged $p$-values $\{P_g^{\text{DPM}}\}$. Genes with small $P_g^{\text{DPM}}$ are directionally prioritised. Comparing DPM to a non-directional Brown merge, genes that are significant in the Brown analysis but not under DPM are termed directionally penalised, as their signals conflict with the directional constraints. For a chosen significance threshold (e.g. $P_g^{\text{DPM}} < 0.05$ or FDR on $\{P_g^{\text{DPM}}\}$), the prioritised genes are taken as candidate biomarkers with consistent multi-omics regulation (e.g. concordant transcript-protein survival effects or inverse methylation-expression changes), whereas penalised genes highlight discordant signals that may reflect complex or context-specific regulation.

#### 1.7 Predictive performance metrics

We assessed binary predictive performance using the area under the receiver operating characteristic curve (AUROC) and the area under the precision-recall curve (AUPR). For a given decision threshold $t$, predicted labels are summarized in terms of true positives (TP), false positives (FP), true negatives (TN) and false negatives (FN). Define

$$\text{TPR}(t) = \frac{\text{TP}(t)}{\text{TP}(t) + \text{FN}(t)}, \qquad \text{FPR}(t) = \frac{\text{FP}(t)}{\text{FP}(t) + \text{TN}(t)},$$

$$\text{Precision}(t) = \frac{\text{TP}(t)}{\text{TP}(t) + \text{FP}(t)}, \qquad \text{Recall}(t) = \text{TPR}(t).$$

The receiver operating characteristic (ROC) curve plots $\text{TPR}(t)$ against $\text{FPR}(t)$ as $t$ varies over all possible thresholds, and AUROC is the area under this curve. The precision-recall (PR) curve plots $\text{Precision}(t)$ against $\text{Recall}(t)$, and the corresponding area under this curve defines AUPR, which is particularly informative in the presence of substantial class imbalance.

### 2 Supplementary Figures

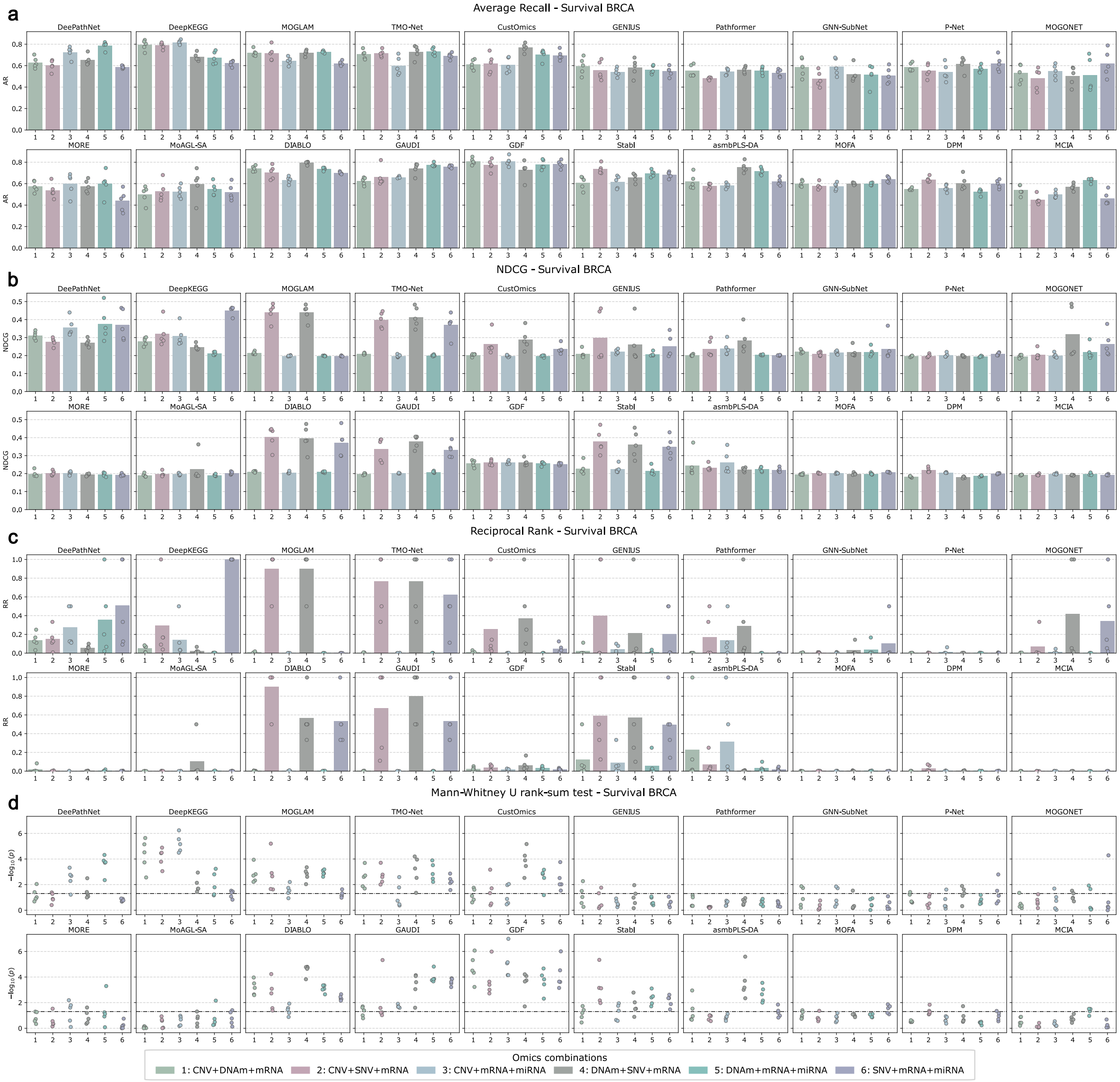

**Supplementary Figure 1: Biomarker identification accuracy by omics combinations for the BRCA survival task.** The performance of five-fold cross-validation is shown, measured by AR (**a**), NDCG (**b**), RR (**c**), and Mann-Whitney U test (**d**). The horizontal dotted line in (d) denotes $p = 0.05$. AR: Average Recall; NDCG: Normalized Discounted Cumulative Gain; RR: Reciprocal Rank.

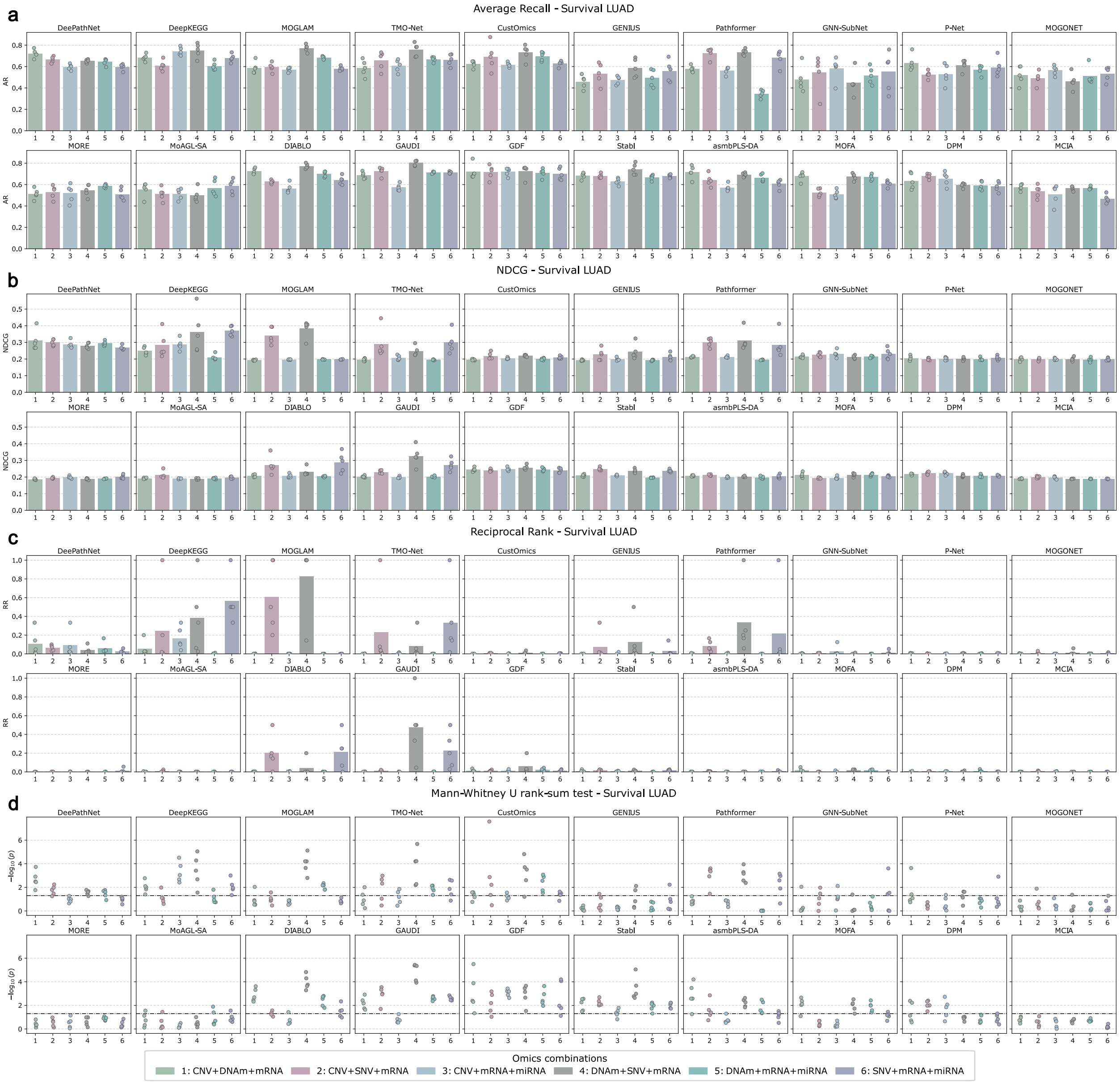

**Supplementary Figure 2: Biomarker identification accuracy by omics combinations for the LUAD survival task.** The performance of five-fold cross-validation is shown, measured by AR (**a**), NDCG (**b**), RR (**c**), and Mann-Whitney U test (**d**). The horizontal dotted line in (d) denotes $p = 0.05$. AR: Average Recall; NDCG: Normalized Discounted Cumulative Gain; RR: Reciprocal Rank.

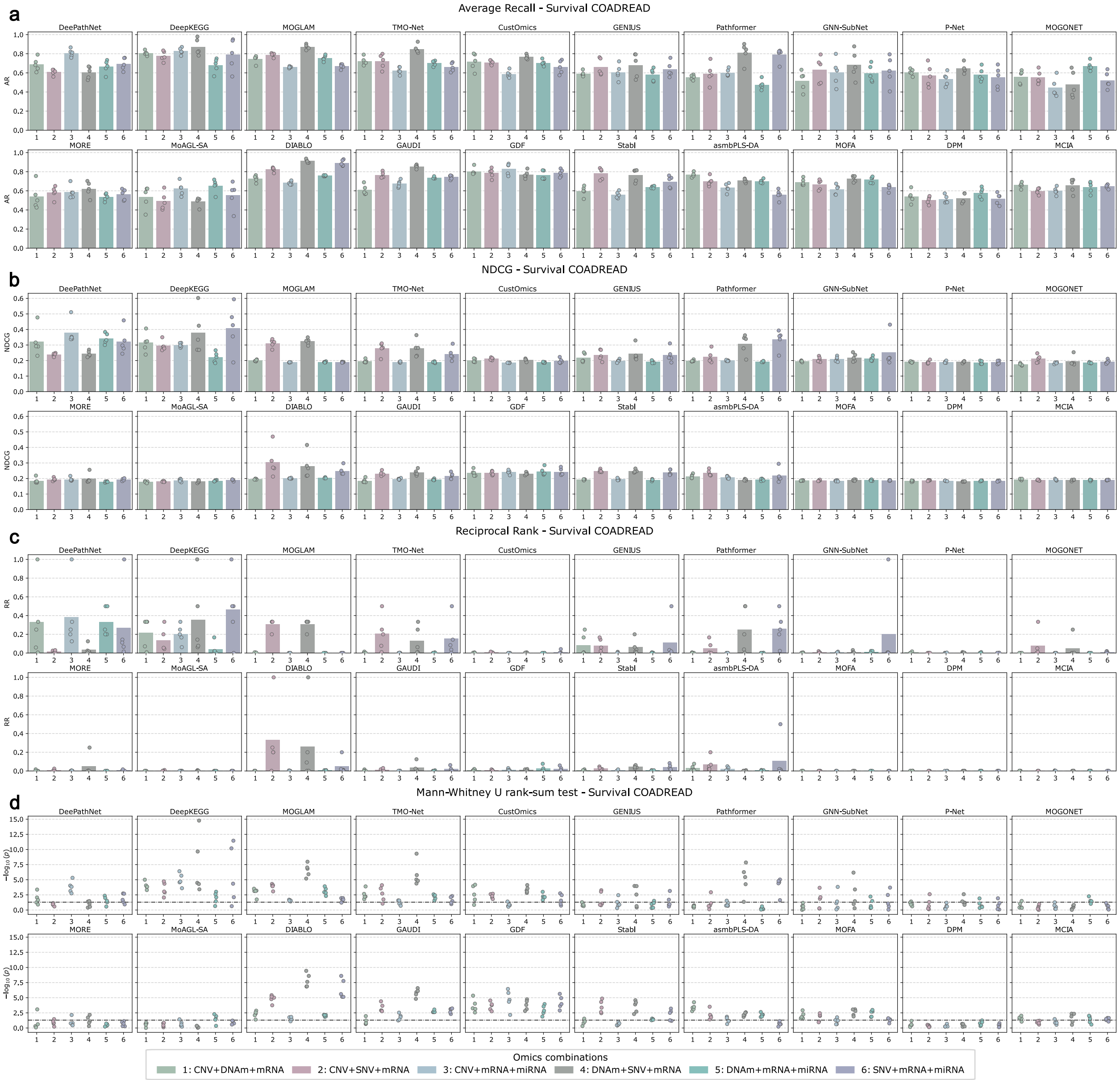

**Supplementary Figure 3: Biomarker identification accuracy by omics combinations for the COADREAD survival task.** The performance of five-fold cross-validation is shown, measured by AR (**a**), NDCG (**b**), RR (**c**), and Mann-Whitney U test (**d**). The horizontal dotted line in (d) denotes $p = 0.05$. AR: Average Recall; NDCG: Normalized Discounted Cumulative Gain; RR: Reciprocal Rank.

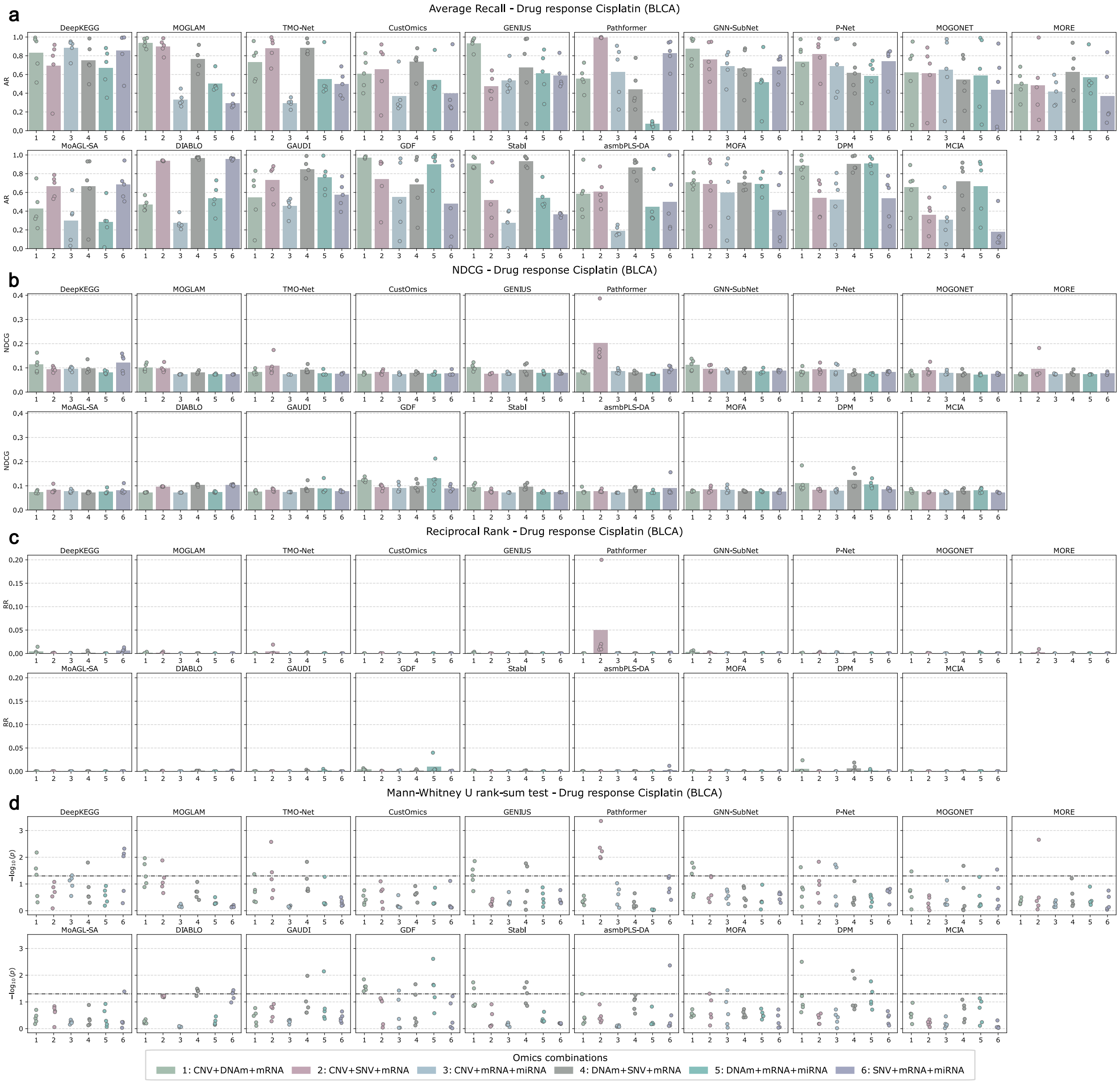

**Supplementary Figure 4: Biomarker identification accuracy by omics combinations for the Cisplatin drug response (BLCA) task.** The performance of five-fold cross-validation is shown, measured by AR (**a**), NDCG (**b**), RR (**c**), and Mann-Whitney U test (**d**). The horizontal dotted line in (d) denotes $p = 0.05$. AR: Average Recall; NDCG: Normalized Discounted Cumulative Gain; RR: Reciprocal Rank.

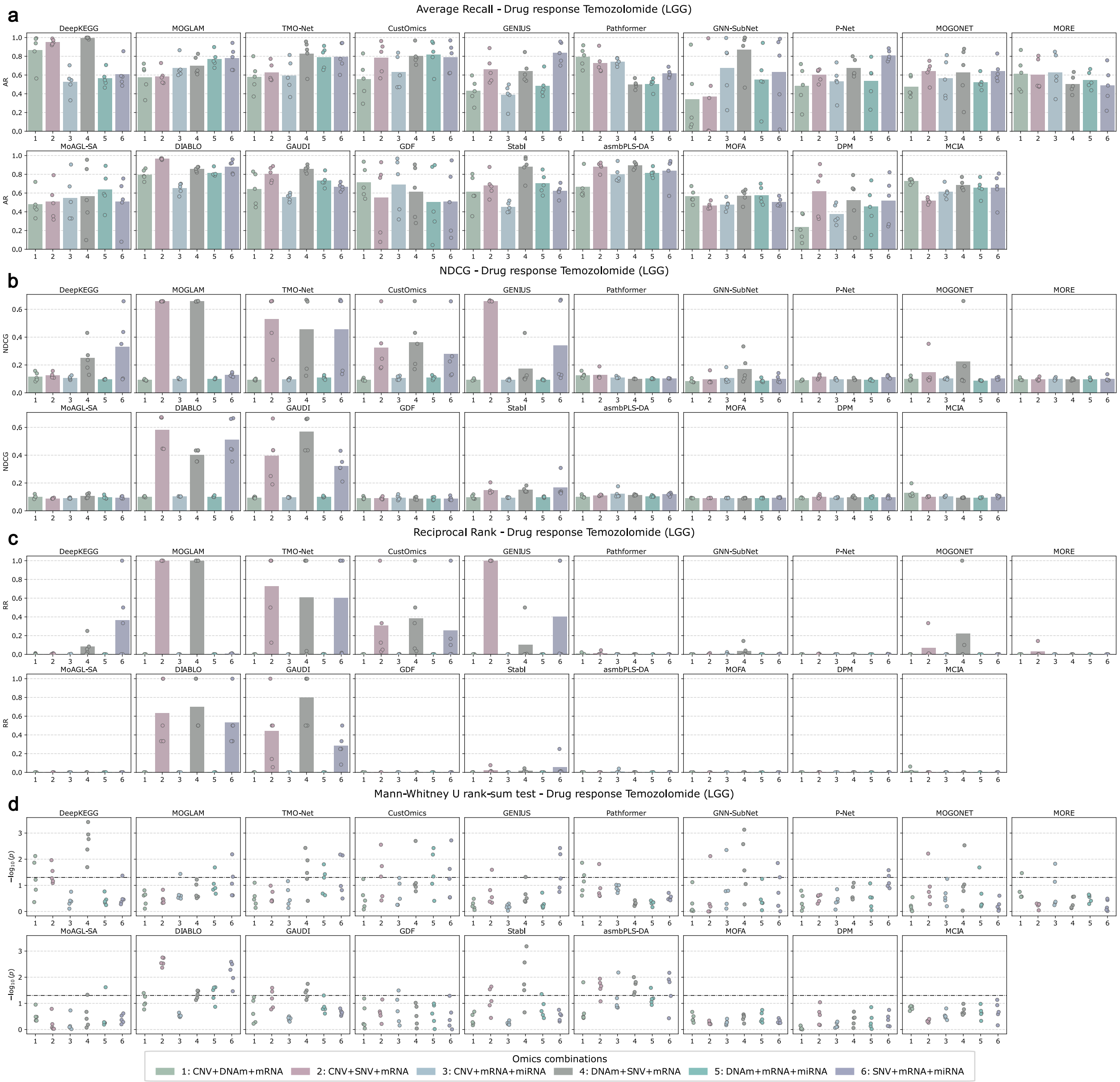

**Supplementary Figure 5: Biomarker identification accuracy by omics combinations for the Temozolomide drug response (LGG) task.** The performance of five-fold cross-validation is shown, measured by AR (**a**), NDCG (**b**), RR (**c**), and Mann-Whitney U test (**d**). The horizontal dotted line in (d) denotes $p = 0.05$. AR: Average Recall; NDCG: Normalized Discounted Cumulative Gain; RR: Reciprocal Rank.

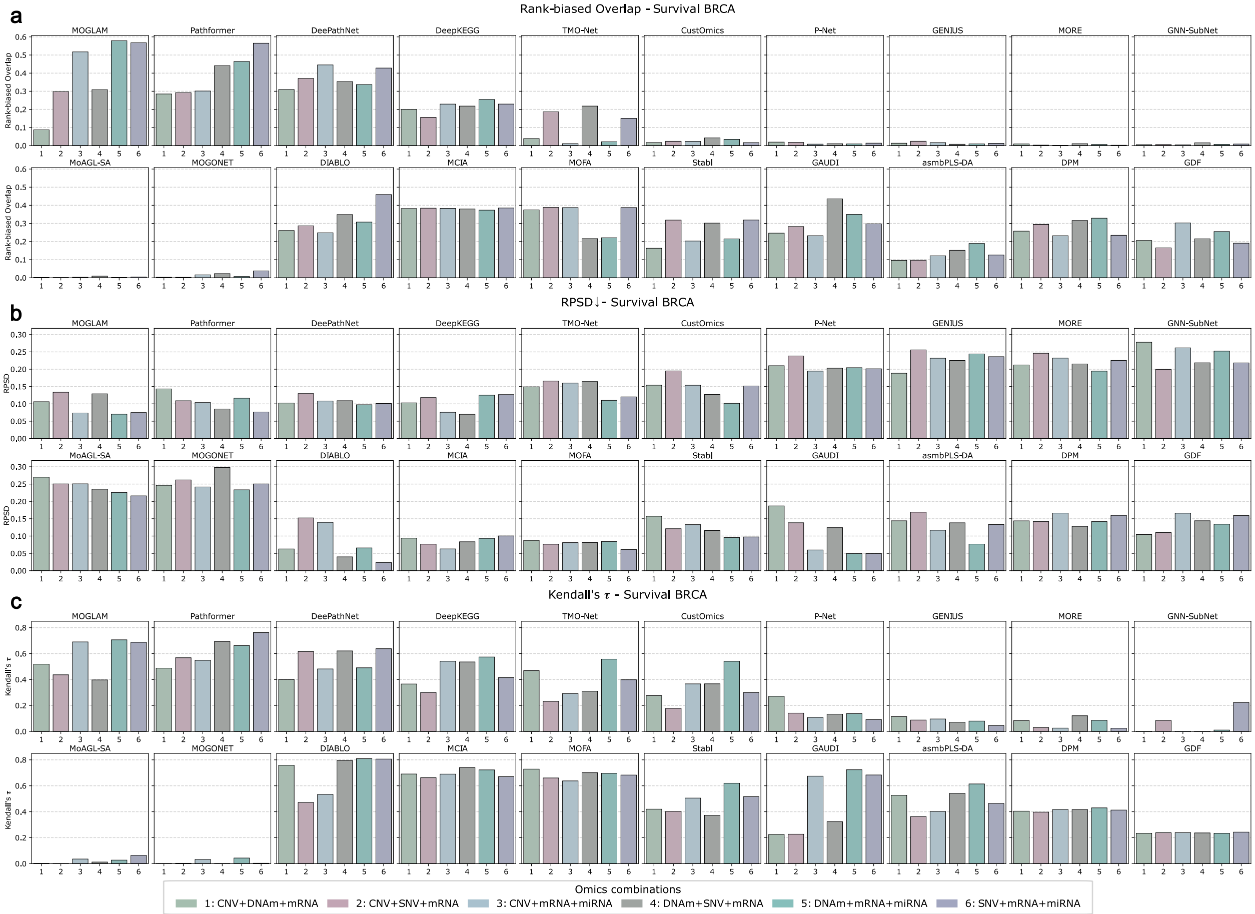

**Supplementary Figure 6: Biomarker identification stability by omics combinations for the BRCA survival task.** The stability of biomarker identification among five-fold cross-validation is shown, measured by Rank-biased Overlap (**a**), RPSD (**b**), and Kendall's $\tau$ (**c**). RPSD: Rank Percentile Standard Deviation.

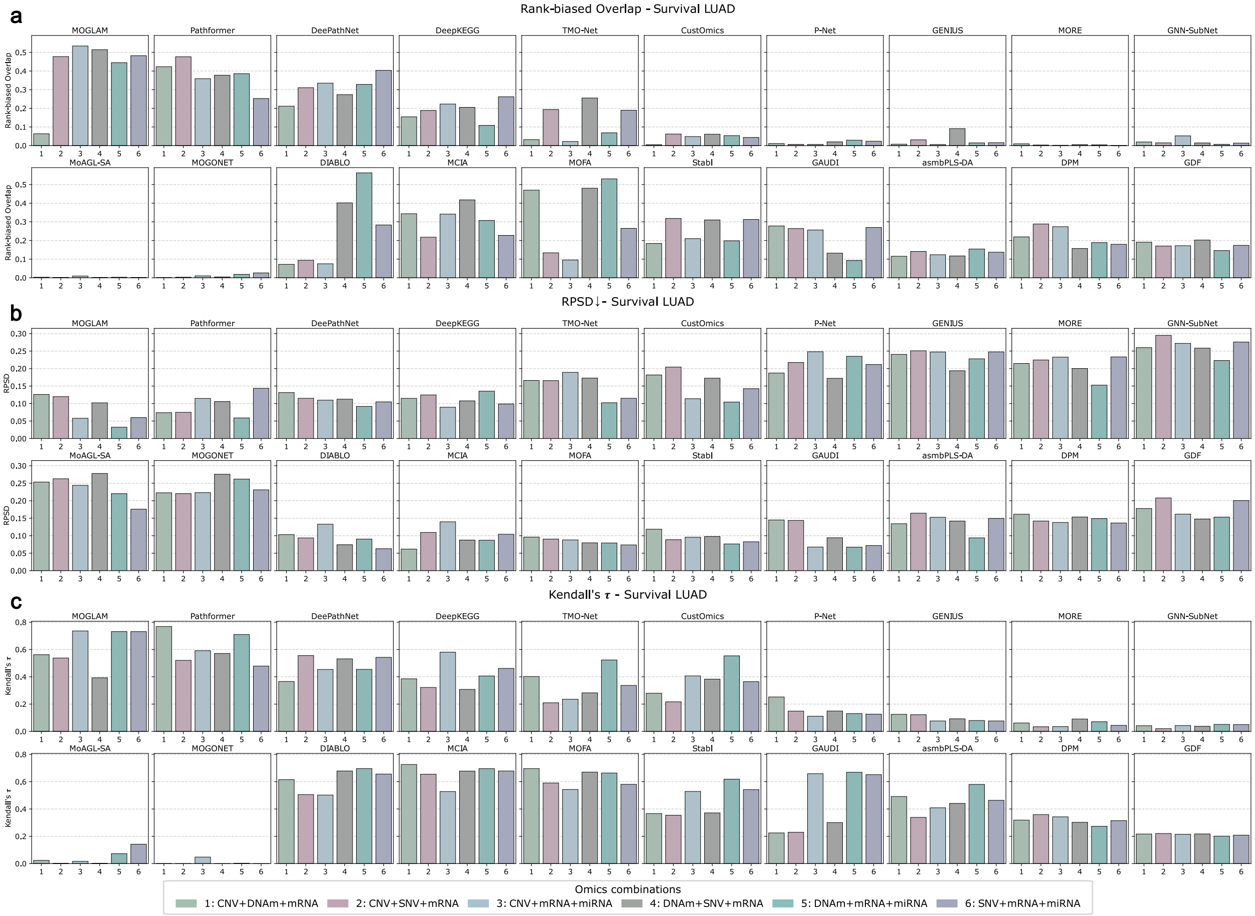

**Supplementary Figure 7: Biomarker identification stability by omics combinations for the LUAD survival task.** The stability of biomarker identification among five-fold cross-validation is shown, measured by Rank-biased Overlap (**a**), RPSD (**b**), and Kendall's $\tau$ (**c**). RPSD: Rank Percentile Standard Deviation.

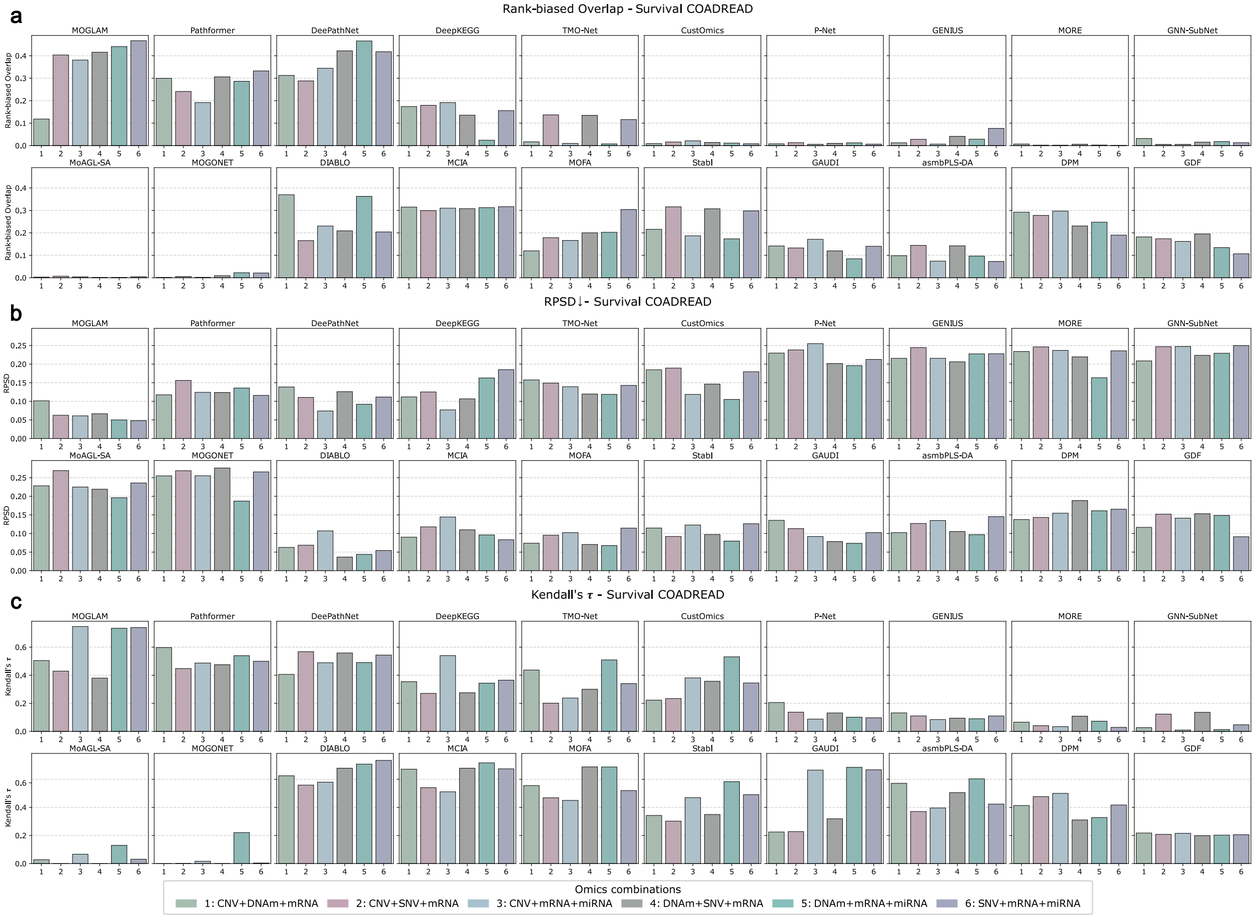

**Supplementary Figure 8: Biomarker identification stability by omics combinations for the COADREAD survival task.** The stability of biomarker identification among five-fold cross-validation is shown, measured by Rank-biased Overlap (**a**), RPSD (**b**), and Kendall's $\tau$ (**c**). RPSD: Rank Percentile Standard Deviation.

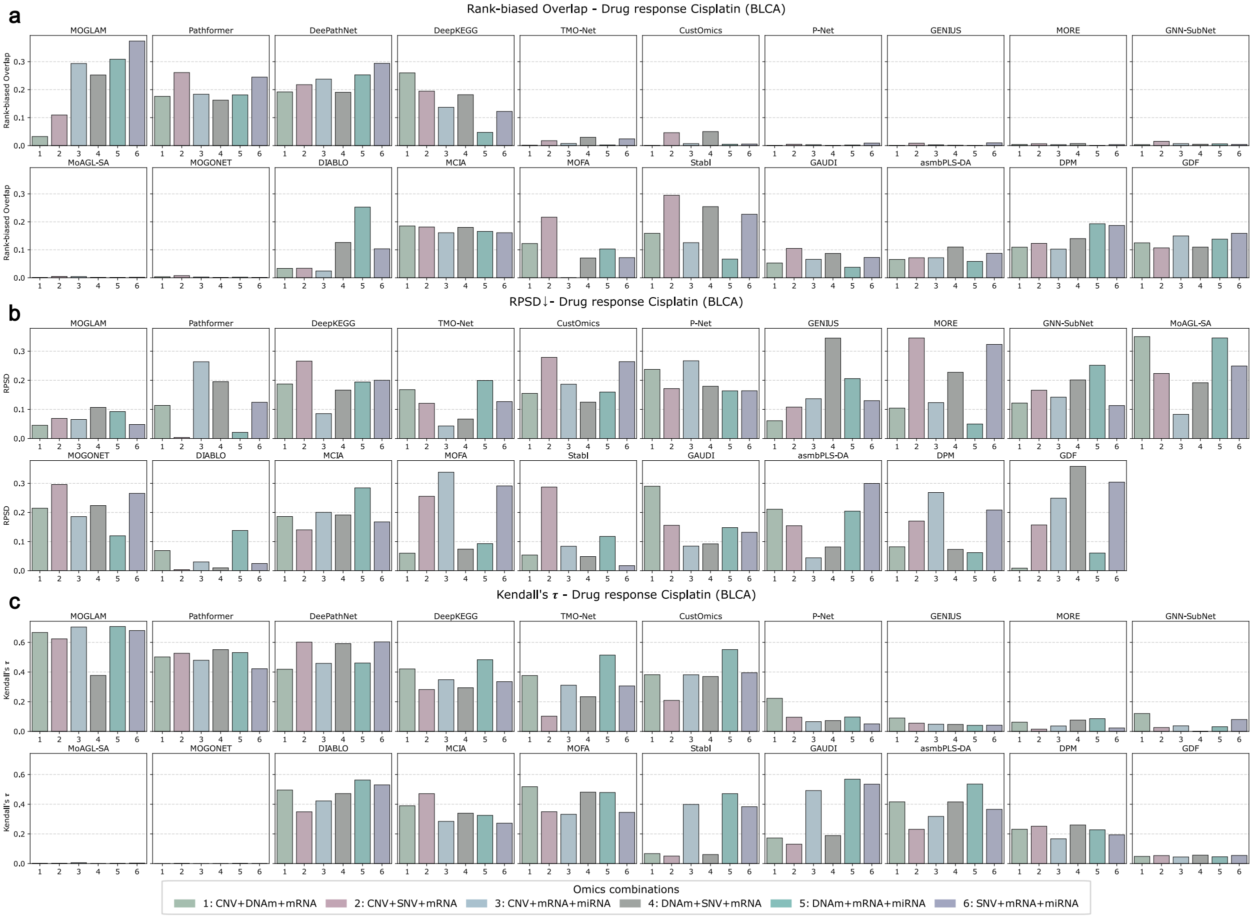

**Supplementary Figure 9: Biomarker identification stability by omics combinations for the Cisplatin drug response (BLCA) task.** The stability of biomarker identification among five-fold cross-validation is shown, measured by Rank-biased Overlap (**a**), RPSD (**b**), and Kendall's $\tau$ (**c**). RPSD: Rank Percentile Standard Deviation.

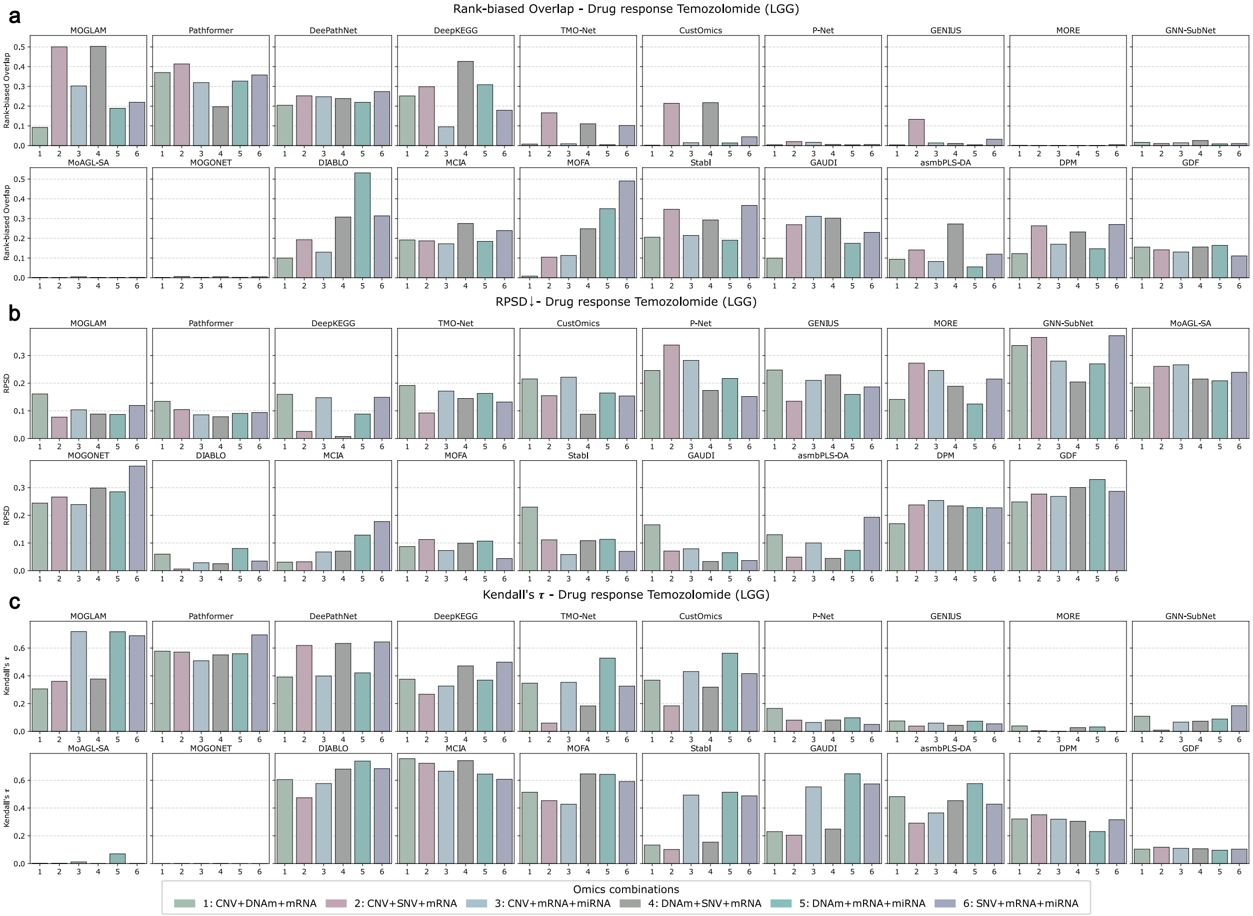

**Supplementary Figure 10: Biomarker identification stability by omics combinations for the Temozolomide drug response (LGG) task.** The stability of biomarker identification among five-fold cross-validation is shown, measured by Rank-biased Overlap (**a**), RPSD (**b**), and Kendall's $\tau$ (**c**). RPSD: Rank Percentile Standard Deviation.

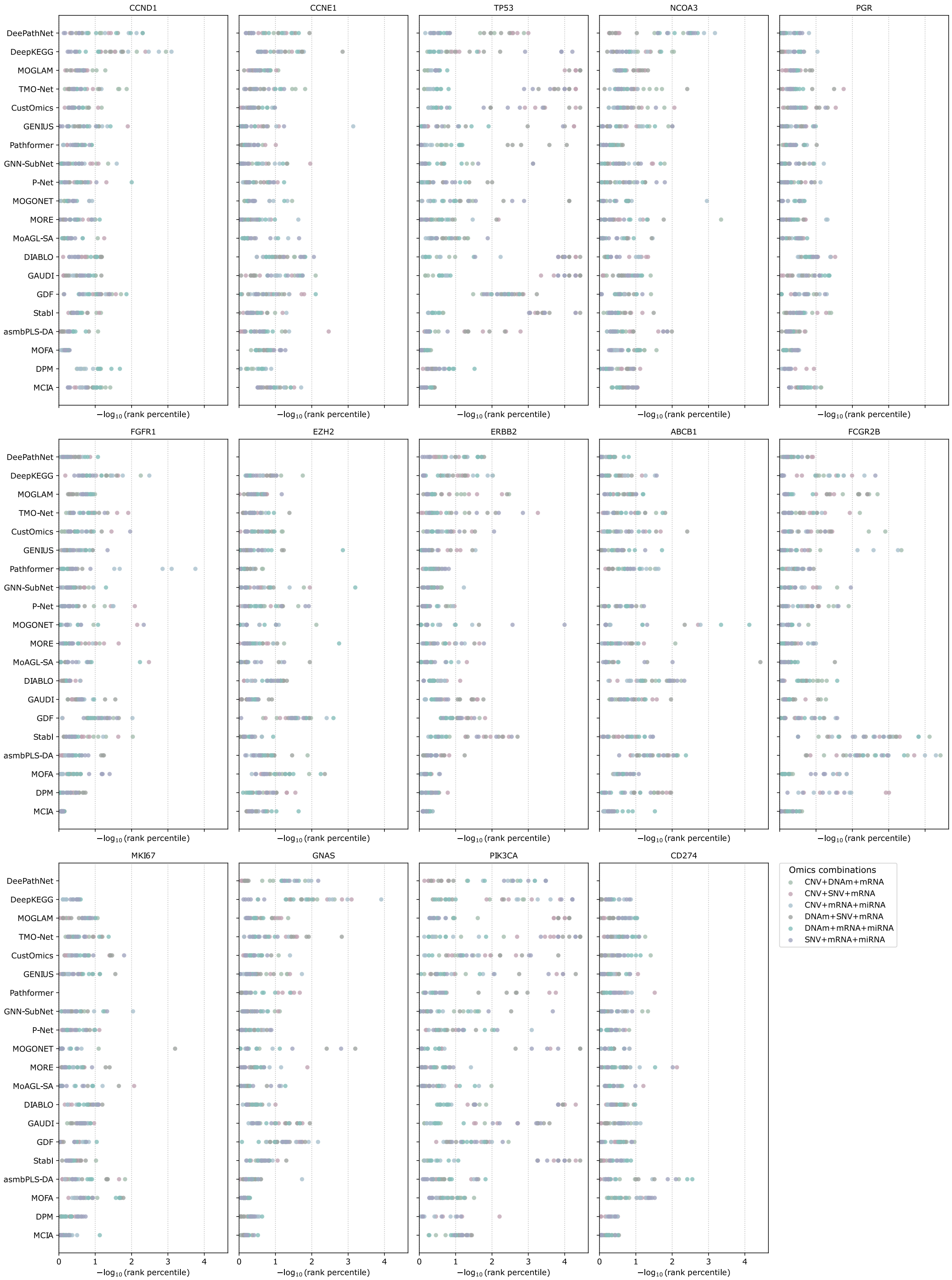

**Supplementary Figure 11: Ranks of each biomarker by omics combinations for the BRCA survival task.** Rankings are shown as negative log-transformed (base 10) rank percentiles.

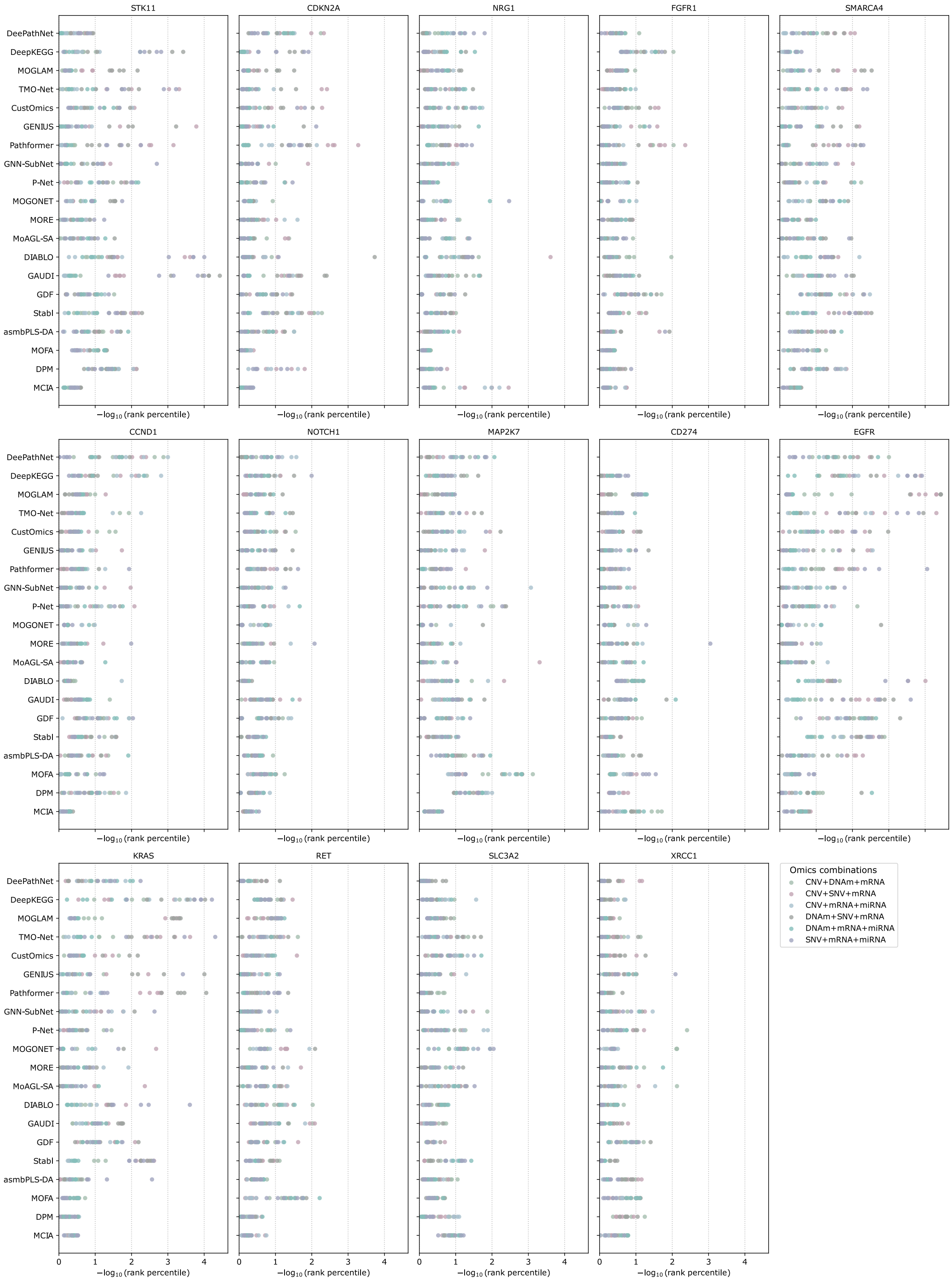

**Supplementary Figure 12: Ranks of each biomarker by omics combinations for the LUAD survival task.** Rankings are shown as negative log-transformed (base 10) rank percentiles.

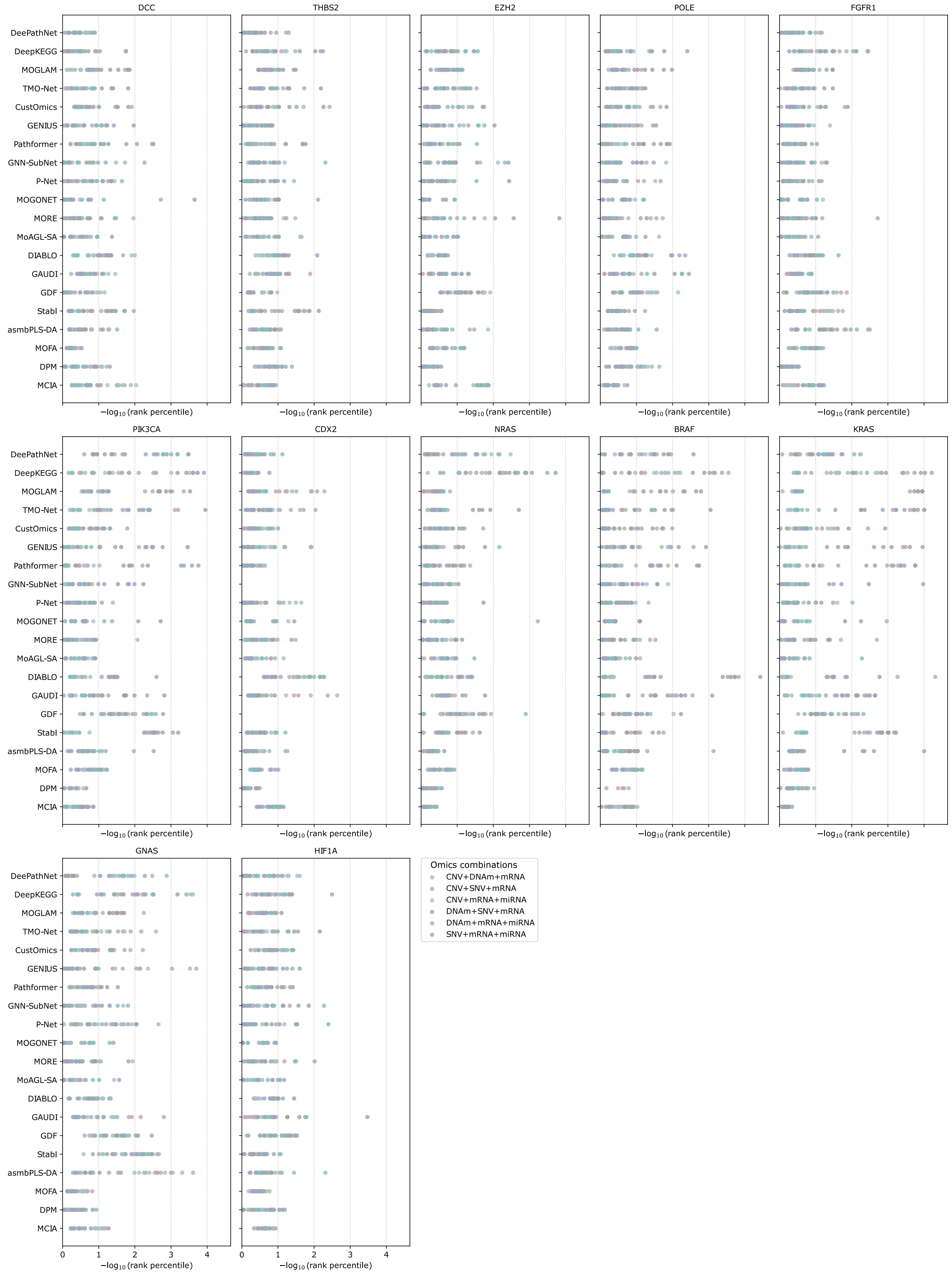

**Supplementary Figure 13: Ranks of each biomarker by omics combinations for the COAD-READ survival task.** Rankings are shown as negative log-transformed (base 10) rank percentiles.

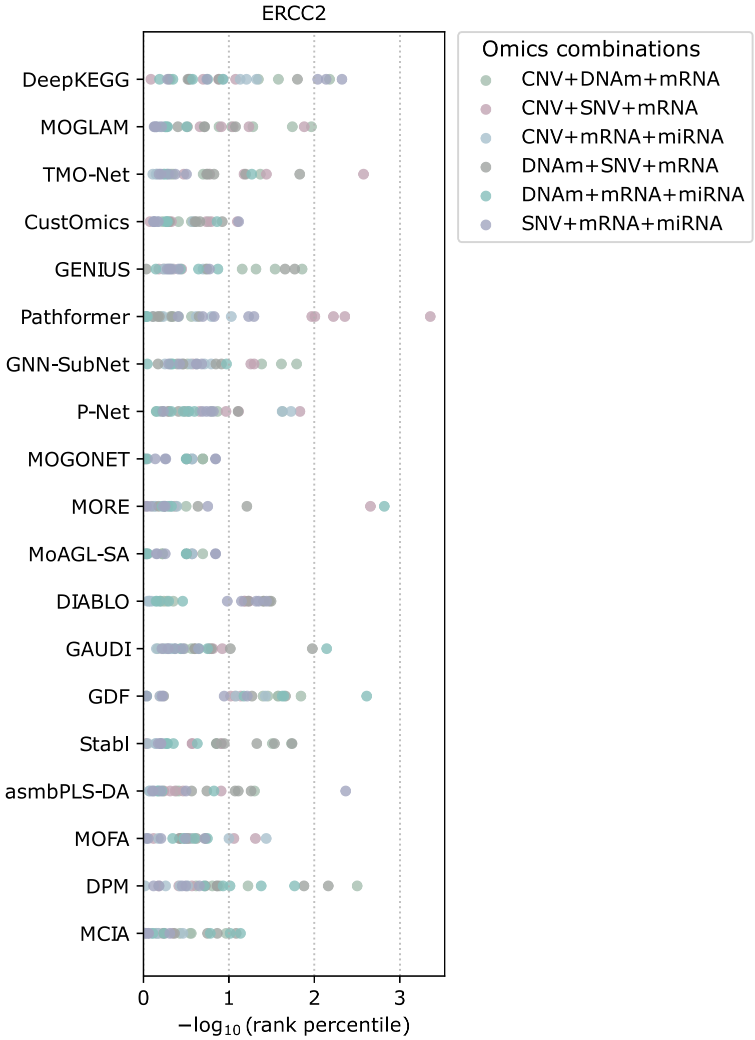

**Supplementary Figure 14: Ranks of each biomarker by omics combinations for the Cisplatin drug response (BLCA) task.** Rankings are shown as negative log-transformed (base 10) rank percentiles.

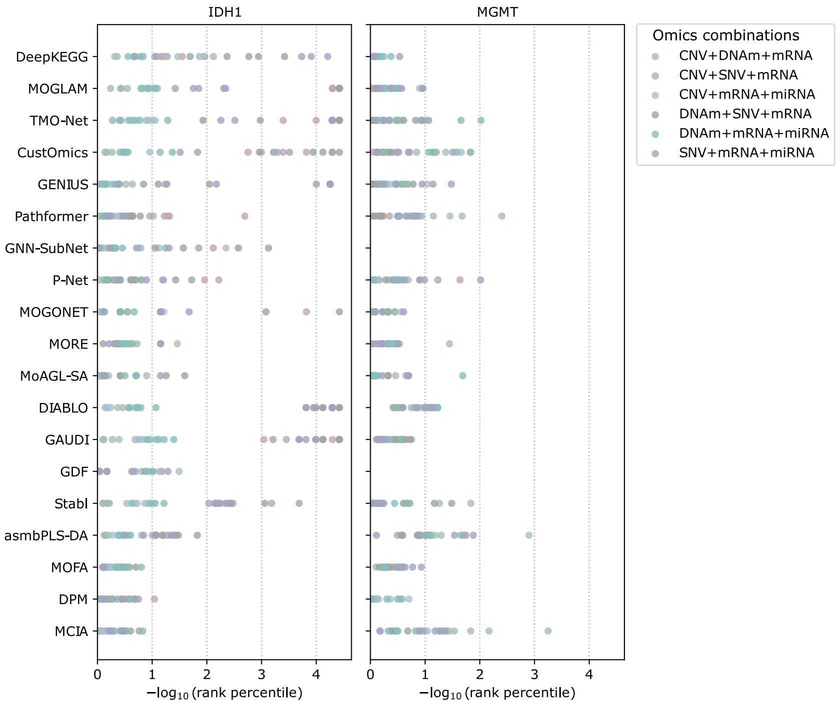

**Supplementary Figure 15: Ranks of each biomarker by omics combinations for the Temozolomide drug response (LGG) task.** Rankings are shown as negative log-transformed (base 10) rank percentiles.

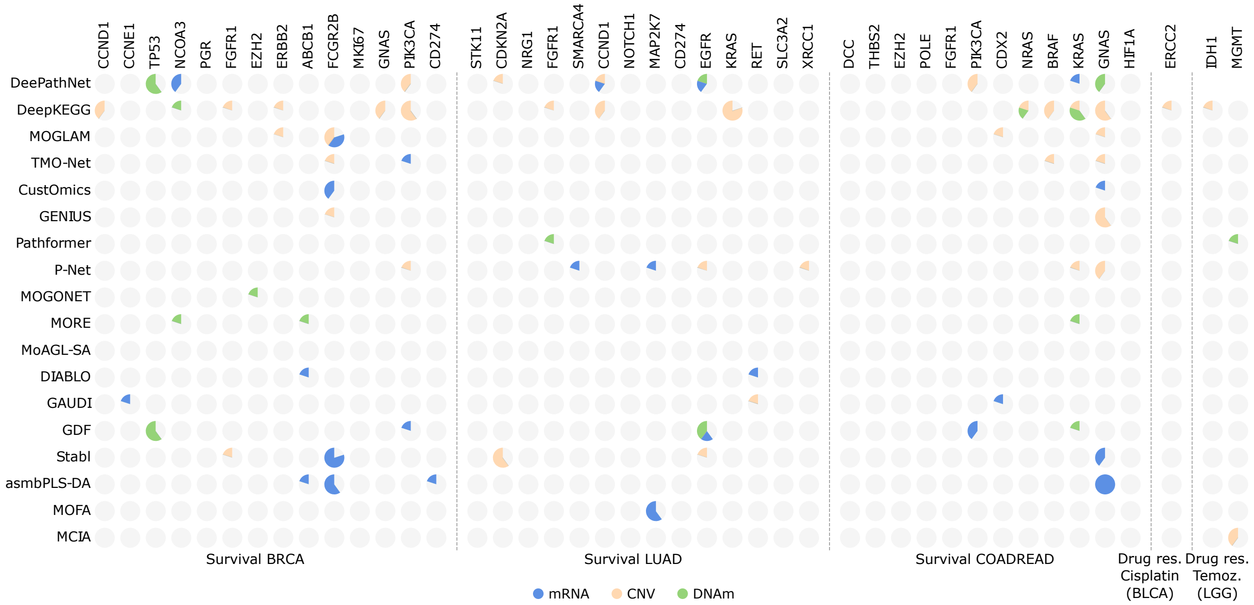

**Supplementary Figure 16: Summary of dominant omics types for biomarkers ranked in the top 1% from experiments with the mRNA+CNV+DNAm omics combination.** Each pie has 5 sections corresponding to the five-fold cross-validation. A section has gray color if the biomarker is not ranked within top 1%.

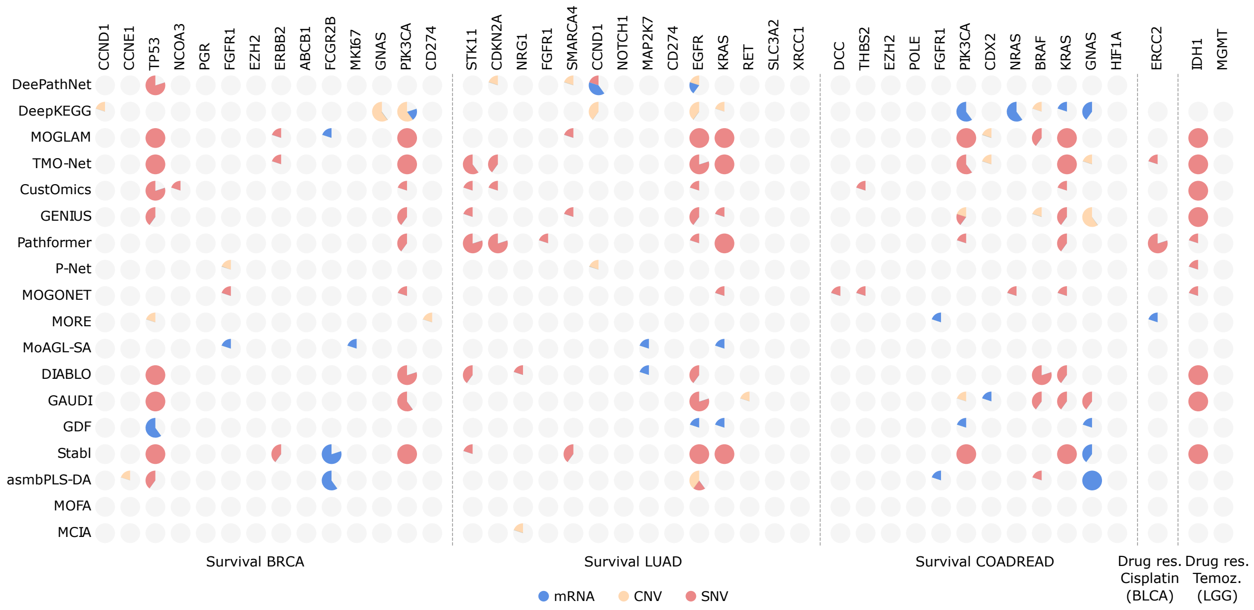

**Supplementary Figure 17: Summary of dominant omics types for biomarkers ranked in the top 1% from experiments with mRNA+CNV+SNV.** Each pie has 5 sections corresponding to the five-fold cross-validation. A section has gray color if the biomarker is not ranked within top 1%.

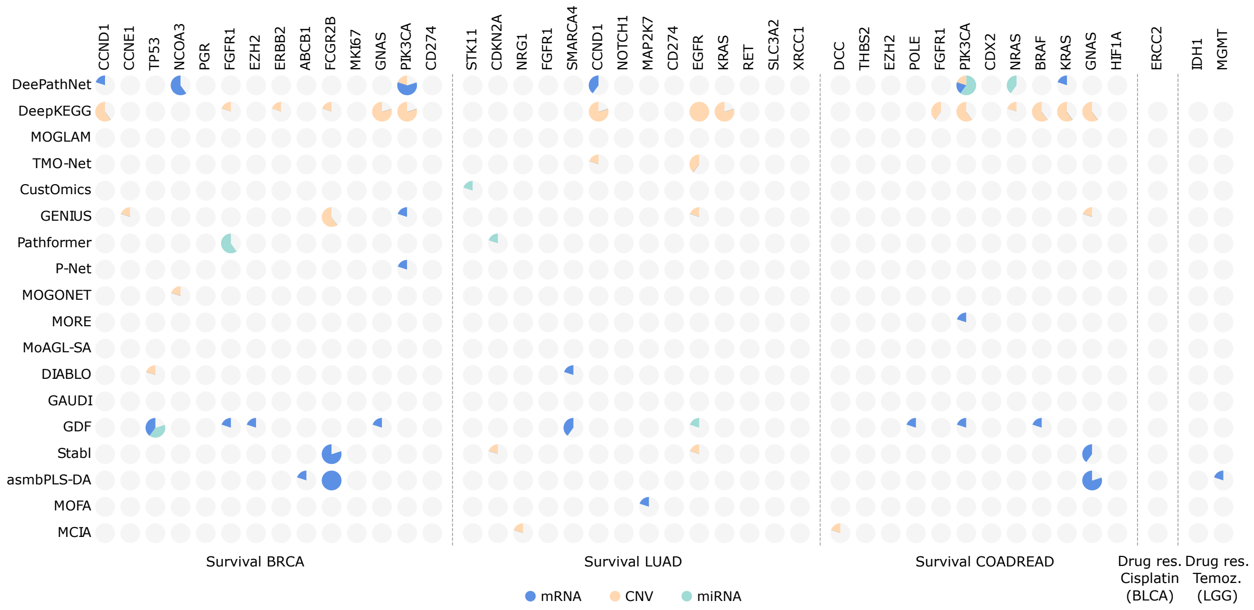

**Supplementary Figure 18: Summary of dominant omics types for biomarkers ranked in the top 1% from experiments with the mRNA+CNV+miRNA omics combination.** Each pie has 5 sections corresponding to the five-fold cross-validation. A section has gray color if the biomarker is not ranked within top 1%.

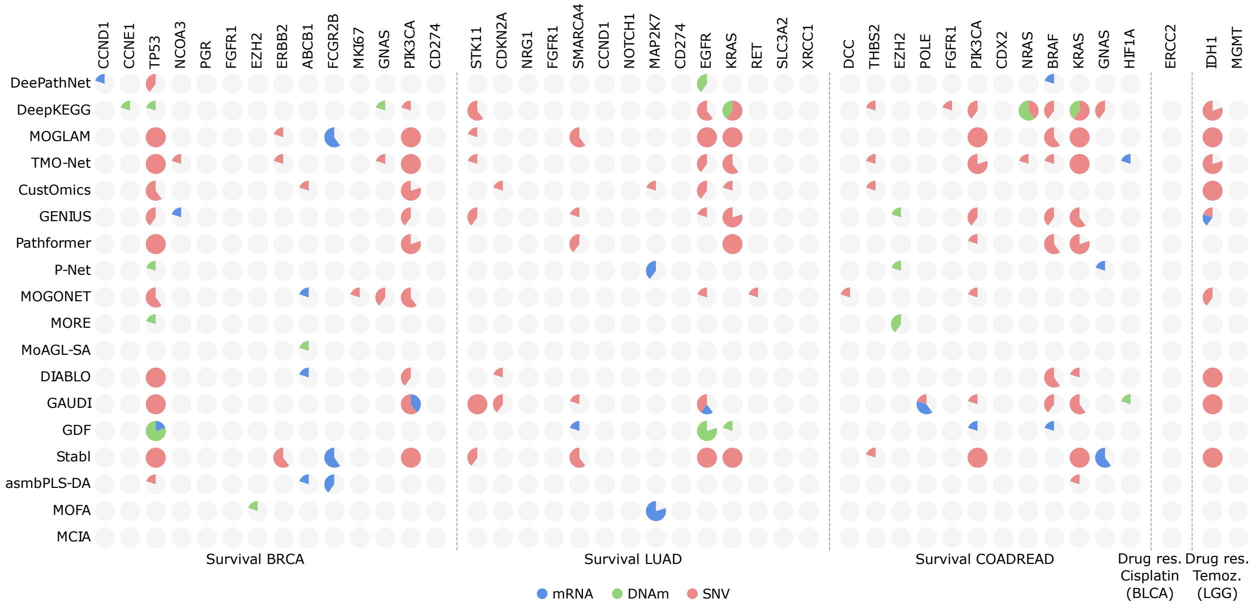

**Supplementary Figure 19: Summary of dominant omics types for biomarkers ranked in the top 1% from experiments with the mRNA+DNAm+SNV omics combination.** Each pie has 5 sections corresponding to the five-fold cross-validation. A section has gray color if the biomarker is not ranked within top 1%.

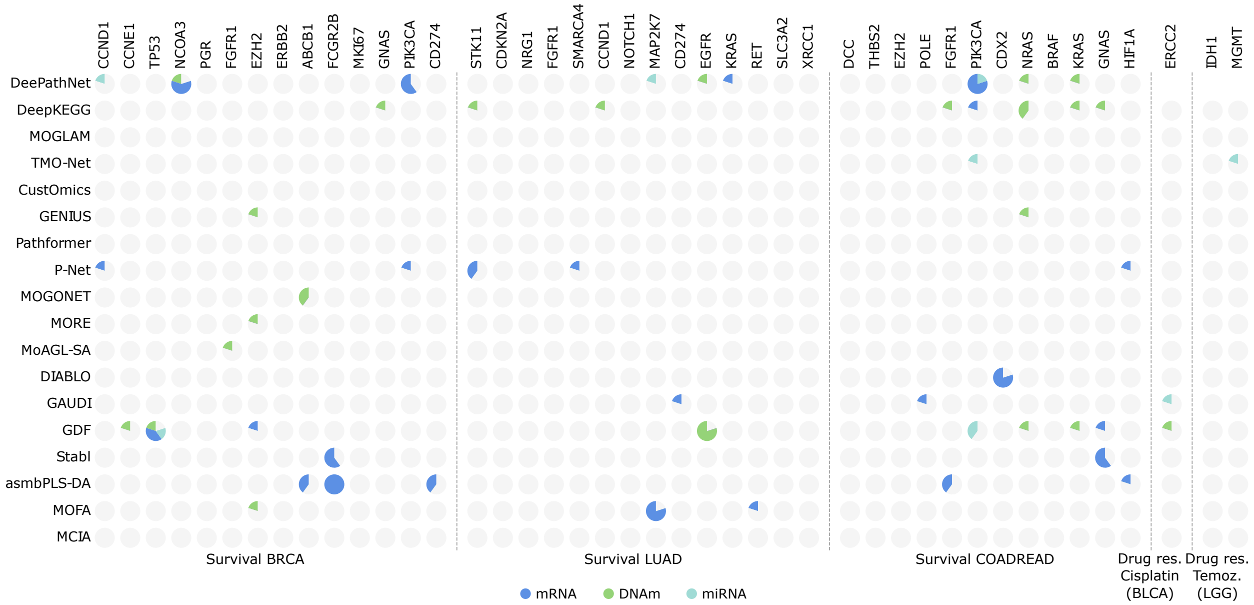

**Supplementary Figure 20: Summary of dominant omics types for biomarkers ranked in the top 1% from experiments with the mRNA+DNAm+miRNA omics combination.** Each pie has 5 sections corresponding to the five-fold cross-validation. A section has gray color if the biomarker is not ranked within top 1%.

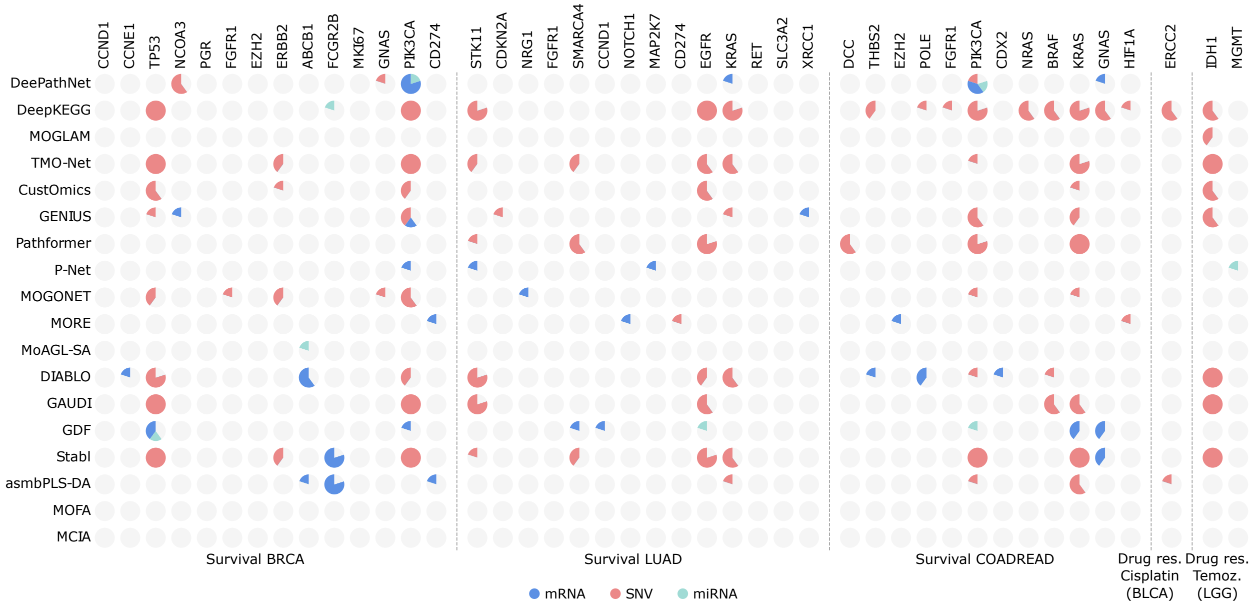

**Supplementary Figure 21: Summary of dominant omics types for biomarkers ranked in the top 1% from experiments with the mRNA+SNV+miRNA omics combination.** Each pie has 5 sections corresponding to the five-fold cross-validation. A section has gray color if the biomarker is not ranked within top 1%.

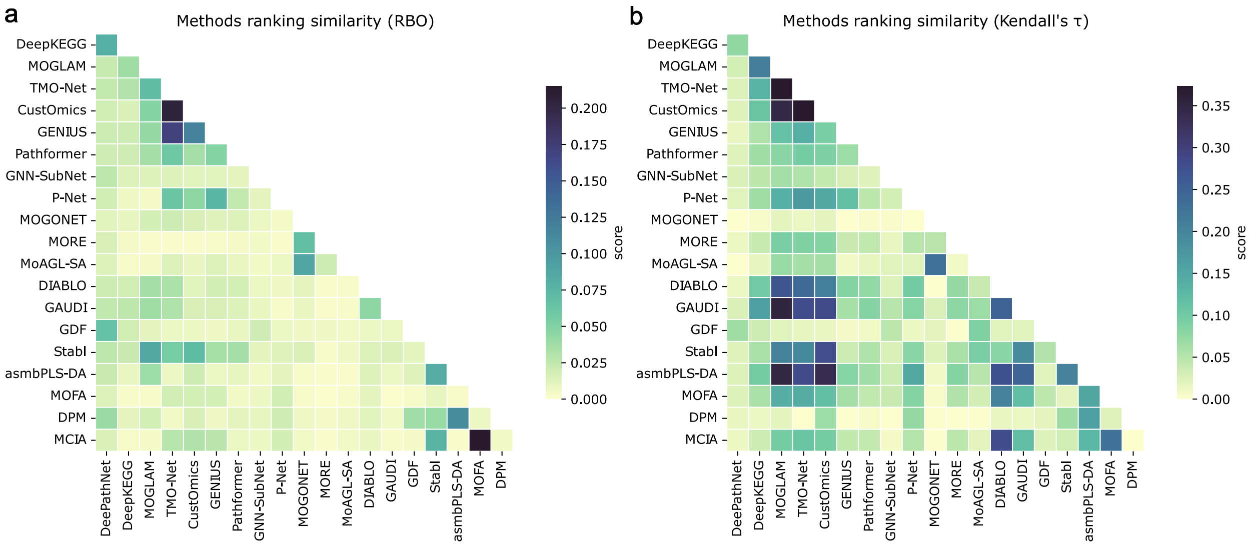

**Supplementary Figure 22: Similarity between the benchmarked methods.** Similarity is measured by the average RBO (**a**) and Kendall's $\tau$ (**b**) of the gene-level ranking lists between each two methods across all 30 experiments, including 6 omics combinations with 5 cross-validation folds.

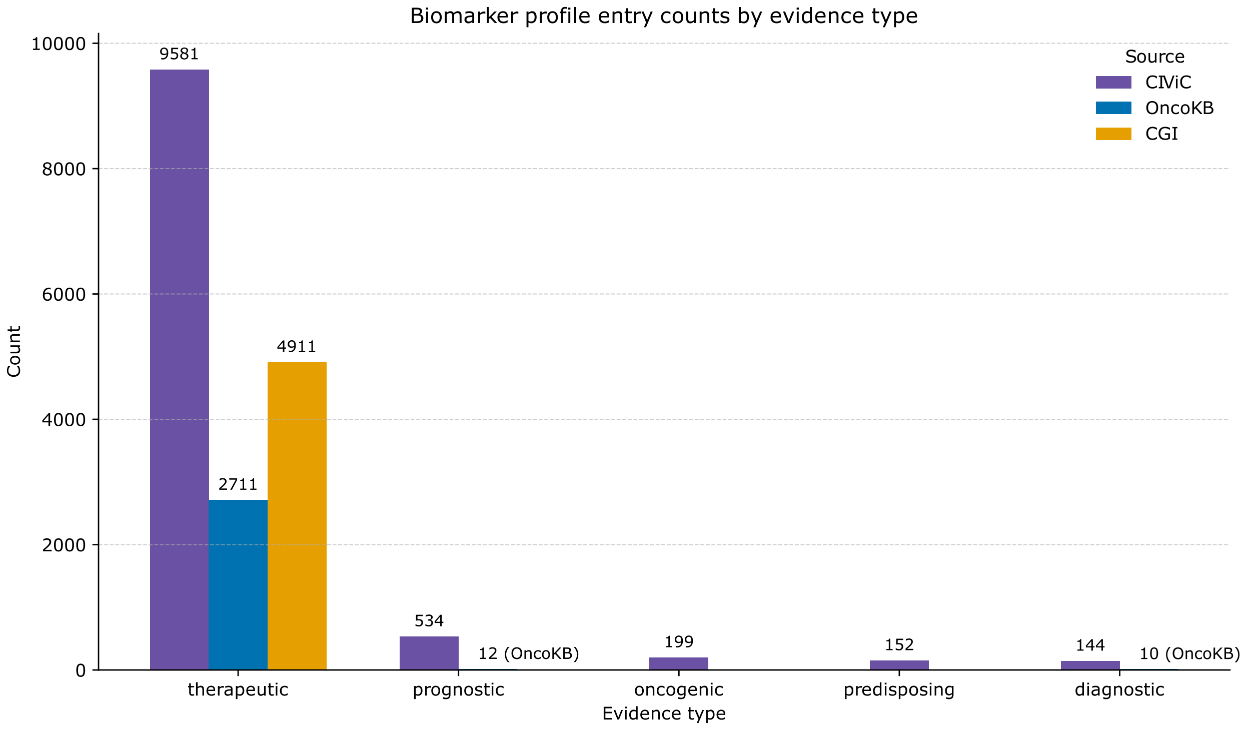

**Supplementary Figure 23:** Collected biomarker numbers by evidence type and source.

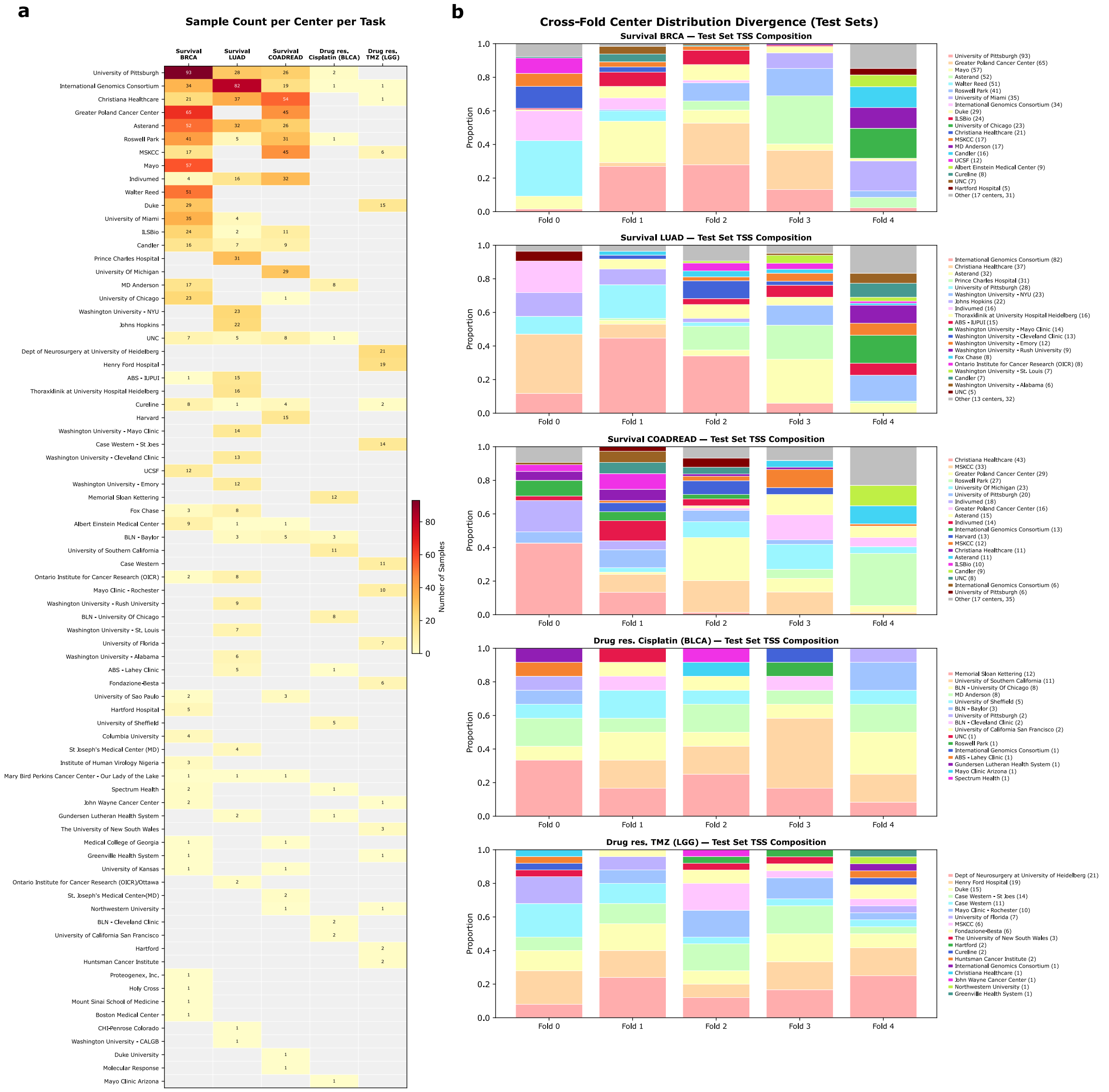

**Supplementary Figure 24: Tissue source site compositions within the task datasets.** **a**, For each task, the sample counts within each center are shown. **b**, For each task, the center proportions within the test sets are shown for each fold. TMZ: Temozolomide.

### 3 Supplementary Tables

**Supplementary Table 1:** Extended list of multi-omics biomarker identification methods. "-" indicates unavailability.

| Name | Year | Code | Selected | DOI |
|---|---|---|---|---|
| **DL** | | | | |
| Amogel | 2025 | Python | N | 10.1186/s12859-025-06111-6 |
| MOGAD | 2025 | Python | N | 10.3390/informatics12030068 |
| MOLUNGN | 2025 | Python | N | 10.3389/fgene.2025.1610284 |
| IGCN | 2025 | Python | N | 10.1093/bioinformatics/btaf313 |
| MULGONET | 2025 | Python | N | 10.1016/j.fmre.2025.01.004 |
| MOGKAN | 2025 | – | N | 10.1038/s41598-025-13337-0 |
| DOMSCNet | 2025 | R | N | 10.1093/bib/bbaf115 |
| GNNRAI | 2025 | – | N | 10.1038/s41540-025-00519-9 |
| DeePathNet | 2024 | Python | Y | 10.1158/2767-9764.CRC-24-0285 |
| DeepKEGG | 2024 | Python | Y | 10.1093/bib/bbae185 |
| TMO-Net | 2024 | Python | Y | 10.1186/s13059-024-03293-9 |
| Pathformer | 2024 | Python | Y | 10.1093/bioinformatics/btae316 |
| MoAGL-SA | 2024 | Python | Y | 10.1186/s12859-024-05989-y |
| MORE | 2024 | Python | Y | 10.1093/bib/bbae658 |
| Multilevel-GNN | 2024 | Python | N | 10.1093/bib/bbae184 |
| DeepMoIC | 2024 | – | N | 10.1186/s12864-024-11112-5 |
| Moiner | 2024 | Python | N | 10.1021/acs.jcim.4c00013 |
| HyperTMO | 2024 | Python | N | 10.1093/bioinformatics/btae159 |
| MoCAT | 2024 | Python | N | 10.1186/s13040-024-00360-6 |
| TEMINET | 2024 | Python | N | 10.3390/ijms25031655 |
| van Hilten et al., 2024 | 2024 | Python | N | 10.1038/s41540-024-00405-w |
| BioMGNN | 2024 | Python | N | 10.3390/biology13050338 |
| MOGDx | 2024 | R/Python | N | 10.1093/bioinformatics/btae523 |
| MCRGCN | 2024 | Python | N | 10.1007/s13755-024-00274-x |
| AUTOSurv | 2024 | Python | N | 10.1038/s41698-023-00494-6 |
| GENIUS | 2023 | Python | Y | 10.7554/eLife.87133.2 |
| MOGLAM | 2023 | Python | Y | 10.1016/j.compbiomed.2023.107303 |
| CustOmics | 2023 | Python | Y | 10.1371/journal.pcbi.1010921 |
| HTML | 2023 | Python | N | 10.1093/bib/bbad378 |
| GOAT | 2023 | Python | N | 10.1093/bioinformatics/btad582 |
| AGCN | 2023 | Python | N | 10.1093/bfgp/elad013 |
| GGNN | 2023 | Python | N | 10.1016/j.compbiomed.2023.107117 |
| DeepInsight-3D | 2023 | Matlab/Python | N | 10.1038/s41598-023-29644-3 |
| Zhuang et al., 2023 | 2023 | Python | N | 10.1371/journal.pone.0284563 |
| Demir Karaman et al., 2023 | 2023 | – | N | 10.3390/medsci11030044 |
| GNN-SubNet | 2022 | Python | Y | 10.1093/bioinformatics/btac478 |
| MoGCN | 2022 | Python | N | 10.3389/fgene.2022.806842 |
| Park et al., 2022 | 2022 | – | N | 10.3390/biom12121839 |
| MOMA | 2022 | Python | N | 10.1093/bioinformatics/btac080 |
| GTN | 2022 | Python | N | 10.1142/9789811250477_0034 |
| RDFS | 2022 | Python | N | 10.1016/j.eswa.2022.116813 |
| SDGCCA | 2022 | Python | N | 10.1089/cmb.2021.0598 |
| P-Net | 2021 | Python | Y | 10.1038/s41586-021-03922-4 |
| MOGONET | 2021 | Python | Y | 10.1038/s41467-021-23774-w |
| AutoGGN | 2021 | – | N | 10.1016/j.ailsci.2021.100019 |
| Subtype-GAN | 2021 | Python | N | 10.1093/bioinformatics/btab109 |
| Hira et al., 2021 | 2021 | Python | N | 10.1038/s41598-021-85285-4 |
| Varmole | 2020 | Python | N | 10.1093/bioinformatics/btaa866 |
| SALMON | 2019 | Python | N | 10.3389/fgene.2019.00166 |
| MiNet | 2019 | Python | N | 10.1007/978-3-030-20242-2_10 |
| Kim et al., 2018 | 2018 | Python | N | 10.3390/genes9100478 |
| **non-DL** | | | | |

Continued on next page

| | | | | |
|---|---|---|---|---|
| GAUDI | 2025 | R | Y | 10.1038/s41467-025-60822-1 |
| MTA-MO | 2025 | – | N | 10.1002/jcsm.13661 |
| Stabl | 2024 | Python | Y | 10.1038/s41587-023-02033-x |
| DPM | 2024 | R | Y | 10.1038/s41467-024-49986-4 |
| StabilityCCA | 2024 | – | N | 10.1371/journal.pone.0309921 |
| NetMIM | 2024 | R | N | 10.1093/bib/bbae454 |
| Hussein et al., 2024 | 2024 | R | N | 10.1038/s41540-024-00371-3 |
| jsCCA | 2024 | – | N | 10.1002/gepi.22566 |
| Tembhare et al., 2024 | 2024 | Python | N | 10.1016/j.imu.2024.101507 |
| asmbPLS-DA | 2023 | R | Y | 10.3390/genes14050961 |
| ioSearch | 2023 | – | N | 10.1002/gepi.22536 |
| MOBILE | 2023 | Matlab | N | 10.1038/s41467-023-39729-2 |
| GDF | 2022 | R | Y | 10.1038/s41598-022-21417-8 |
| DRAGON | 2022 | R/Python | N | 10.1093/nar/gkac1157 |
| QLattice | 2022 | Python | N | 10.1093/bioinformatics/btac405 |
| PIntMF | 2021 | R | N | 10.1093/bioinformatics/btab786 |
| TiMEG | 2021 | R | N | 10.1038/s41598-021-03034-z |
| ProMS | 2021 | Python | N | 10.1016/j.mcpro.2021.100083 |
| MAINE | 2021 | Web tool | N | 10.1093/bioinformatics/btab862 |
| KiMONo | 2021 | R | N | 10.1038/s41598-021-85544-4 |
| iDRW | 2021 | R | N | 10.1093/bioinformatics/btab086 |
| msPLS | 2020 | R | N | 10.1186/s12859-019-3286-3 |
| SMSPL | 2020 | R | N | 10.1109/TCYB.2020.3006240 |
| GARBO | 2020 | Python | N | 10.1093/bioinformatics/btaa144 |
| DIABLO | 2019 | R | Y | 10.1093/bioinformatics/bty1054 |
| LUCID | 2019 | R | N | 10.1093/bioinformatics/btz667 |
| MOFA | 2018 | R/Python | Y | 10.15252/msb.20178124 |
| IntLIM | 2018 | R | N | 10.1186/s12859-018-2085-6 |
| MRMR-mv | 2018 | Python | N | 10.1186/s12920-018-0388-0 |
| IPF-LASSO | 2017 | R | N | 10.1155/2017/7691937 |
| MCIA | 2014 | R | Y | 10.1186/1471-2105-15-162 |

**Supplementary Table 2:** Harmonization of biomarker evidence levels for different knowledge bases based on AMP/ASCO/CAP guidelines[40].

| **Evidence** | **AMP/ASCO/CAP description** | **CIViC** | **OncoKB** | **CGI** |
|---|---|---|---|---|
| Level A | Prognostic: Biomarkers included in professional guidelines as prognostic for a specific type of tumor.<br>Therapeutic: Biomarkers that predict response or resistance to FDA-approved therapies for a specific type of tumor; Biomarkers included in professional guidelines that predict response or resistance to therapies for a specific type of tumor. | A | Px1, Px2, R1, 1, 2 | FDA guidelines, NCCN guidelines, NCCN/CAP guidelines, CPIC guidelines, European LeukemiaNet guidelines |
| Level B | Prognostic: Biomarkers of prognostic significance for a specific type of tumor based on well-powered studies with consensus from experts in the field.<br>Therapeutic: Biomarkers that predict response or resistance to therapies for a specific type of tumor based on well-powered studies with consensus from experts in the field. | B | Px3, 3 | Late trials |
| Level C | Prognostic: Biomarkers of prognostic significance based on the results of multiple small studies.<br>Therapeutic: Biomarkers that predict response or resistance to therapies approved by the FDA or professional societies for a different type of tumor; Biomarkers that serve as inclusion criteria for clinical trials. | C | – | Early trials, Case report |
| Level D | Prognostic: Biomarkers that may assist disease prognosis themselves or along with other biomarkers based on small studies or a few case reports.<br>Therapeutic: Biomarkers that show plausible therapeutic significance based on preclinical studies. | D, E | R2, 4 | Pre-clinical |

**Supplementary Table 3:** Biomarkers for the real data tasks.

| Task | Gene | Level | Source | Omics | Direction | Key biological evidence |
|---|---|---|---|---|---|---|
| Survival BRCA | ABCB1 | B | CIViC | Mutation | Better | ABCB1 variants (G2677T, C3435T) associated with longer PFS in HER2+ mBC treated with taxane+trastuzumab. |
| | CCND1 | B | CIViC | Expression | Poor | Cyclin D1 overexpression linked to shorter OS and increased metastasis in ER+ breast cancer. |
| | CCNE1 | B | CIViC | Expression | Poor | High cyclin E expression strongly associated with poor prognosis (HR = 13.3 vs. normal levels). |
| | CD274 | B | CIViC | Expression | Mixed | PD-L1 overexpression correlates with shortened OS; however, PD-L1+ TIICs indicate better prognosis. |
| | ERBB2 | B | CIViC | Protein | Poor | Higher serum HER2 associated with worse PFS and OS in breast cancer (HR per 10 ng/mL: 1.024). |
| | EZH2 | B | CIViC | Expression | Poor | High EZH2 expression associated with poorer outcome (meta-analysis: 51 studies, 9444 patients). |
| | FCGR2B | B | CIViC | Mutation | Poor | FCGR2B I232T carriers did not benefit from trastuzumab, unlike I/I patients ($p = 0.03$ interaction). |
| | FGFR1 | B | CIViC | CNV, Expression | Poor | FGFR1 amplification/expression predicts worse OS (meta-analysis) and poor DFS in luminal A subtype. |
| | GNAS | B | CIViC | Mutation | Poor | GNAS T393C TT genotype associated with higher death risk; 10-yr survival 63% (CC) vs. 23% (TT). |
| | MKI67 | B | CIViC | Expression | Poor | Ki-67 $\geq$25% by IHC prognostic for worse OS (meta-analysis HR = 2.05). |
| | NCOA3 | B | CIViC | CNV, Expression | Poor | NCOA3 amplification and high expression both associated with shorter disease-specific survival. |
| | PGR | B | CIViC | Expression | Mixed | PgR expression predicts better relapse-free survival; low expression predicts worse DFS. |
| | PIK3CA | B | CIViC | Mutation | Mixed | H1047R mutations predict longer OS; exon 9 mutations predict worse OS and DFS; overall effect context-dependent. |
| | TP53 | B | CIViC | Mutation | Poor | Multiple hotspot mutations (R248W worst) predict poor OS; DNA-contact region mutations worsen RFS. |
| Survival LUAD | CCND1 | B | CIViC | CNV, Expression | Poor | Increased CCND1 copy number and expression associated with poorer OS in NSCLC. |
| | CD274 | B | CIViC | Expression | Mixed | PD-L1+/CD8-low tumors show shortest PFS; PD-L1+ TIICs may indicate better prognosis. |
| | CDKN2A | B | CIViC | Methylation, Expression | Poor | p16 promoter hypermethylation and low p16 protein linked to shorter recurrence time and OS in NSCLC. |
| | EGFR | B | CIViC | Mutation | Mixed | L858R predicts better OS; T790M predicts worse PFS and OS; activating mutations overall protective. |
| | FGFR1 | B | CIViC | CNV | Mixed | Moderate FGFR1 copy number (4–6) reduces death risk; high amplification worsens OS (meta-analysis). |
| | KRAS | B | CIViC | Mutation | Poor | KRAS mutations (esp. G12V, G12C) associated with worse OS, PFS, and higher recurrence in NSCLC. |
| | MAP2K7 | B | CIViC | Mutation | Poor | E116K genotype reduces median survival by 4/7 months; increases cancer death risk (HR up to 1.94). |
| | NOTCH1 | B | CIViC | Mutation | Poor | Gain-of-function mutations (D1642H, R2327W, etc.) correlate with poor prognosis in TP53-wt lung cancer. |
| | NRG1 | B | CIViC | Fusion | Poor | SLC3A2–NRG1 fusion associated with inferior survival in mucinous lung adenocarcinoma. |
| | RET | B | CIViC | Expression | Poor | High RET mRNA expression correlates with shorter OS in ASCL1-expressing lung adenocarcinoma. |
| | SLC3A2 | B | CIViC | Fusion | Poor | SLC3A2–NRG1 fusion-positive patients demonstrate inferior OS in lung adenocarcinoma. |
| | SMARCA4 | B | CIViC | Mutation, Expression | Poor | Loss-of-function mutations and low expression both predict worse OS in lung adenocarcinoma. |

*Continued on next page*

*Table 3 continued*

| Task | Gene | Level | Source | Omics | Direction | Key biological evidence |
|---|---|---|---|---|---|---|
| | STK11 | B | CIViC | Mutation | Poor | Exon 1–2 mutations significantly shorten OS (24 vs. 69 months) in non-squamous NSCLC. |
| | XRCC1 | B | CIViC | Mutation | Better | R399Q variant correlates with higher OS in NSCLC patients treated with gemcitabine+platinum. |
| Survival COADREAD | BRAF | B | CIViC | Mutation | Mixed | V600E predicts poor OS (HR ∼ 2–5×); non-V600 mutations show longer survival than V600E and wt. |
| | CDX2 | B | CIViC | Expression | Better | CDX2 loss predicts lower 5-yr DFS in stage II/III CRC (HR = 3.44 discovery; 2.42 validation). |
| | DCC | B | CIViC | Expression | Better | DCC expression predicts better 5-yr survival (94% vs. 62% stage II; 59% vs. 33% stage III). |
| | EZH2 | B | CIViC | Mutation, Expression | Poor | Intron 6 variant and high expression both correlate with lower PFS and OS in metastatic CRC. |
| | FGFR1 | B | CIViC | CNV | Poor | FGFR1 amplification predicts worse OS across cancer types (meta-analysis). |
| | GNAS | B | CIViC | Mutation | Better | GNAS T393C TT genotype shows higher 5-yr survival (88%) vs. CC (50%) in stage I–II CRC. |
| | HIF1A | B | CIViC | Expression | Poor | HIF1A overexpression linked to higher CRC-specific mortality (adjusted HR = 1.72, $p = 0.0007$). |
| | KRAS | B | CIViC | Mutation | Mixed | G12/G13 mutations reduce PFS and OS; however, G12D shows longer survival than other G12 variants. |
| | NRAS | B | CIViC | Mutation | Poor | NRAS mutations associated with poorer survival and worse prognosis in CRC. |
| | PIK3CA | B | CIViC | Mutation | Poor | Mutations reduce relapse-free survival; E545K especially associated with high recurrence (89%). |
| | POLE | B | CIViC | Mutation | Better | Proofreading domain mutations (P286R, V411L, S459F) identify immunogenic CRCs with excellent prognosis. |
| | THBS2 | B | CIViC | Expression | Poor | Low THBS2 expression is a negative prognostic factor for DFS (HR = 3.057, $p = 0.002$). |
| Drug Cisplatin (BLCA) | ERCC2 | B | OncoKB | Mutation | Sensitivity | Oncogenic ERCC2 mutations predict sensitivity to cisplatin-based chemotherapy in bladder cancer. |
| Drug Temozolomide (LGG) | IDH1 | B | CIViC | Mutation | Sensitivity | IDH mutations improve temozolomide response rate (61% vs. 17%, $p = 0.01$) in low-grade glioma. |
| | MGMT | B | CIViC | Expression | Sensitivity | Low MGMT protein expression associated with objective response to temozolomide ($p < 0.04$). |

**Notes.** Source refers to the knowledge base. CNV = copy number alteration (amplification or deletion). Direction indicates the prognostic or predictive association: Poor = associated with worse prognosis; Better = associated with improved prognosis; Mixed = direction depends on the specific variant or context; Sensitivity = the alteration predicts response to the indicated drug.

**Supplementary Table 4:** Overview of the tasks and corresponding omics combinations of the benchmarked methods. Only cancer-related, sample-level prediction tasks using bulk-level multi-omics data are included, and tasks with cell lines, single-cell omics, or single omics data are excluded. "–" indicates no satisfied tasks.

| Name | Tasks | Omics combinations |
|---|---|---|
| **DL** | | |
| P-Net [14] | Cancer metastasis prediction | CNV, SNV |
| | Cancer metastasis prediction (with mRNA-seq derived fusion) | mRNA, CNV, SNV |
| GENIUS [17] | Cancer type classification (TCGA) | mRNA, DNAm, CNV, SNV |
| | Cancer stage prediction (TCGA) | mRNA, DNAm, CNV, SNV |
| TMO-Net [7] | Breast cancer subtype classification (TCGA) | mRNA, CNV, SNV |
| | Breast cancer subtype classification (METABRIC) | mRNA, SNV |
| | Cancer metastasis prediction (TCGA) | mRNA, DNAm |
| | Cancer survival prediction (TCGA) | mRNA, DNAm, CNV, SNV |
| | Cancer survival prediction (CPTAC) | mRNA, DNAm, CNV, SNV, Protein |
| CustOmics [19] | Cancer survival prediction (TCGA) | mRNA, DNAm, CNV |
| | cancer type classification (TCGA) | mRNA, DNAm, CNV |
| | Breast cancer subtype classification (TCGA) | mRNA, DNAm, CNV |
| MOGONET [21] | Breast cancer subtype classification (TCGA) | mRNA, miRNA, DNAm |
| | LGG grade classification (TCGA) | mRNA, miRNA, DNAm |
| MoAGL-SA [22] | Breast cancer subtype classification (TCGA) | mRNA, miRNA, DNAm |
| | Cancer stage prediction (TCGA) | mRNA, miRNA, DNAm |
| MORE [23] | Breast cancer subtype classification (TCGA) | mRNA, miRNA, DNAm |
| | Glioblastoma subtype classification (TCGA) | mRNA, miRNA, DNAm |
| MOGLAM [24] | Cancer type classification (TCGA) | mRNA, miRNA, DNAm |
| | Breast cancer classification (TCGA) | mRNA, miRNA, DNAm |
| GNN-SubNet [25] | Cancer type classification (TCGA) | mRNA, DNAm |
| Pathformer [5] | Cancer survival prediction (TCGA) | mRNA, DNAm, CNV |
| | Cancer stage prediction (TCGA) | mRNA, DNAm, CNV |
| | Cancer drug response prediction (TCGA) | mRNA, DNAm, CNV |
| DeePathNet [6] | Cancer type classification (TCGA) | mRNA, CNV, SNV |
| | Breast cancer subtype classification (TCGA for train, CPTAC for test) | mRNA, CNV, SNV |
| DeepKEGG [29] | Cancer recurrence prediction (TCGA) | mRNA, miRNA, SNV |
| | Cancer recurrence prediction (TARGETS) | mRNA, miRNA |
| **Statistical and ML** | | |
| MCIA [31] | - | - |
| MOFA [32] | (Unsupervised) multi-omics integration (a chronic lymphocytic leukaemia cohort) | mRNA, DNAm, SNV |
| GAUDI [8] | Cancer survival prediction (TCGA) | mRNA, miRNA, DNAm |
| | Clinical variable associations (TCGA) | mRNA, miRNA, DNAm |
| DIABLO [9] | Breast cancer subtype classification (TCGA) | mRNA, miRNA, DNAm, Protein |
| | Cancer survival prediction (TCGA) | mRNA, miRNA, DNAm |
| asmbPLS-DA [35] | Breast cancer tumor vs normal classification (TCGA) | mRNA, miRNA |
| | Breast cancer stage prediction (TCGA) | mRNA, miRNA |
| | Breast cancer stage prediction (TCGA) | mRNA, miRNA, Protein |
| Stabl [36] | - | - |
| GDF [38] | Cancer survival prediction (TCGA) | mRNA, DNAm |
| | Cancer type classification (TCGA) | mRNA, DNAm |
| DPM [39] | Ovarian cancer survival analysis (CPTAC-3 and TCGA) | mRNA, Protein (CPTAC-3), clinical covariates |
| | Glioma IDH status prediction (mutant vs wild type) (TCGA and CPTAC-3/ProteomeXchange) | mRNA, DNAm, Protein |

**Supplementary Table 5:** Summary of key statistics of the simulated datasets used in this study.

| Signal strength | Sample size | | #Feature | | | #Biomarker |
|---|---|---|---|---|---|---|
| | Class1 | Class2 | DNAm | mRNA | Protein | |
| $\delta = 0.5$ | 50 | 50 | 6 273 | 3 862 | 95 | 138 |
| $\delta = 1.0$ | 50 | 50 | 6 273 | 3 862 | 95 | 147 |
| $\delta = 2.0$ | 50 | 50 | 6 273 | 3 862 | 95 | 169 |
| $\delta = 3.0$ | 50 | 50 | 6 273 | 3 862 | 95 | 123 |
| $\delta = 4.0$ | 50 | 50 | 6 273 | 3 862 | 95 | 168 |
| $\delta = 5.0$ | 50 | 50 | 6 273 | 3 862 | 95 | 150 |

**Supplementary Table 6:** Sample size and collected biomarker number for each TCGA project. COAD and READ are combined into COADREAD. Samples with a missing omics type among mRNA, miRNA, CNV, SNV, or DNA methylation are not counted. Biomarker numbers are directly recorded from collected gold-reference sets before the manual inspection filtering step (see Methods in the main text).

| TCGA Project | # Sample (all five omics types) | # Biomarker (before manual inspection) |
|---|---|---|
| BRCA | 647 | 14 |
| LGG | 489 | 6 |
| HNSC | 478 | 12 |
| THCA | 475 | 7 |
| PRAD | 471 | 9 |
| SKCM | 430 | 8 |
| LUAD | 423 | 21 |
| UCEC | 381 | 7 |
| BLCA | 377 | 6 |
| COADREAD | 371 | 15 |
| LIHC | 350 | 5 |
| STAD | 345 | 9 |
| LUSC | 341 | 16 |
| CESC | 274 | 5 |
| KIRC | 258 | 9 |
| KIRP | 253 | 6 |
| SARC | 227 | 7 |
| ESCA | 179 | 8 |
| PCPG | 173 | 3 |
| PAAD | 166 | 9 |
| TGCT | 124 | 3 |
| THYM | 118 | 3 |
| LAML | 95 | 23 |
| MESO | 78 | 4 |
| ACC | 75 | 4 |
| UVM | 75 | 4 |
| KICH | 64 | 6 |
| UCS | 53 | 3 |
| CHOL | 35 | 4 |
| DLBC | 34 | 5 |
| OV | 9 | 10 |

**Supplementary Table 7:** Excluded biomarkers after manual inspection for the candidate tasks. Biomarkers, the corresponding tasks, exclusion rationales, and record links are listed.

| Biomarker | Task | Exclusion rationale | Record link |
|---|---|---|---|
| TP53 | LUSC/LUAD survival | No prognosis effects. | civicdb.org/links/evidence_items/2999 |
| STK11 | LUSC survival | Not for lung squamous cell carcinoma. | civicdb.org/links/evidence_items/750 |
| RB1 | LUSC/LUAD survival | Low rating. | civicdb.org/links/evidence_items/1313 |
| PIM1 | LUSC/LUAD survival | Low rating. | civicdb.org/links/evidence_items/1168 |
| CBLB | LUSC/LUAD survival | Low rating. | civicdb.org/links/evidence_items/1637 |
| EZH2 | LUSC/LUAD survival | Non-significant result with lung cancer according to the source publication. | civicdb.org/links/evidence_items/731 |
| ACTA1 | LUAD survival | Low rating. | civicdb.org/links/evidence_items/1161 |
| MAP2K1 | LUAD survival | Not established as a prognostic biomarker according to the source publication. | civicdb.org/links/evidence_items/2935 |
| CD274 | COADREAD survival | Low rating. | civicdb.org/links/evidence_items/4856 |
| CD274 | COADREAD/HNSC survival | Non-significant prognostic effects. | civicdb.org/links/evidence_items/5507 |
| NOTCH1 | COADREAD survival | Low rating. | civicdb.org/links/evidence_items/812 |
| MIR218-1 | COADREAD survival | Non-protein-coding; not present in our multi-omics gene set. | civicdb.org/links/evidence_items/1119 |
| PTPN11 | HNSC survival | Low rating. | civicdb.org/links/evidence_items/1316 |
| MDM2 | HNSC survival | Low rating. | civicdb.org/links/evidence_items/1169 |

**Supplementary Table 8:** Number of responders:non-responders with biomarker (AMP/ASCO/CAP[40] Tier I evidence) counts in parentheses. Sample sizes are from TCGA clinical data and the missingness of omics types are not accounted for, thus the actual sample size with all five omics types may be smaller. "-" indicates the drug-cancer-type-pair does not have a responder, non-responder, or biomarker. Drugs or cancer types with all entries "-" are omitted from this table.

| Drug | BLCA | BRCA | COADREAD | GBM | HNSC | LGG | LIHC | LUAD | LUSC | OV | PAAD | PRAD | SKCM | STAD | UCEC |
|---|---|---|---|---|---|---|---|---|---|---|---|---|---|---|---|
| Bevacizumab | - | 2:2 (2) | 8:17 (2) | - | - | 3:20 (1) | - | - | - | - | - | - | - | - | - |
| Cetuximab | - | - | 2:4 (11) | - | - | - | - | - | - | - | - | - | - | - | - |
| Cisplatin | 40:23 (1) | - | - | - | 45:5 (1) | - | - | 41:11 (1) | 27:5 (1) | - | - | - | - | 23:15 (1) | - |
| Dabrafenib | - | - | - | - | - | - | - | - | - | - | - | - | 1:2 (1) | - | - |
| Docetaxel | - | - | - | - | - | - | - | - | - | - | - | 1:2 (1) | - | 1:3 (1) | - |
| Doxorubicin | - | 88:8 (2) | - | - | - | - | - | - | - | - | - | - | - | - | - |
| Epirubicin | - | 24:1 (1) | - | - | - | - | - | - | - | - | - | - | - | - | - |
| Erlotinib | - | - | - | - | - | - | - | 1:5 (2) | - | - | - | - | - | - | - |
| Exemestane | - | 4:2 (1) | - | - | - | - | - | - | - | - | - | - | - | - | - |
| Fluorouracil | - | - | 72:29 (4) | - | - | - | - | - | - | - | - | - | - | 60:32 (1) | - |
| Gemcitabine | - | - | - | - | - | - | - | 6:3 (1) | 9:8 (1) | - | 29:42 (2) | - | - | - | - |
| Irinotecan | - | - | 8:15 (2) | - | - | - | - | - | - | - | - | - | - | - | - |
| Methotrexate | 5:5 (2) | - | - | - | - | - | - | - | - | - | - | - | - | - | - |
| Oxaliplatin | - | - | 52:20 (1) | - | - | - | - | - | - | - | - | - | - | 15:8 (1) | - |
| Paclitaxel | - | 54:7 (1) | - | - | - | - | - | 14:12 (1) | 5:5 (1) | 5:1 (1) | - | - | - | - | 44:8 (1) |
| Pemetrexed | - | - | - | - | - | - | - | 18:12 (2) | - | - | - | - | - | - | - |
| Sorafenib | - | - | - | - | - | - | 3:15 (1) | - | - | - | - | - | - | - | - |
| Tamoxifen | - | 15:7 (4) | - | - | - | - | - | - | - | - | - | - | - | - | - |
| Temozolomide | - | - | - | 2:10 (2) | - | 21:106 (2) | - | - | - | - | - | - | - | - | - |
| Trametinib | - | - | - | - | - | - | - | - | - | - | - | - | 1:1 (2) | - | - |
| Trastuzumab | - | 16:1 (5) | - | - | - | - | - | - | - | - | - | - | - | - | - |

**Supplementary Table 9:** Classifiability test on candidate tasks with random forest (RF) and support vector machine (SVM). AUROC is used as metric. Mean and standard deviation across five-fold are reported. Results below the random baseline are underscored.

| | CNV+DNAm+mRNA | | CNV+SNV+mRNA | | CNV+mRNA+miRNA | | DNAm+SNV+mRNA | | DNAm+mRNA+miRNA | | SNV+mRNA+miRNA | |
|---|---|---|---|---|---|---|---|---|---|---|---|---|
| | RF | SVM | RF | SVM | RF | SVM | RF | SVM | RF | SVM | RF | SVM |
| Survival BRCA | 0.62 (0.06) | 0.64 (0.08) | 0.61 (0.06) | 0.63 (0.08) | 0.60 (0.07) | 0.63 (0.09) | 0.62 (0.05) | 0.64 (0.08) | 0.63 (0.07) | 0.64 (0.09) | 0.61 (0.06) | 0.62 (0.08) |
| Survival LUAD | 0.57 (0.06) | 0.60 (0.06) | 0.55 (0.04) | 0.56 (0.05) | 0.55 (0.06) | 0.55 (0.04) | 0.53 (0.08) | 0.60 (0.09) | 0.57 (0.03) | 0.60 (0.09) | 0.55 (0.07) | 0.54 (0.05) |
| Survival LUSC | 0.45 (0.10) | 0.52 (0.03) | 0.53 (0.08) | 0.51 (0.07) | 0.53 (0.12) | 0.51 (0.07) | 0.52 (0.07) | 0.53 (0.04) | 0.48 (0.09) | 0.54 (0.04) | 0.54 (0.12) | 0.51 (0.04) |
| Survival COADREAD | 0.59 (0.05) | 0.57 (0.05) | 0.60 (0.04) | 0.55 (0.05) | 0.60 (0.07) | 0.55 (0.05) | 0.55 (0.06) | 0.59 (0.06) | 0.56 (0.07) | 0.58 (0.05) | 0.58 (0.05) | 0.55 (0.04) |
| Cisplatin (BLCA) | 0.60 (0.15) | 0.60 (0.03) | 0.67 (0.09) | 0.56 (0.16) | 0.63 (0.11) | 0.58 (0.15) | 0.54 (0.12) | 0.61 (0.06) | 0.57 (0.04) | 0.61 (0.06) | 0.50 (0.14) | 0.65 (0.18) |
| Fluorouracil (STAD) | 0.49 (0.15) | 0.50 (0.23) | 0.34 (0.14) | 0.39 (0.13) | 0.40 (0.18) | 0.37 (0.15) | 0.49 (0.15) | 0.50 (0.18) | 0.52 (0.21) | 0.52 (0.19) | 0.39 (0.17) | 0.40 (0.12) |
| Gemcitabine (PAAD) | 0.45 (0.14) | 0.56 (0.14) | 0.48 (0.11) | 0.55 (0.10) | 0.48 (0.12) | 0.54 (0.11) | 0.50 (0.15) | 0.54 (0.18) | 0.57 (0.06) | 0.52 (0.16) | 0.56 (0.10) | 0.52 (0.12) |
| Temozolomide (LGG) | 0.58 (0.13) | 0.62 (0.17) | 0.54 (0.17) | 0.60 (0.19) | 0.53 (0.16) | 0.60 (0.19) | 0.56 (0.12) | 0.61 (0.17) | 0.54 (0.16) | 0.61 (0.16) | 0.52 (0.19) | 0.59 (0.20) |

**Supplementary Table 10:** Prediction performance for the BRCA survival task. Mean and standard deviation across five-fold are reported. The following methods are excluded due to a lack of sample-level label prediction module: MOFA, MCIA, GAUDI, and DPM.

| Model | CNV+DNAm+mRNA | | CNV+SNV+mRNA | | CNV+mRNA+miRNA | | DNAm+SNV+mRNA | | DNAm+mRNA+miRNA | | SNV+mRNA+miRNA | |
|---|---|---|---|---|---|---|---|---|---|---|---|---|
| | AUPR | AUROC | AUPR | AUROC | AUPR | AUROC | AUPR | AUROC | AUPR | AUROC | AUPR | AUROC |
| DeePathNet | 0.59 (0.06) | 0.58 (0.07) | 0.57 (0.07) | 0.56 (0.07) | 0.60 (0.08) | 0.59 (0.07) | 0.59 (0.06) | 0.58 (0.05) | 0.58 (0.07) | 0.57 (0.05) | 0.58 (0.06) | 0.57 (0.04) |
| DeepKEGG | 0.62 (0.08) | 0.61 (0.09) | 0.63 (0.10) | 0.62 (0.09) | 0.66 (0.04) | 0.66 (0.05) | 0.63 (0.08) | 0.65 (0.07) | 0.66 (0.08) | 0.64 (0.08) | 0.61 (0.06) | 0.61 (0.05) |
| MOGLAM | 0.63 (0.09) | 0.61 (0.09) | 0.51 (0.04) | 0.49 (0.05) | 0.62 (0.05) | 0.60 (0.07) | 0.59 (0.09) | 0.59 (0.08) | 0.61 (0.06) | 0.58 (0.06) | 0.59 (0.06) | 0.57 (0.08) |
| TMO-Net | 0.55 (0.02) | 0.55 (0.03) | 0.58 (0.05) | 0.56 (0.06) | 0.57 (0.04) | 0.57 (0.05) | 0.55 (0.07) | 0.55 (0.09) | 0.56 (0.05) | 0.58 (0.06) | 0.61 (0.04) | 0.60 (0.07) |
| CustOmics | 0.68 (0.07) | 0.67 (0.07) | 0.59 (0.06) | 0.59 (0.06) | 0.63 (0.06) | 0.63 (0.06) | 0.63 (0.09) | 0.62 (0.08) | 0.65 (0.10) | 0.64 (0.08) | 0.62 (0.06) | 0.60 (0.06) |
| GENIUS | 0.59 (0.05) | 0.59 (0.05) | 0.56 (0.05) | 0.56 (0.06) | 0.59 (0.05) | 0.61 (0.07) | 0.59 (0.05) | 0.61 (0.07) | 0.61 (0.09) | 0.63 (0.10) | 0.58 (0.05) | 0.58 (0.06) |
| Pathformer | 0.58 (0.06) | 0.59 (0.05) | 0.59 (0.04) | 0.60 (0.04) | 0.58 (0.08) | 0.57 (0.08) | 0.60 (0.06) | 0.60 (0.07) | 0.64 (0.08) | 0.64 (0.06) | 0.64 (0.08) | 0.63 (0.07) |
| GNN-SubNet | 0.52 (0.02) | 0.49 (0.03) | 0.54 (0.04) | 0.52 (0.05) | 0.50 (0.04) | 0.49 (0.06) | 0.53 (0.05) | 0.50 (0.05) | 0.54 (0.05) | 0.51 (0.07) | 0.50 (0.07) | 0.46 (0.08) |
| P-Net | 0.59 (0.05) | 0.58 (0.07) | 0.58 (0.04) | 0.57 (0.04) | 0.60 (0.05) | 0.59 (0.05) | 0.60 (0.05) | 0.59 (0.06) | 0.60 (0.06) | 0.59 (0.06) | 0.61 (0.07) | 0.59 (0.06) |
| MOGONET | 0.60 (0.11) | 0.56 (0.11) | 0.54 (0.06) | 0.53 (0.06) | 0.54 (0.06) | 0.53 (0.07) | 0.49 (0.03) | 0.46 (0.05) | 0.55 (0.06) | 0.51 (0.06) | 0.51 (0.07) | 0.48 (0.10) |
| MORE | 0.59 (0.08) | 0.58 (0.08) | 0.57 (0.05) | 0.54 (0.07) | 0.58 (0.06) | 0.58 (0.06) | 0.61 (0.06) | 0.61 (0.05) | 0.56 (0.05) | 0.53 (0.06) | 0.54 (0.03) | 0.52 (0.05) |
| MoAGL-SA | 0.52 (0.01) | 0.49 (0.03) | 0.50 (0.06) | 0.50 (0.08) | 0.55 (0.08) | 0.54 (0.08) | 0.48 (0.05) | 0.46 (0.07) | 0.54 (0.08) | 0.50 (0.09) | 0.52 (0.02) | 0.50 (0.01) |
| DIABLO | 0.57 (0.07) | 0.55 (0.07) | 0.52 (0.03) | 0.50 (0.04) | 0.54 (0.09) | 0.53 (0.09) | 0.55 (0.05) | 0.56 (0.07) | 0.57 (0.07) | 0.58 (0.09) | 0.59 (0.04) | 0.58 (0.04) |
| GDF | 0.62 (0.06) | 0.64 (0.05) | 0.59 (0.04) | 0.60 (0.03) | 0.63 (0.07) | 0.64 (0.06) | 0.61 (0.06) | 0.63 (0.05) | 0.61 (0.06) | 0.64 (0.06) | 0.64 (0.04) | 0.64 (0.06) |
| Stabl | 0.56 (0.05) | 0.54 (0.07) | 0.51 (0.05) | 0.50 (0.08) | 0.54 (0.03) | 0.56 (0.01) | 0.53 (0.07) | 0.50 (0.07) | 0.55 (0.05) | 0.54 (0.07) | 0.53 (0.04) | 0.53 (0.05) |
| asmbPLS-DA | 0.62 (0.09) | 0.62 (0.08) | 0.60 (0.07) | 0.59 (0.07) | 0.62 (0.06) | 0.62 (0.06) | 0.65 (0.09) | 0.64 (0.08) | 0.65 (0.09) | 0.65 (0.08) | 0.66 (0.08) | 0.65 (0.07) |

**Supplementary Table 11:** Prediction performance for the LUAD survival task. Mean and standard deviation across five-fold are reported. The following methods are excluded due to a lack of sample-level label prediction module: MOFA, MCIA, GAUDI, and DPM.

| Model | CNV+DNAm+mRNA | | CNV+SNV+mRNA | | CNV+mRNA+miRNA | | DNAm+SNV+mRNA | | DNAm+mRNA+miRNA | | SNV+mRNA+miRNA | |
|---|---|---|---|---|---|---|---|---|---|---|---|---|
| | AUPR | AUROC | AUPR | AUROC | AUPR | AUROC | AUPR | AUROC | AUPR | AUROC | AUPR | AUROC |
| DeePathNet | 0.58 (0.04) | 0.54 (0.06) | 0.58 (0.03) | 0.55 (0.05) | 0.56 (0.03) | 0.54 (0.04) | 0.56 (0.02) | 0.55 (0.04) | 0.55 (0.02) | 0.54 (0.04) | 0.54 (0.02) | 0.55 (0.04) |
| DeepKEGG | 0.57 (0.07) | 0.55 (0.08) | 0.55 (0.07) | 0.54 (0.10) | 0.56 (0.02) | 0.52 (0.04) | 0.60 (0.06) | 0.60 (0.07) | 0.54 (0.06) | 0.55 (0.07) | 0.54 (0.04) | 0.51 (0.03) |
| MOGLAM | 0.57 (0.05) | 0.55 (0.05) | 0.54 (0.04) | 0.54 (0.05) | 0.56 (0.05) | 0.53 (0.05) | 0.60 (0.08) | 0.55 (0.09) | 0.58 (0.04) | 0.56 (0.03) | 0.53 (0.06) | 0.50 (0.07) |
| TMO-Net | 0.57 (0.04) | 0.57 (0.05) | 0.54 (0.05) | 0.54 (0.05) | 0.50 (0.04) | 0.50 (0.07) | 0.56 (0.06) | 0.58 (0.06) | 0.54 (0.04) | 0.56 (0.06) | 0.53 (0.08) | 0.54 (0.08) |
| CustOmics | 0.53 (0.04) | 0.53 (0.06) | 0.52 (0.03) | 0.53 (0.04) | 0.57 (0.03) | 0.58 (0.03) | 0.55 (0.07) | 0.53 (0.11) | 0.54 (0.05) | 0.55 (0.06) | 0.54 (0.04) | 0.52 (0.05) |
| GENIUS | 0.57 (0.04) | 0.58 (0.05) | 0.56 (0.03) | 0.55 (0.04) | 0.56 (0.06) | 0.55 (0.05) | 0.54 (0.04) | 0.54 (0.03) | 0.55 (0.02) | 0.55 (0.02) | 0.55 (0.05) | 0.55 (0.04) |
| Pathformer | 0.54 (0.03) | 0.55 (0.05) | 0.52 (0.03) | 0.51 (0.05) | 0.56 (0.04) | 0.54 (0.06) | 0.55 (0.02) | 0.56 (0.03) | 0.52 (0.06) | 0.51 (0.06) | 0.55 (0.04) | 0.54 (0.05) |
| GNN-SubNet | 0.52 (0.05) | 0.49 (0.08) | 0.56 (0.07) | 0.54 (0.08) | 0.50 (0.04) | 0.47 (0.04) | 0.53 (0.03) | 0.52 (0.03) | 0.52 (0.03) | 0.52 (0.07) | 0.52 (0.04) | 0.50 (0.05) |
| P-Net | 0.56 (0.06) | 0.54 (0.07) | 0.53 (0.02) | 0.52 (0.04) | 0.54 (0.04) | 0.53 (0.05) | 0.55 (0.06) | 0.55 (0.07) | 0.54 (0.07) | 0.55 (0.07) | 0.53 (0.04) | 0.53 (0.04) |
| MOGONET | 0.56 (0.06) | 0.54 (0.08) | 0.56 (0.06) | 0.53 (0.07) | 0.54 (0.03) | 0.51 (0.06) | 0.50 (0.05) | 0.48 (0.08) | 0.53 (0.06) | 0.51 (0.05) | 0.51 (0.05) | 0.46 (0.06) |
| MORE | 0.56 (0.09) | 0.53 (0.08) | 0.55 (0.04) | 0.53 (0.04) | 0.55 (0.06) | 0.53 (0.05) | 0.54 (0.07) | 0.51 (0.10) | 0.52 (0.06) | 0.49 (0.07) | 0.54 (0.06) | 0.52 (0.07) |
| MoAGL-SA | 0.52 (0.05) | 0.48 (0.07) | 0.51 (0.04) | 0.48 (0.03) | 0.56 (0.06) | 0.51 (0.05) | 0.52 (0.09) | 0.51 (0.09) | 0.54 (0.06) | 0.48 (0.06) | 0.53 (0.04) | 0.48 (0.02) |
| DIABLO | 0.53 (0.04) | 0.53 (0.04) | 0.54 (0.06) | 0.50 (0.09) | 0.50 (0.04) | 0.46 (0.08) | 0.56 (0.05) | 0.57 (0.05) | 0.57 (0.06) | 0.56 (0.06) | 0.54 (0.03) | 0.56 (0.05) |
| GDF | 0.59 (0.08) | 0.59 (0.06) | 0.57 (0.06) | 0.56 (0.06) | 0.59 (0.08) | 0.56 (0.04) | 0.59 (0.08) | 0.59 (0.06) | 0.56 (0.05) | 0.58 (0.05) | 0.60 (0.07) | 0.57 (0.05) |
| Stabl | 0.54 (0.09) | 0.53 (0.10) | 0.56 (0.07) | 0.55 (0.07) | 0.51 (0.01) | 0.49 (0.03) | 0.57 (0.07) | 0.55 (0.07) | 0.57 (0.08) | 0.56 (0.06) | 0.57 (0.08) | 0.55 (0.09) |
| asmbPLS-DA | 0.58 (0.06) | 0.60 (0.05) | 0.58 (0.07) | 0.59 (0.06) | 0.55 (0.07) | 0.55 (0.05) | 0.57 (0.07) | 0.59 (0.07) | 0.56 (0.08) | 0.56 (0.07) | 0.55 (0.09) | 0.55 (0.07) |

**Supplementary Table 12:** Prediction performance for the COADREAD survival task. Mean and standard deviation across five-fold are reported. The following methods are excluded due to a lack of sample-level label prediction module: MOFA, MCIA, GAUDI, and DPM.

| Model | CNV+DNAm+mRNA | | CNV+SNV+mRNA | | CNV+mRNA+miRNA | | DNAm+SNV+mRNA | | DNAm+mRNA+miRNA | | SNV+mRNA+miRNA | |
|---|---|---|---|---|---|---|---|---|---|---|---|---|
| | AUPR | AUROC | AUPR | AUROC | AUPR | AUROC | AUPR | AUROC | AUPR | AUROC | AUPR | AUROC |
| DeePathNet | 0.59 (0.08) | 0.55 (0.07) | 0.56 (0.11) | 0.53 (0.10) | 0.59 (0.08) | 0.59 (0.08) | 0.62 (0.08) | 0.61 (0.07) | 0.61 (0.08) | 0.60 (0.08) | 0.61 (0.08) | 0.59 (0.07) |
| DeepKEGG | 0.57 (0.07) | 0.56 (0.06) | 0.63 (0.03) | 0.61 (0.05) | 0.60 (0.07) | 0.60 (0.10) | 0.56 (0.07) | 0.56 (0.08) | 0.52 (0.06) | 0.52 (0.04) | 0.58 (0.09) | 0.59 (0.11) |
| MOGLAM | 0.55 (0.08) | 0.54 (0.09) | 0.55 (0.04) | 0.49 (0.07) | 0.54 (0.05) | 0.51 (0.06) | 0.58 (0.05) | 0.52 (0.04) | 0.58 (0.05) | 0.56 (0.06) | 0.56 (0.07) | 0.54 (0.11) |
| TMO-Net | 0.55 (0.04) | 0.55 (0.05) | 0.55 (0.10) | 0.53 (0.10) | 0.57 (0.06) | 0.56 (0.06) | 0.57 (0.02) | 0.59 (0.03) | 0.54 (0.04) | 0.53 (0.06) | 0.49 (0.05) | 0.46 (0.06) |
| CustOmics | 0.57 (0.06) | 0.58 (0.05) | 0.52 (0.10) | 0.50 (0.10) | 0.55 (0.05) | 0.56 (0.08) | 0.52 (0.04) | 0.52 (0.06) | 0.56 (0.07) | 0.54 (0.06) | 0.54 (0.09) | 0.52 (0.12) |
| GENIUS | 0.54 (0.05) | 0.53 (0.04) | 0.55 (0.07) | 0.56 (0.06) | 0.53 (0.03) | 0.53 (0.03) | 0.52 (0.07) | 0.50 (0.07) | 0.51 (0.04) | 0.50 (0.05) | 0.53 (0.02) | 0.53 (0.03) |
| Pathformer | 0.51 (0.06) | 0.50 (0.05) | 0.58 (0.08) | 0.55 (0.06) | 0.59 (0.07) | 0.56 (0.07) | 0.52 (0.04) | 0.48 (0.06) | 0.58 (0.09) | 0.56 (0.09) | 0.54 (0.04) | 0.49 (0.04) |
| GNN-SubNet | 0.56 (0.12) | 0.53 (0.10) | 0.59 (0.04) | 0.57 (0.05) | 0.61 (0.08) | 0.58 (0.09) | 0.59 (0.11) | 0.57 (0.11) | 0.52 (0.05) | 0.51 (0.06) | 0.51 (0.08) | 0.47 (0.08) |
| P-Net | 0.58 (0.06) | 0.58 (0.08) | 0.60 (0.07) | 0.60 (0.07) | 0.59 (0.07) | 0.58 (0.09) | 0.59 (0.02) | 0.60 (0.03) | 0.60 (0.06) | 0.58 (0.06) | 0.59 (0.05) | 0.57 (0.06) |
| MOGONET | 0.55 (0.03) | 0.53 (0.04) | 0.49 (0.05) | 0.46 (0.07) | 0.51 (0.03) | 0.49 (0.07) | 0.54 (0.05) | 0.52 (0.05) | 0.48 (0.02) | 0.45 (0.04) | 0.50 (0.05) | 0.44 (0.02) |
| MORE | 0.58 (0.09) | 0.56 (0.12) | 0.54 (0.06) | 0.50 (0.08) | 0.52 (0.02) | 0.50 (0.02) | 0.59 (0.05) | 0.57 (0.04) | 0.52 (0.05) | 0.50 (0.09) | 0.56 (0.02) | 0.52 (0.07) |
| MoAGL-SA | 0.56 (0.06) | 0.55 (0.04) | 0.59 (0.03) | 0.57 (0.06) | 0.57 (0.05) | 0.56 (0.04) | 0.53 (0.04) | 0.52 (0.04) | 0.58 (0.02) | 0.57 (0.04) | 0.52 (0.03) | 0.51 (0.06) |
| DIABLO | 0.57 (0.07) | 0.54 (0.06) | 0.58 (0.05) | 0.56 (0.06) | 0.55 (0.07) | 0.52 (0.07) | 0.52 (0.04) | 0.49 (0.08) | 0.56 (0.07) | 0.55 (0.06) | 0.54 (0.06) | 0.52 (0.06) |
| GDF | 0.55 (0.03) | 0.56 (0.05) | 0.59 (0.04) | 0.59 (0.03) | 0.55 (0.03) | 0.57 (0.04) | 0.55 (0.04) | 0.58 (0.04) | 0.54 (0.05) | 0.54 (0.05) | 0.53 (0.04) | 0.56 (0.05) |
| Stabl | 0.51 (0.05) | 0.48 (0.06) | 0.51 (0.05) | 0.49 (0.11) | 0.50 (0.06) | 0.44 (0.05) | 0.55 (0.03) | 0.54 (0.01) | 0.53 (0.05) | 0.49 (0.07) | 0.51 (0.03) | 0.48 (0.06) |
| asmbPLS-DA | 0.54 (0.05) | 0.52 (0.06) | 0.56 (0.04) | 0.54 (0.00) | 0.54 (0.05) | 0.52 (0.07) | 0.59 (0.06) | 0.57 (0.05) | 0.53 (0.02) | 0.53 (0.03) | 0.54 (0.02) | 0.53 (0.03) |

**Supplementary Table 13:** Prediction performance for the Cisplatin drug response (BLCA) prediction task. Mean and standard deviation across five-fold are reported. The following methods are excluded due to a lack of sample-level label prediction module: MOFA, MCIA, GAUDI, and DPM.

| Model | CNV+DNAm+mRNA | | CNV+SNV+mRNA | | CNV+mRNA+miRNA | | DNAm+SNV+mRNA | | DNAm+mRNA+miRNA | | SNV+mRNA+miRNA | |
|---|---|---|---|---|---|---|---|---|---|---|---|---|
| | AUPR | AUROC | AUPR | AUROC | AUPR | AUROC | AUPR | AUROC | AUPR | AUROC | AUPR | AUROC |
| DeePathNet | 0.86 (0.15) | 0.72 (0.23) | 0.81 (0.06) | 0.64 (0.10) | 0.66 (0.10) | 0.41 (0.17) | 0.75 (0.11) | 0.53 (0.16) | 0.71 (0.13) | 0.52 (0.20) | 0.74 (0.12) | 0.51 (0.19) |
| DeepKEGG | 0.83 (0.05) | 0.61 (0.11) | 0.78 (0.05) | 0.54 (0.07) | 0.81 (0.08) | 0.62 (0.18) | 0.82 (0.05) | 0.61 (0.08) | 0.79 (0.11) | 0.53 (0.20) | 0.80 (0.13) | 0.57 (0.25) |
| MOGLAM | 0.78 (0.12) | 0.58 (0.18) | 0.70 (0.08) | 0.46 (0.11) | 0.72 (0.05) | 0.52 (0.07) | 0.76 (0.11) | 0.58 (0.16) | 0.73 (0.06) | 0.49 (0.12) | 0.74 (0.13) | 0.53 (0.18) |
| TMO-Net | 0.79 (0.14) | 0.60 (0.23) | 0.80 (0.14) | 0.64 (0.21) | 0.72 (0.12) | 0.51 (0.20) | 0.80 (0.10) | 0.57 (0.18) | 0.79 (0.09) | 0.59 (0.12) | 0.76 (0.06) | 0.56 (0.11) |
| CustOmics | 0.76 (0.08) | 0.56 (0.12) | 0.79 (0.12) | 0.61 (0.21) | 0.79 (0.04) | 0.61 (0.14) | 0.80 (0.05) | 0.57 (0.08) | 0.79 (0.06) | 0.62 (0.10) | 0.83 (0.05) | 0.64 (0.13) |
| GENIUS | 0.80 (0.11) | 0.62 (0.12) | 0.83 (0.11) | 0.68 (0.16) | 0.83 (0.08) | 0.65 (0.15) | 0.82 (0.08) | 0.63 (0.16) | 0.74 (0.04) | 0.52 (0.08) | 0.75 (0.13) | 0.57 (0.18) |
| Pathformer | 0.75 (0.09) | 0.54 (0.11) | 0.72 (0.18) | 0.49 (0.28) | 0.69 (0.10) | 0.41 (0.14) | 0.83 (0.06) | 0.63 (0.15) | 0.64 (0.12) | 0.36 (0.23) | 0.78 (0.07) | 0.61 (0.11) |
| GNN-SubNet | 0.72 (0.13) | 0.50 (0.26) | 0.78 (0.09) | 0.59 (0.13) | 0.76 (0.06) | 0.59 (0.09) | 0.73 (0.08) | 0.54 (0.10) | 0.68 (0.09) | 0.41 (0.15) | 0.75 (0.09) | 0.56 (0.12) |
| P-Net | 0.81 (0.06) | 0.62 (0.07) | 0.79 (0.08) | 0.61 (0.11) | 0.77 (0.08) | 0.54 (0.10) | 0.80 (0.02) | 0.57 (0.05) | 0.81 (0.02) | 0.60 (0.07) | 0.75 (0.07) | 0.54 (0.09) |
| MOGONET | 0.77 (0.05) | 0.54 (0.07) | 0.71 (0.09) | 0.51 (0.18) | 0.75 (0.07) | 0.56 (0.11) | 0.70 (0.07) | 0.47 (0.14) | 0.75 (0.07) | 0.55 (0.10) | 0.75 (0.19) | 0.53 (0.32) |
| MORE | 0.72 (0.12) | 0.54 (0.14) | 0.83 (0.12) | 0.62 (0.24) | 0.75 (0.11) | 0.53 (0.21) | 0.72 (0.12) | 0.52 (0.18) | 0.75 (0.07) | 0.56 (0.09) | 0.68 (0.07) | 0.46 (0.15) |
| MoAGL-SA | 0.72 (0.14) | 0.48 (0.24) | 0.76 (0.07) | 0.55 (0.08) | 0.78 (0.12) | 0.56 (0.25) | 0.76 (0.09) | 0.55 (0.16) | 0.77 (0.09) | 0.56 (0.09) | 0.64 (0.13) | 0.34 (0.25) |
| DIABLO | 0.74 (0.05) | 0.51 (0.12) | 0.79 (0.09) | 0.56 (0.15) | 0.70 (0.12) | 0.46 (0.15) | 0.76 (0.04) | 0.57 (0.08) | 0.79 (0.05) | 0.58 (0.09) | 0.70 (0.07) | 0.42 (0.11) |
| GDF | 0.84 (0.07) | 0.64 (0.15) | 0.84 (0.08) | 0.66 (0.18) | 0.87 (0.08) | 0.70 (0.18) | 0.74 (0.08) | 0.56 (0.09) | 0.79 (0.07) | 0.62 (0.10) | 0.77 (0.07) | 0.59 (0.15) |
| Stabl | 0.68 (0.08) | 0.41 (0.12) | 0.82 (0.06) | 0.65 (0.11) | 0.59 (0.04) | 0.31 (0.14) | 0.76 (0.07) | 0.59 (0.13) | 0.73 (0.07) | 0.54 (0.08) | 0.78 (0.17) | 0.59 (0.34) |
| asmbPLS-DA | 0.77 (0.10) | 0.56 (0.14) | 0.75 (0.13) | 0.53 (0.24) | 0.75 (0.12) | 0.56 (0.17) | 0.82 (0.09) | 0.60 (0.18) | 0.80 (0.09) | 0.57 (0.13) | 0.82 (0.09) | 0.61 (0.16) |

**Supplementary Table 14:** Prediction performance for the Temozolomide drug response (LGG) prediction task. Mean and standard deviation across five-fold are reported. The following methods are excluded due to a lack of sample-level label prediction module: MOFA, MCIA, GAUDI, and DPM.

| Model | CNV+DNAm+mRNA | | CNV+SNV+mRNA | | CNV+mRNA+miRNA | | DNAm+SNV+mRNA | | DNAm+mRNA+miRNA | | SNV+mRNA+miRNA | |
|---|---|---|---|---|---|---|---|---|---|---|---|---|
| | AUPR | AUROC | AUPR | AUROC | AUPR | AUROC | AUPR | AUROC | AUPR | AUROC | AUPR | AUROC |
| DeePathNet | 0.23 (0.13) | 0.44 (0.13) | 0.24 (0.11) | 0.46 (0.19) | 0.29 (0.23) | 0.55 (0.18) | 0.24 (0.05) | 0.55 (0.09) | 0.34 (0.13) | 0.58 (0.19) | 0.18 (0.05) | 0.35 (0.10) |
| DeepKEGG | 0.39 (0.19) | 0.70 (0.13) | 0.32 (0.11) | 0.56 (0.15) | 0.28 (0.10) | 0.53 (0.12) | 0.36 (0.22) | 0.58 (0.25) | 0.28 (0.17) | 0.57 (0.19) | 0.24 (0.10) | 0.53 (0.13) |
| MOGLAM | 0.44 (0.19) | 0.64 (0.17) | 0.28 (0.17) | 0.58 (0.19) | 0.22 (0.05) | 0.50 (0.11) | 0.24 (0.07) | 0.56 (0.13) | 0.33 (0.08) | 0.62 (0.04) | 0.24 (0.11) | 0.50 (0.14) |
| TMO-Net | 0.23 (0.09) | 0.55 (0.18) | 0.34 (0.14) | 0.65 (0.17) | 0.29 (0.08) | 0.55 (0.13) | 0.27 (0.15) | 0.50 (0.25) | 0.27 (0.14) | 0.57 (0.19) | 0.33 (0.28) | 0.57 (0.24) |
| CustOmics | 0.29 (0.15) | 0.52 (0.18) | 0.32 (0.13) | 0.58 (0.18) | 0.38 (0.14) | 0.65 (0.15) | 0.27 (0.10) | 0.57 (0.14) | 0.38 (0.13) | 0.66 (0.15) | 0.26 (0.10) | 0.51 (0.11) |
| GENIUS | 0.37 (0.14) | 0.65 (0.21) | 0.37 (0.19) | 0.59 (0.16) | 0.38 (0.11) | 0.65 (0.13) | 0.34 (0.19) | 0.64 (0.21) | 0.33 (0.08) | 0.61 (0.18) | 0.31 (0.07) | 0.56 (0.12) |
| Pathformer | 0.37 (0.15) | 0.71 (0.10) | 0.32 (0.08) | 0.63 (0.12) | 0.29 (0.10) | 0.59 (0.10) | 0.30 (0.13) | 0.65 (0.11) | 0.27 (0.20) | 0.51 (0.23) | 0.24 (0.14) | 0.43 (0.21) |
| GNN-SubNet | 0.24 (0.08) | 0.54 (0.18) | 0.23 (0.05) | 0.52 (0.14) | 0.30 (0.07) | 0.63 (0.04) | 0.26 (0.10) | 0.47 (0.13) | 0.18 (0.03) | 0.43 (0.12) | 0.25 (0.14) | 0.53 (0.21) |
| P-Net | 0.30 (0.11) | 0.62 (0.16) | 0.31 (0.13) | 0.61 (0.18) | 0.33 (0.13) | 0.62 (0.18) | 0.30 (0.13) | 0.63 (0.18) | 0.29 (0.11) | 0.61 (0.16) | 0.28 (0.11) | 0.59 (0.15) |
| MOGONET | 0.28 (0.15) | 0.57 (0.25) | 0.32 (0.19) | 0.57 (0.13) | 0.28 (0.10) | 0.57 (0.12) | 0.37 (0.25) | 0.55 (0.23) | 0.33 (0.15) | 0.58 (0.14) | 0.32 (0.17) | 0.53 (0.15) |
| MORE | 0.34 (0.09) | 0.63 (0.15) | 0.23 (0.11) | 0.51 (0.21) | 0.23 (0.11) | 0.45 (0.18) | 0.27 (0.12) | 0.55 (0.16) | 0.28 (0.08) | 0.58 (0.16) | 0.25 (0.14) | 0.49 (0.23) |
| MoAGL-SA | 0.20 (0.07) | 0.40 (0.13) | 0.40 (0.12) | 0.67 (0.14) | 0.40 (0.22) | 0.58 (0.17) | 0.21 (0.13) | 0.37 (0.16) | 0.35 (0.17) | 0.58 (0.17) | 0.24 (0.09) | 0.48 (0.09) |
| DIABLO | 0.27 (0.07) | 0.63 (0.11) | 0.30 (0.10) | 0.62 (0.17) | 0.26 (0.06) | 0.61 (0.11) | 0.25 (0.11) | 0.52 (0.14) | 0.30 (0.11) | 0.61 (0.19) | 0.39 (0.14) | 0.57 (0.10) |
| GDF | 0.24 (0.09) | 0.60 (0.11) | 0.22 (0.10) | 0.53 (0.17) | 0.23 (0.15) | 0.53 (0.15) | 0.21 (0.10) | 0.51 (0.14) | 0.28 (0.13) | 0.57 (0.16) | 0.17 (0.05) | 0.48 (0.16) |
| Stabl | 0.30 (0.11) | 0.57 (0.07) | 0.31 (0.14) | 0.54 (0.18) | 0.21 (0.07) | 0.53 (0.16) | 0.21 (0.06) | 0.48 (0.11) | 0.27 (0.10) | 0.56 (0.14) | 0.17 (0.03) | 0.39 (0.09) |
| asmbPLS-DA | 0.26 (0.08) | 0.59 (0.14) | 0.29 (0.20) | 0.52 (0.17) | 0.32 (0.13) | 0.60 (0.17) | 0.26 (0.08) | 0.56 (0.16) | 0.27 (0.10) | 0.54 (0.10) | 0.26 (0.11) | 0.51 (0.15) |

**Supplementary Table 15:** InterSIM prediction results. Mean and standard deviation values across five-fold are reported. The following methods are excluded due to a lack of sample-level label prediction module: MOFA, MCIA, GAUDI, and DPM.

| Model | $\delta = 0.5$ | | $\delta = 1.0$ | | $\delta = 2.0$ | | $\delta = 3.0$ | | $\delta = 4.0$ | | $\delta = 5.0$ | |
|---|---|---|---|---|---|---|---|---|---|---|---|---|
| | AUPR | AUROC | AUPR | AUROC | AUPR | AUROC | AUPR | AUROC | AUPR | AUROC | AUPR | AUROC |
| DeePathNet | 0.58 | 0.49 | 0.52 | 0.43 | 0.72 | 0.71 | 0.57 | 0.50 | 0.65 | 0.55 | 0.65 | 0.62 |
| | (0.16) | (0.20) | (0.10) | (0.21) | (0.13) | (0.14) | (0.08) | (0.05) | (0.14) | (0.20) | (0.12) | (0.13) |
| DeepKEGG | 0.71 | 0.74 | 0.85 | 0.83 | 0.95 | 0.94 | 1.00 | 1.00 | 0.97 | 0.96 | 1.00 | 1.00 |
| | (0.29) | (0.26) | (0.21) | (0.24) | (0.09) | (0.13) | (0.00) | (0.00) | (0.07) | (0.07) | (0.00) | (0.00) |
| MOGLAM | 0.98 | 0.98 | 1.00 | 1.00 | 1.00 | 1.00 | 1.00 | 1.00 | 1.00 | 1.00 | 1.00 | 1.00 |
| | (0.02) | (0.02) | (0.00) | (0.00) | (0.00) | (0.00) | (0.00) | (0.00) | (0.00) | (0.00) | (0.00) | (0.00) |
| TMO-Net | 0.95 | 0.92 | 1.00 | 1.00 | 1.00 | 1.00 | 1.00 | 1.00 | 1.00 | 1.00 | 1.00 | 1.00 |
| | (0.04) | (0.11) | (0.00) | (0.00) | (0.00) | (0.00) | (0.00) | (0.00) | (0.00) | (0.00) | (0.00) | (0.00) |
| CustOmics | 0.99 | 0.99 | 1.00 | 1.00 | 1.00 | 1.00 | 1.00 | 1.00 | 1.00 | 1.00 | 1.00 | 1.00 |
| | (0.01) | (0.01) | (0.00) | (0.00) | (0.00) | (0.00) | (0.00) | (0.00) | (0.00) | (0.00) | (0.00) | (0.00) |
| GENIUS | 0.96 | 0.97 | 1.00 | 1.00 | 1.00 | 1.00 | 1.00 | 1.00 | 1.00 | 1.00 | 1.00 | 1.00 |
| | (0.04) | (0.03) | (0.00) | (0.00) | (0.00) | (0.00) | (0.00) | (0.00) | (0.00) | (0.00) | (0.00) | (0.00) |
| Pathformer | 0.90 | 0.90 | 0.99 | 0.99 | 1.00 | 1.00 | 1.00 | 1.00 | 1.00 | 1.00 | 1.00 | 1.00 |
| | (0.10) | (0.09) | (0.01) | (0.01) | (0.00) | (0.00) | (0.00) | (0.00) | (0.01) | (0.01) | (0.00) | (0.00) |
| GNN-SubNet | 0.71 | 0.70 | 0.82 | 0.76 | 0.94 | 0.96 | 0.96 | 0.95 | 0.74 | 0.77 | 0.97 | 0.97 |
| | (0.09) | (0.08) | (0.04) | (0.08) | (0.08) | (0.04) | (0.08) | (0.09) | (0.09) | (0.09) | (0.06) | (0.05) |
| P-Net | 0.83 | 0.79 | 0.97 | 0.97 | 1.00 | 1.00 | 1.00 | 1.00 | 1.00 | 1.00 | 1.00 | 1.00 |
| | (0.11) | (0.13) | (0.01) | (0.01) | (0.00) | (0.00) | (0.00) | (0.00) | (0.00) | (0.00) | (0.00) | (0.00) |
| MOGONET | 0.92 | 0.92 | 0.98 | 0.97 | 0.99 | 0.98 | 1.00 | 1.00 | 0.99 | 0.99 | 1.00 | 1.00 |
| | (0.11) | (0.12) | (0.02) | (0.03) | (0.01) | (0.02) | (0.00) | (0.00) | (0.01) | (0.02) | (0.00) | (0.00) |
| MORE | 0.81 | 0.78 | 0.86 | 0.85 | 0.97 | 0.97 | 0.98 | 0.98 | 0.98 | 0.98 | 0.99 | 0.99 |
| | (0.14) | (0.16) | (0.20) | (0.19) | (0.04) | (0.04) | (0.02) | (0.02) | (0.02) | (0.02) | (0.02) | (0.02) |
| MoAGL-SA | 0.70 | 0.68 | 0.98 | 0.97 | 0.95 | 0.94 | 0.99 | 0.99 | 1.00 | 0.99 | 1.00 | 0.99 |
| | (0.25) | (0.25) | (0.03) | (0.04) | (0.05) | (0.07) | (0.01) | (0.01) | (0.01) | (0.01) | (0.01) | (0.01) |
| DIABLO | 0.96 | 0.91 | 1.00 | 1.00 | 1.00 | 1.00 | 1.00 | 1.00 | 1.00 | 1.00 | 1.00 | 1.00 |
| | (0.07) | (0.17) | (0.01) | (0.01) | (0.00) | (0.00) | (0.00) | (0.00) | (0.00) | (0.00) | (0.00) | (0.00) |
| GDF | 1.00 | 1.00 | 1.00 | 1.00 | 1.00 | 1.00 | 1.00 | 1.00 | 1.00 | 1.00 | 1.00 | 1.00 |
| | (0.00) | (0.00) | (0.00) | (0.00) | (0.00) | (0.00) | (0.00) | (0.00) | (0.00) | (0.00) | (0.00) | (0.00) |
| Stabl | 1.00 | 1.00 | 1.00 | 1.00 | 1.00 | 1.00 | 1.00 | 1.00 | 1.00 | 1.00 | 1.00 | 1.00 |
| | (0.00) | (0.00) | (0.00) | (0.00) | (0.00) | (0.00) | (0.00) | (0.00) | (0.00) | (0.00) | (0.00) | (0.00) |
| asmbPLS-DA | 1.00 | 1.00 | 1.00 | 1.00 | 1.00 | 1.00 | 1.00 | 1.00 | 1.00 | 1.00 | 1.00 | 1.00 |
| | (0.00) | (0.00) | (0.00) | (0.00) | (0.00) | (0.00) | (0.00) | (0.00) | (0.00) | (0.00) | (0.00) | (0.00) |

[27] Zhitao Ying, Dylan Bourgeois, Jiaxuan You, Marinka Zitnik, and Jure Leskovec. GNNExplainer: Generating Explanations for Graph Neural Networks. In *Advances in Neural Information Processing Systems*, volume 32, 2019.

[36] Julien Hédou, Ivana Marić, Grégoire Bellan, Jakob Einhaus, Dyani K. Gaudillière, Francois-Xavier Ladant, Franck Verdonk, Ina A. Stelzer, Dorien Feyaerts, Amy S. Tsai, Edward A. Ganio, Maximilian Sabayev, Joshua Gillard, Jonas Amar, Amelie Cambriel, Tomiko T. Oskotsky, Alennie Roldan, Jonathan L. Golob, Marina Sirota, Thomas A. Bonham, Masaki Sato, Maïgane Diop, Xavier Durand,

Martin S. Angst, David K. Stevenson, Nima Aghaeepour, Andrea Montanari, and Brice Gaudillière. Discovery of sparse, reliable omic biomarkers with Stabl. *Nature Biotechnology*, 42(10):1581–1593, October 2024. ISSN 1546-1696. 10.1038/s41587-023-02033-x. URL https://www.nature.com/articles/s41587-023-02033-x. Publisher: Nature Publishing Group.

[37] Emmanuel Candès, Yingying Fan, Lucas Janson, and Jinchi Lv. Panning for Gold: 'Model-X' Knockoffs for High Dimensional Controlled Variable Selection. *Journal of the Royal Statistical Society Series B: Statistical Methodology*, 80(3):551–577, June 2018. ISSN 1369-7412. 10.1111/rssb.12265. URL https://doi.org/10.1111/rssb.12265.

[38] Bastian Pfeifer, Hubert Baniecki, Anna Saranti, Przemyslaw Biecek, and Andreas Holzinger. Multi-omics disease module detection with an explainable Greedy Decision Forest. *Scientific Reports*, 12(1): 16857, October 2022. ISSN 2045-2322. 10.1038/s41598-022-21417-8. URL https://www.nature.com/articles/s41598-022-21417-8. Publisher: Nature Publishing Group.

[39] Mykhaylo Slobodyanyuk, Alexander T. Bahcheli, Zoe P. Klein, Masroor Bayati, Lisa J. Strug, and Jüri Reimand. Directional integration and pathway enrichment analysis for multi-omics data. *Nature Communications*, 15(1):5690, July 2024. ISSN 2041-1723. 10.1038/s41467-024-49986-4. URL https://www.nature.com/articles/s41467-024-49986-4. Publisher: Nature Publishing Group.

[40] Marilyn M. Li, Michael Datto, Eric J. Duncavage, Shashikant Kulkarni, Neal I. Lindeman, Somak Roy, Apostolia M. Tsimberidou, Cindy L. Vnencak-Jones, Daynna J. Wolff, Anas Younes, and Marina N. Nikiforova. Standards and Guidelines for the Interpretation and Reporting of Sequence Variants in Cancer. *The Journal of Molecular Diagnostics*, 19(1):4–23, January 2017. ISSN 1525-1578. 10.1016/j.jmoldx.2016.10.002. URL https://linkinghub.elsevier.com/retrieve/pii/S1525157816302239. Publisher: Elsevier BV.
